## Supplemental File 4 for "OxPhos Dysfunction Causes Hypermetabolism and Reduces Lifespan in Cells and in Patients with Mitochondrial Diseases"

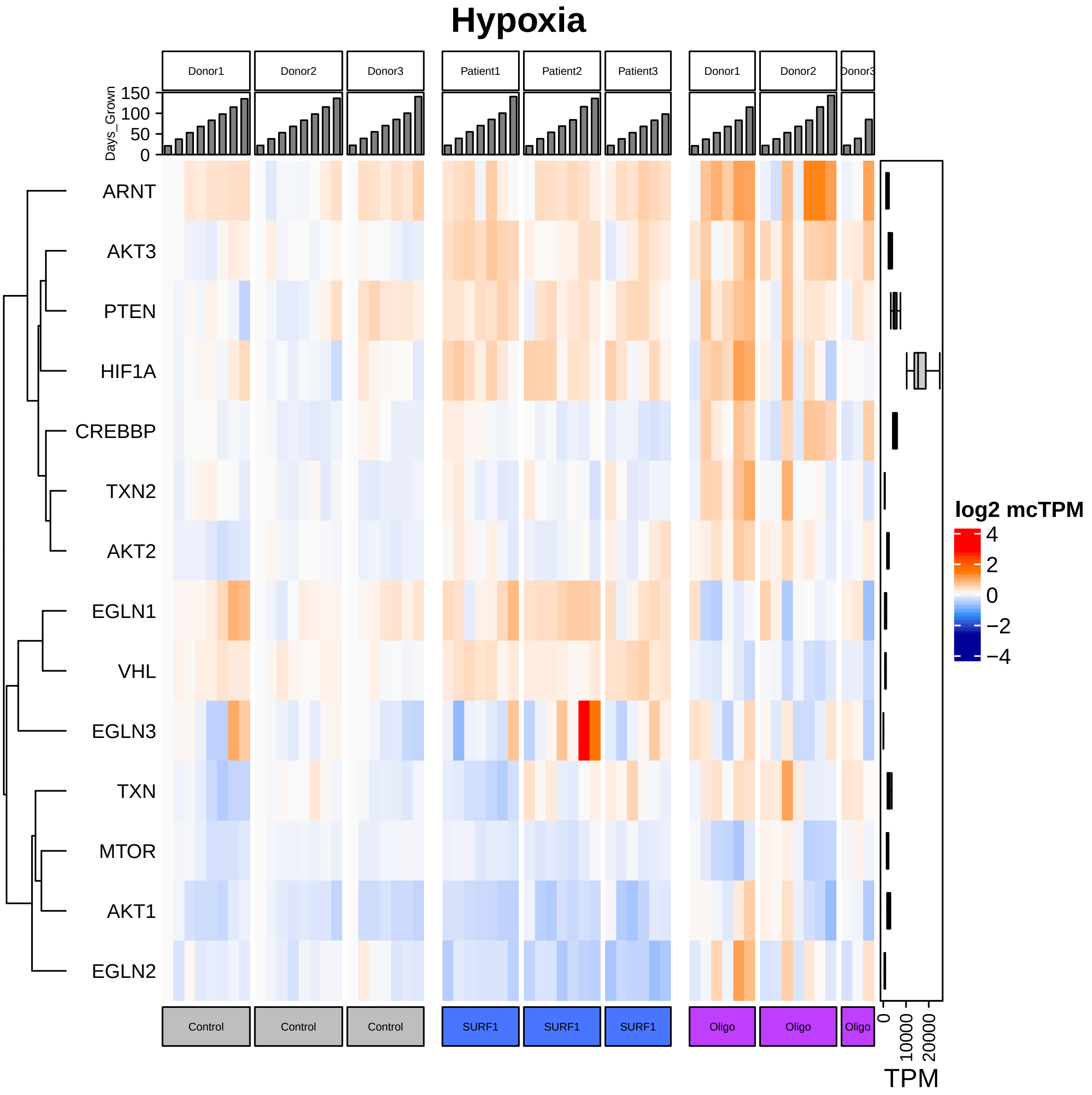
Hypoxia
Donor1
Donor2
Donor3
Patient1
Patient2
Patient3
Donor1
Donor2
Donor3
Days_Grown
150
100
50
0
ARNT
AKT3
PTEN
HIF1A
CREBBP
TXN2
AKT2
EGLN1
VHL
EGLN3
TXN
MTOR
AKT1
EGLN2
log2 mcTPM
4
2
0
−2
−4
Control
Control
Control
SURF1
SURF1
SURF1
Oligo
Oligo
Oligo
0
10000
20000
TPM

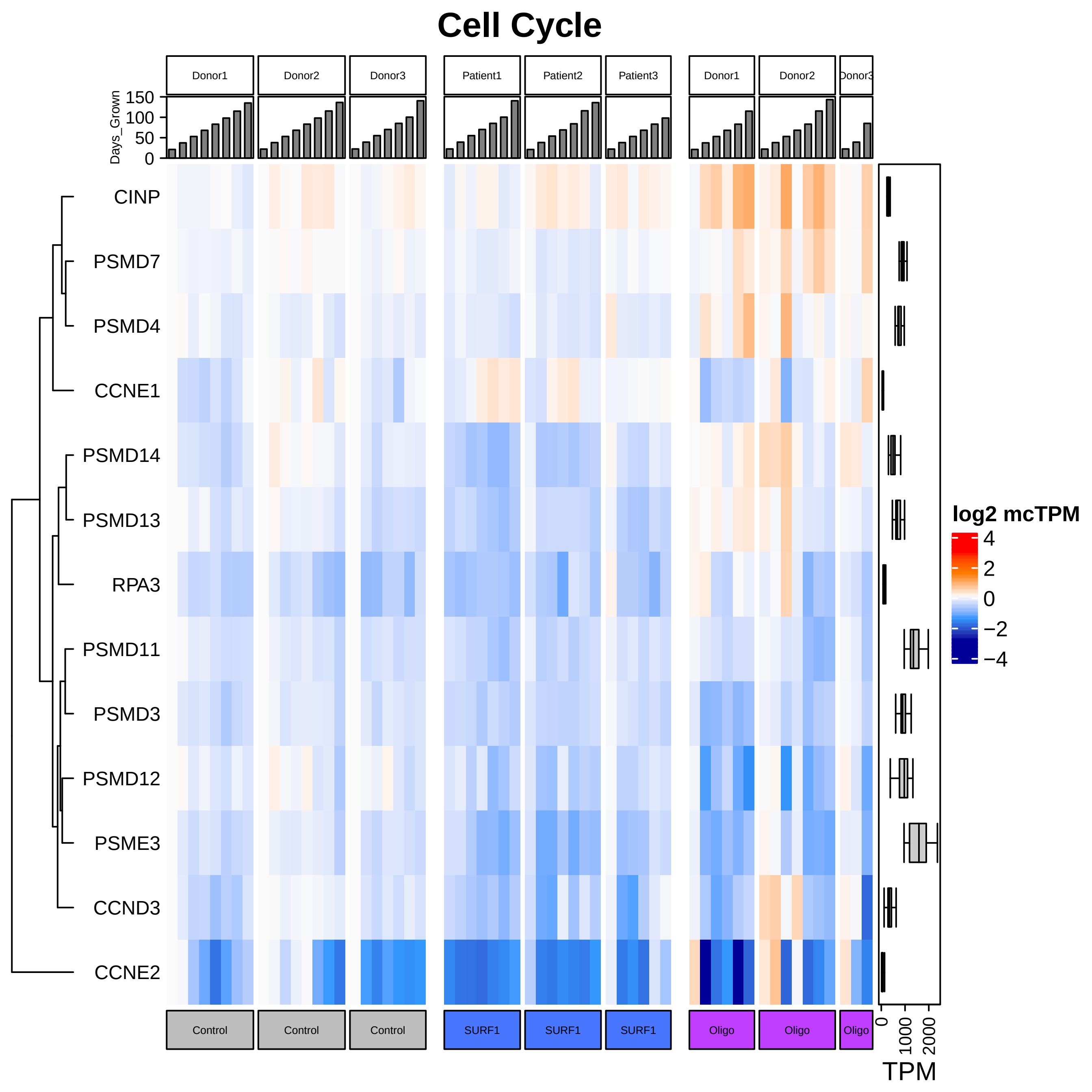
Cell Cycle
Donor1
Donor2
Donor3
Patient1
Patient2
Patient3
Donor1
Donor2
Donor3
Days_Grown
150
100
50
0
CINP
PSMD7
PSMD4
CCNE1
PSMD14
PSMD13
RPA3
PSMD11
PSMD3
PSMD12
PSME3
CCND3
CCNE2
log2 mcTPM
4
2
0
−2
−4
Control
Control
Control
SURF1
SURF1
SURF1
Oligo
Oligo
Oligo
0
1000
2000
TPM

### DNA Replication

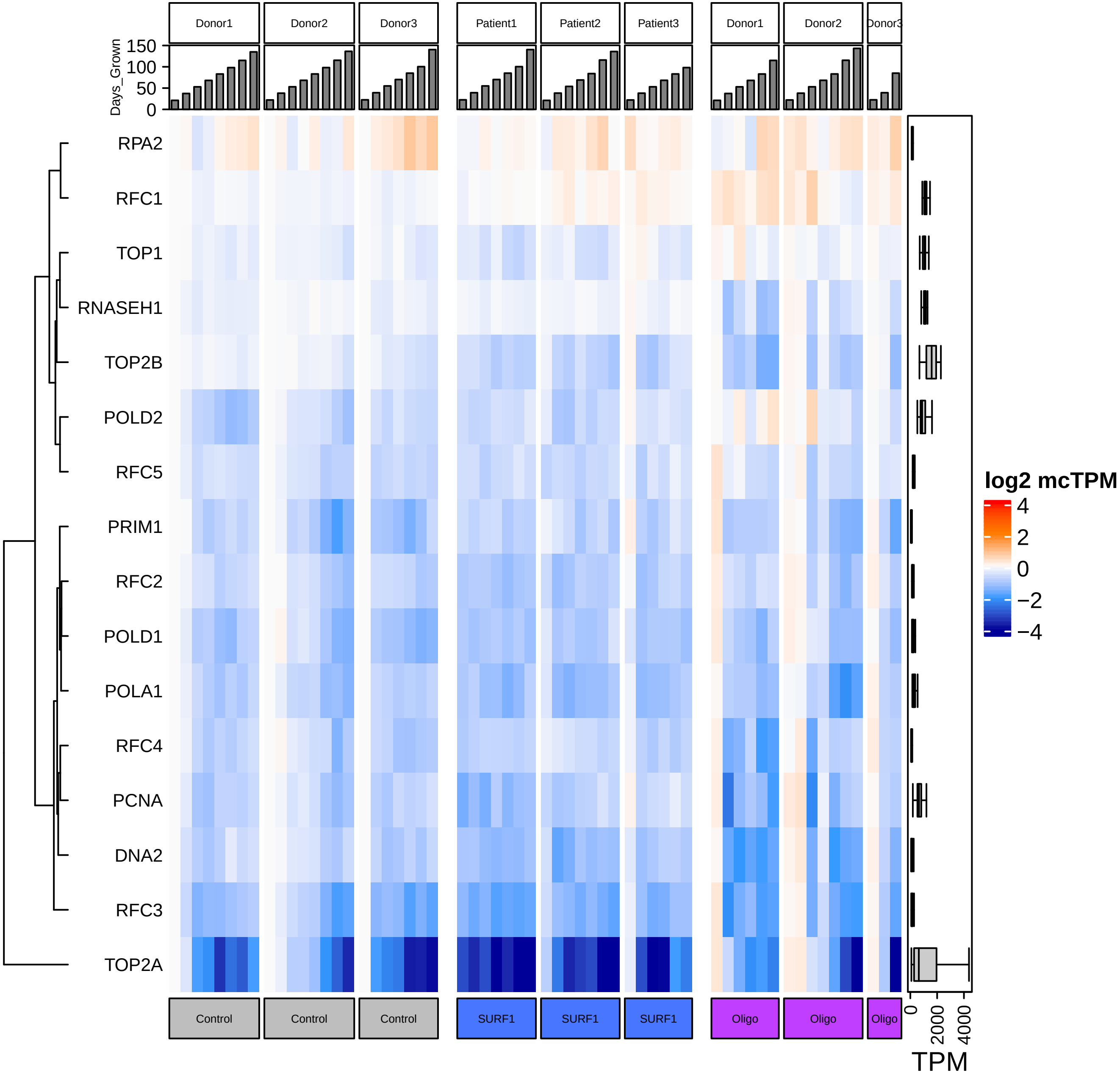

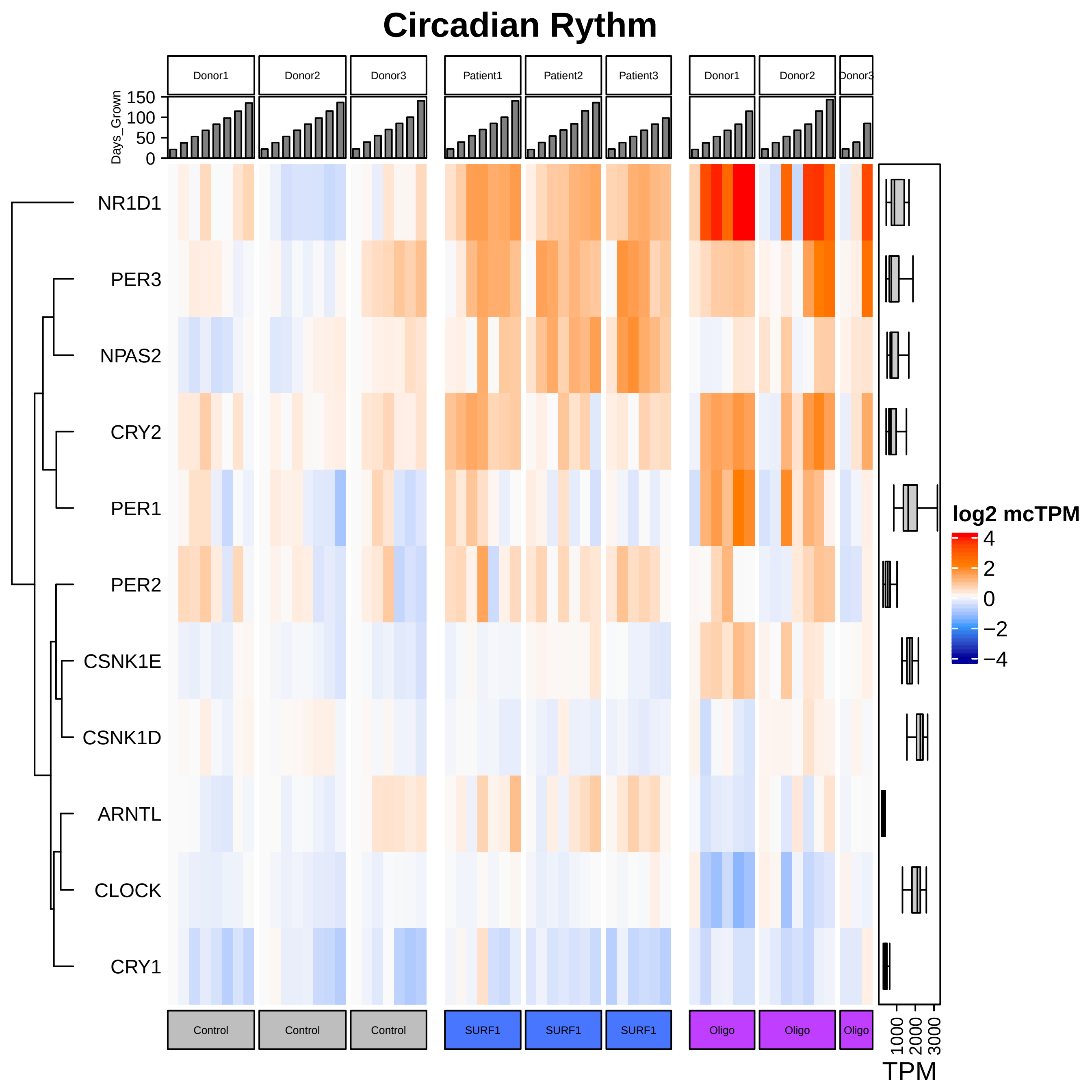
Circadian Rythm
Donor1
Donor2
Donor3
Patient1
Patient2
Patient3
Donor1
Donor2
Donor3
Days_Grown
150
100
50
0
NR1D1
PER3
NPAS2
CRY2
PER1
PER2
CSNK1E
CSNK1D
ARNTL
CLOCK
CRY1
log2 mcTPM
4
2
0
−2
−4
Control
Control
Control
SURF1
SURF1
SURF1
Oligo
Oligo
Oligo
1000
2000
3000
TPM

### Heme_Biosynthesis

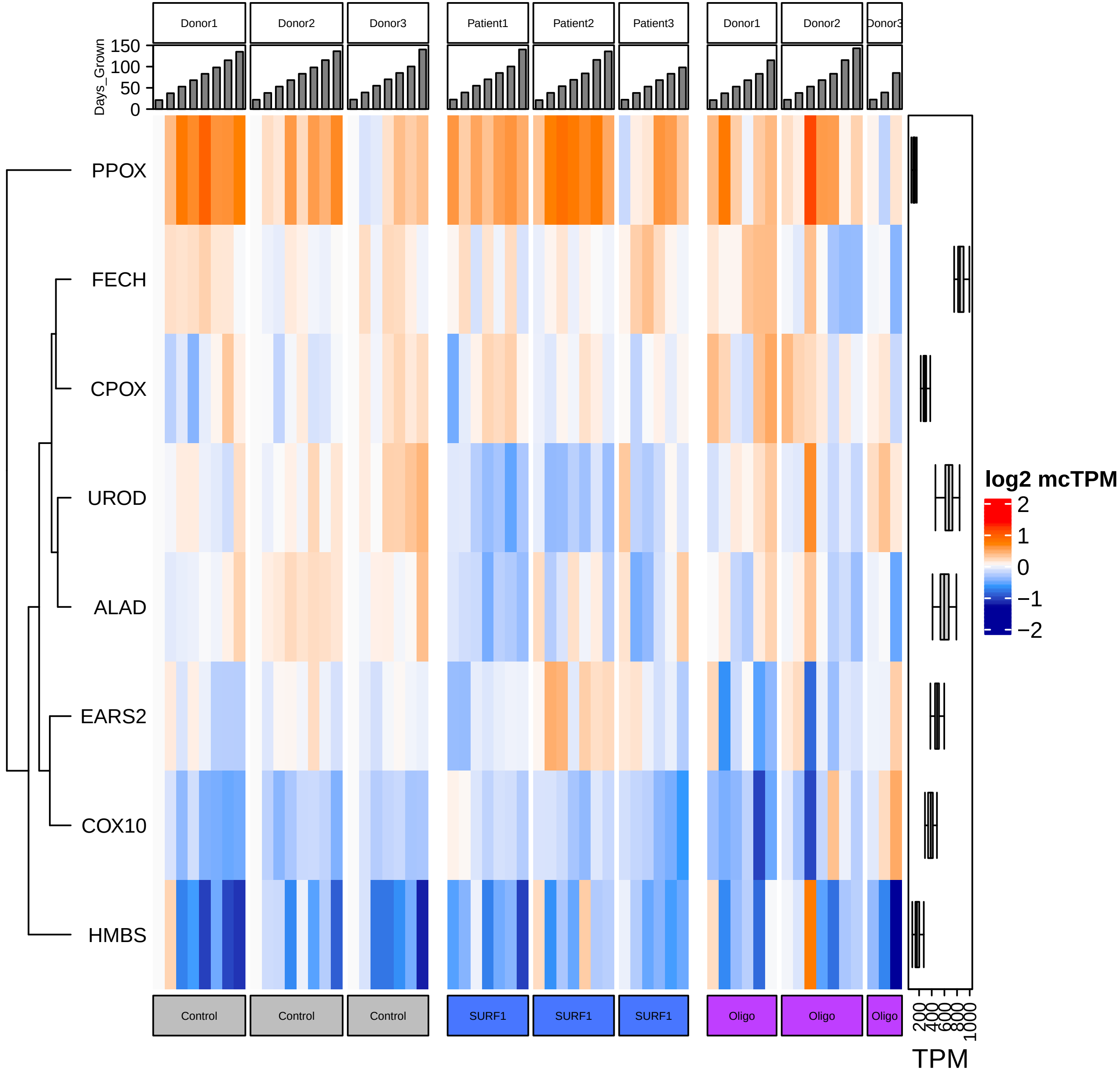

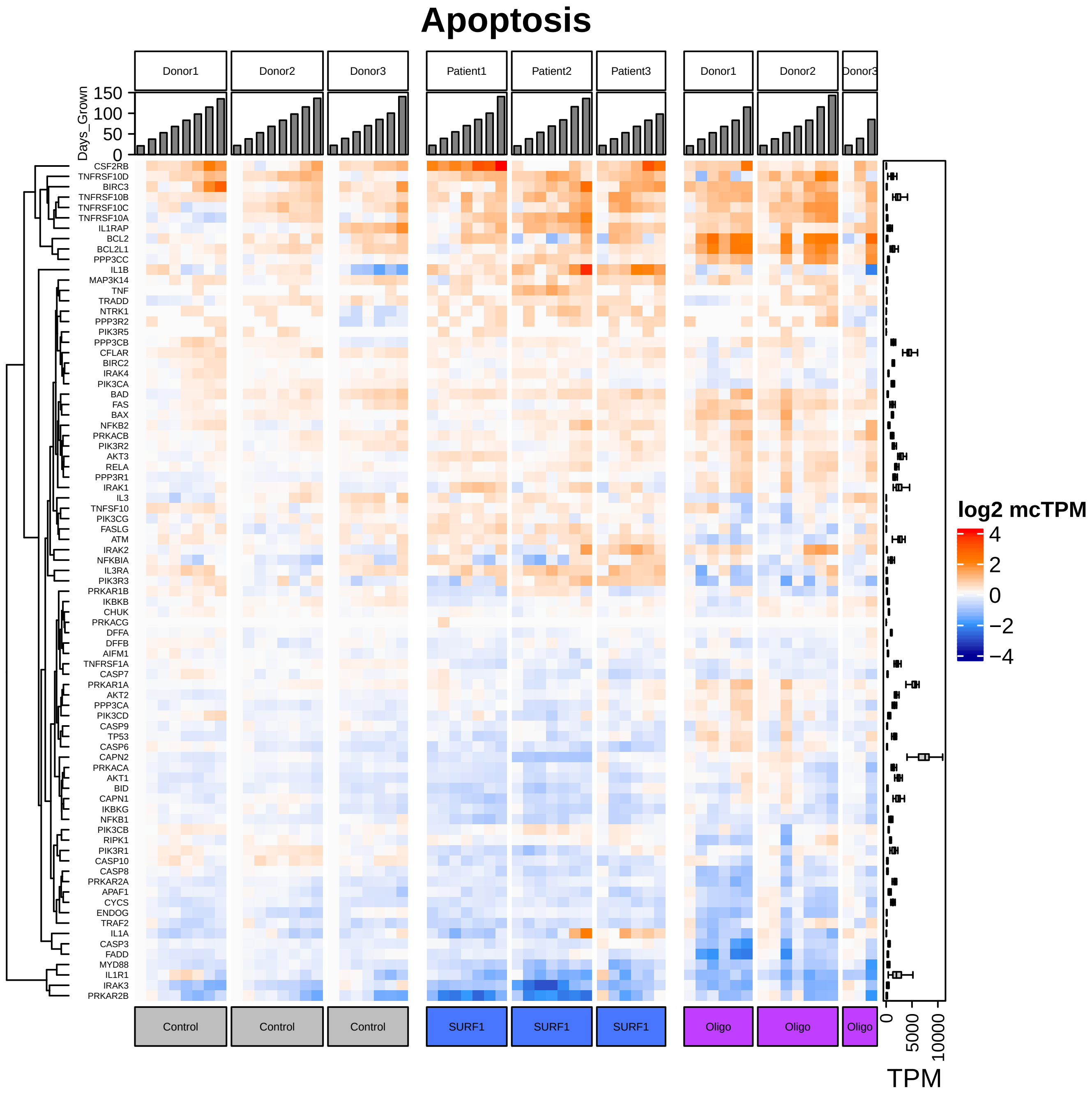
Apoptosis
Donor1
Donor2
Donor3
Patient1
Patient2
Patient3
Donor1
Donor2
Donor3
Days_Grown
150
100
50
0
CSF2RB
TNFRSF10D
BIRC3
TNFRSF10B
TNFRSF10C
TNFRSF10A
IL1RAP
BCL2
BCL2L1
PPP3CC
IL1B
MAP3K14
TNF
TRADD
NTRK1
PPP3R2
PIK3R5
PPP3CB
CFLAR
BIRC2
IRAK4
PIK3CA
BAD
FAS
BAX
NFKB2
PRKACB
PIK3R2
AKT3
RELA
PPP3R1
IRAK1
IL3
TNFSF10
PIK3CG
FASLG
ATM
IRAK2
NFKBIA
IL3RA
PIK3R3
PRKAR1B
IKBKB
CHUK
PRKACG
DFFA
DFFB
AIFM1
TNFRSF1A
CASP7
PRKAR1A
AKT2
PPP3CA
PIK3CD
CASP9
TP53
CASP6
CAPN2
PRKACA
AKT1
BID
CAPN1
IKBKG
NFKB1
PIK3CB
RIPK1
PIK3R1
CASP10
CASP8
PRKAR2A
APAF1
CYCS
ENDOG
TRAF2
IL1A
CASP3
FADD
MYD88
IL1R1
IRAK3
PRKAR2B
Control
Control
Control
SURF1
SURF1
SURF1
Oligo
Oligo
Oligo
log2 mcTPM
4
2
0
−2
−4
0
5000
10000
TPM

### Positive_Apoptosis

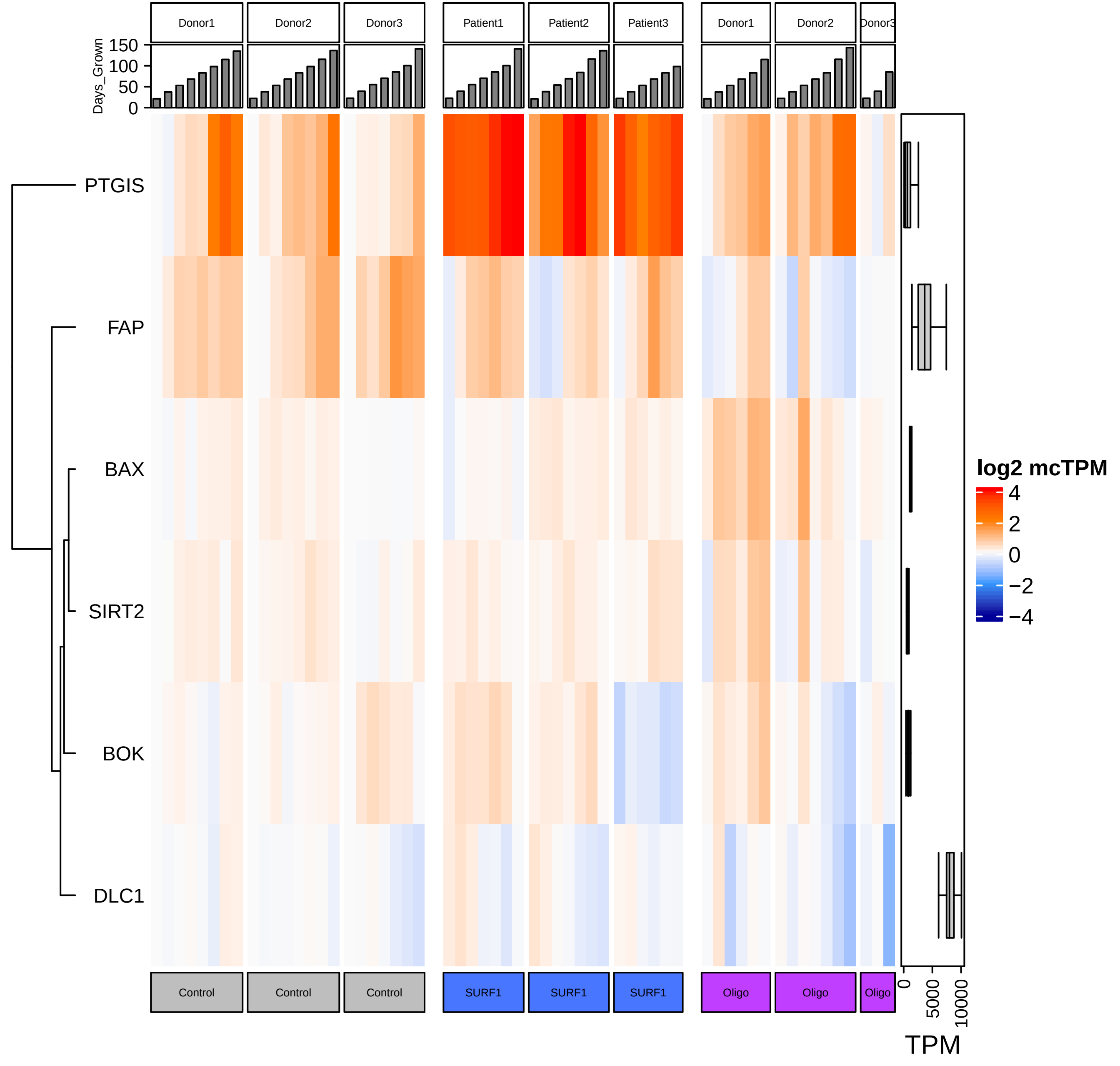

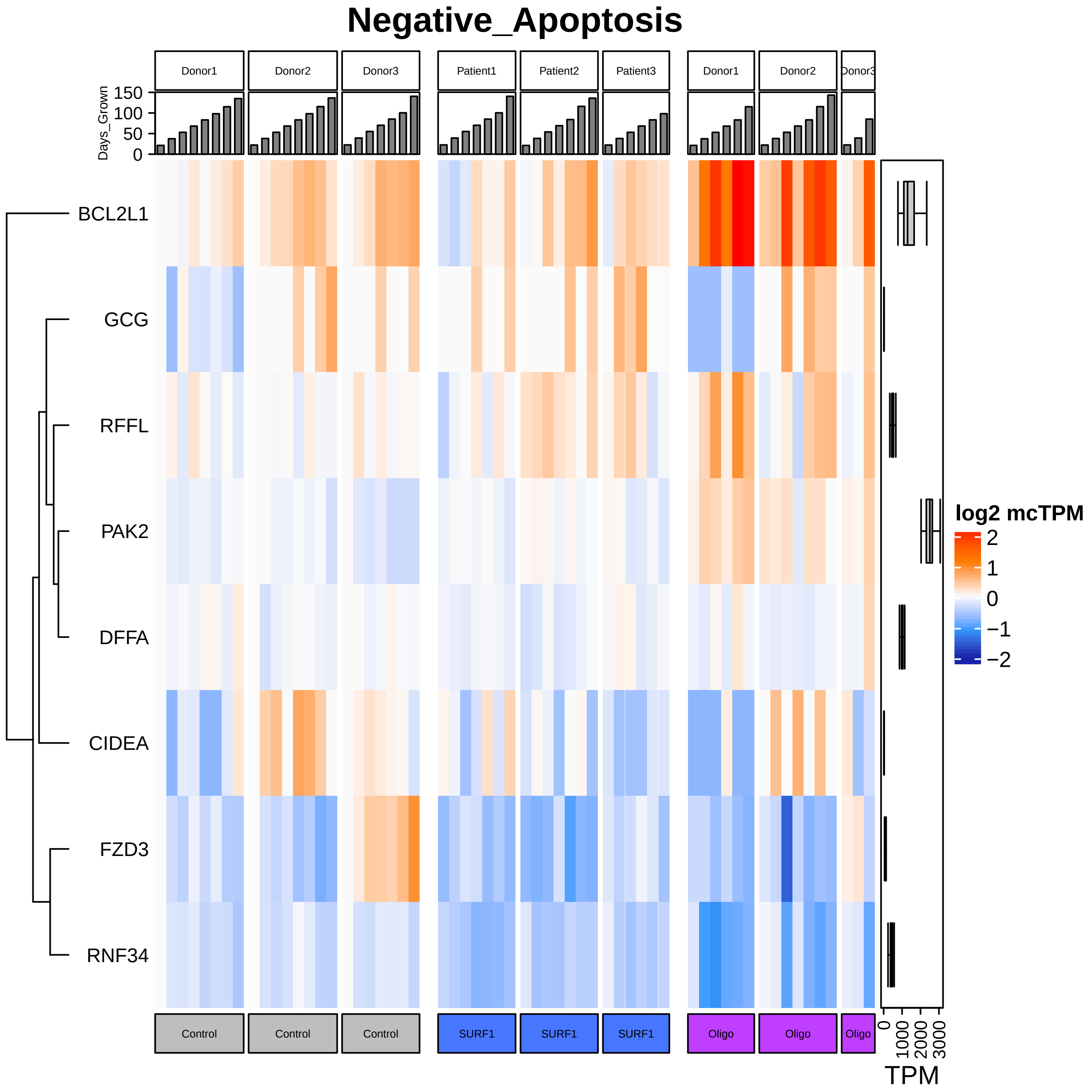

Negative_Apoptosis
Donor1
Donor2
Donor3
Patient1
Patient2
Patient3
Donor1
Donor2
Donor3
Days_Grown
150
100
50
0
BCL2L1
GCG
RFFL
PAK2
DFFA
CIDEA
FZD3
RNF34
log2 mcTPM
2
1
0
−1
−2
Control
Control
Control
SURF1
SURF1
SURF1
Oligo
Oligo
Oligo
0
1000
2000
3000
TPM

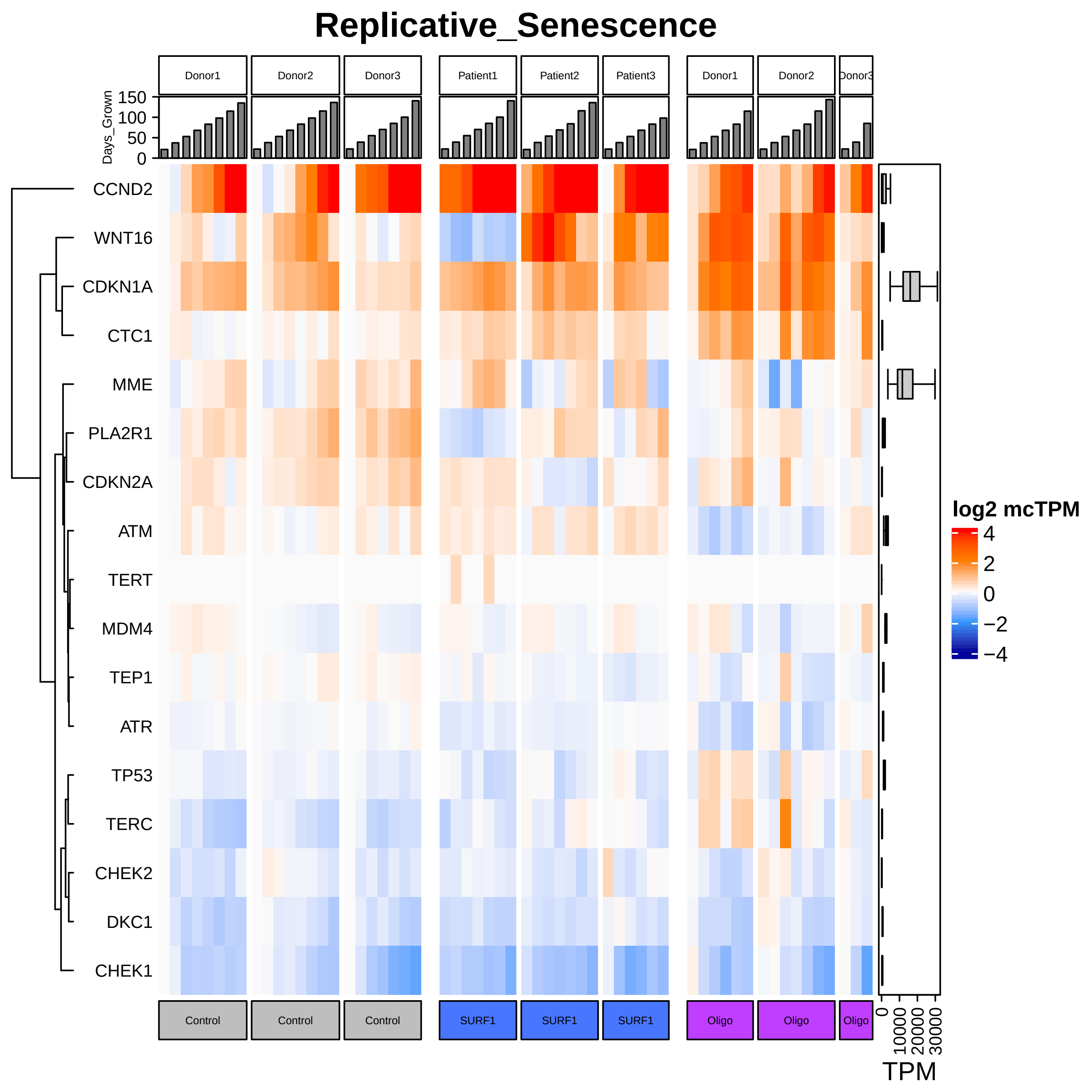

Replicative_Senescence
Donor1
Donor2
Donor3
Patient1
Patient2
Patient3
Donor1
Donor2
Donor3
Days_Grown
150
100
50
0
CCND2
WNT16
CDKN1A
CTC1
MME
PLA2R1
CDKN2A
ATM
TERT
MDM4
TEP1
ATR
TP53
TERC
CHEK2
DKC1
CHEK1
log2 mcTPM
4
2
0
−2
−4
Control
Control
Control
SURF1
SURF1
SURF1
Oligo
Oligo
Oligo
0
10000
20000
30000
TPM

### Regulation_of_Senescence

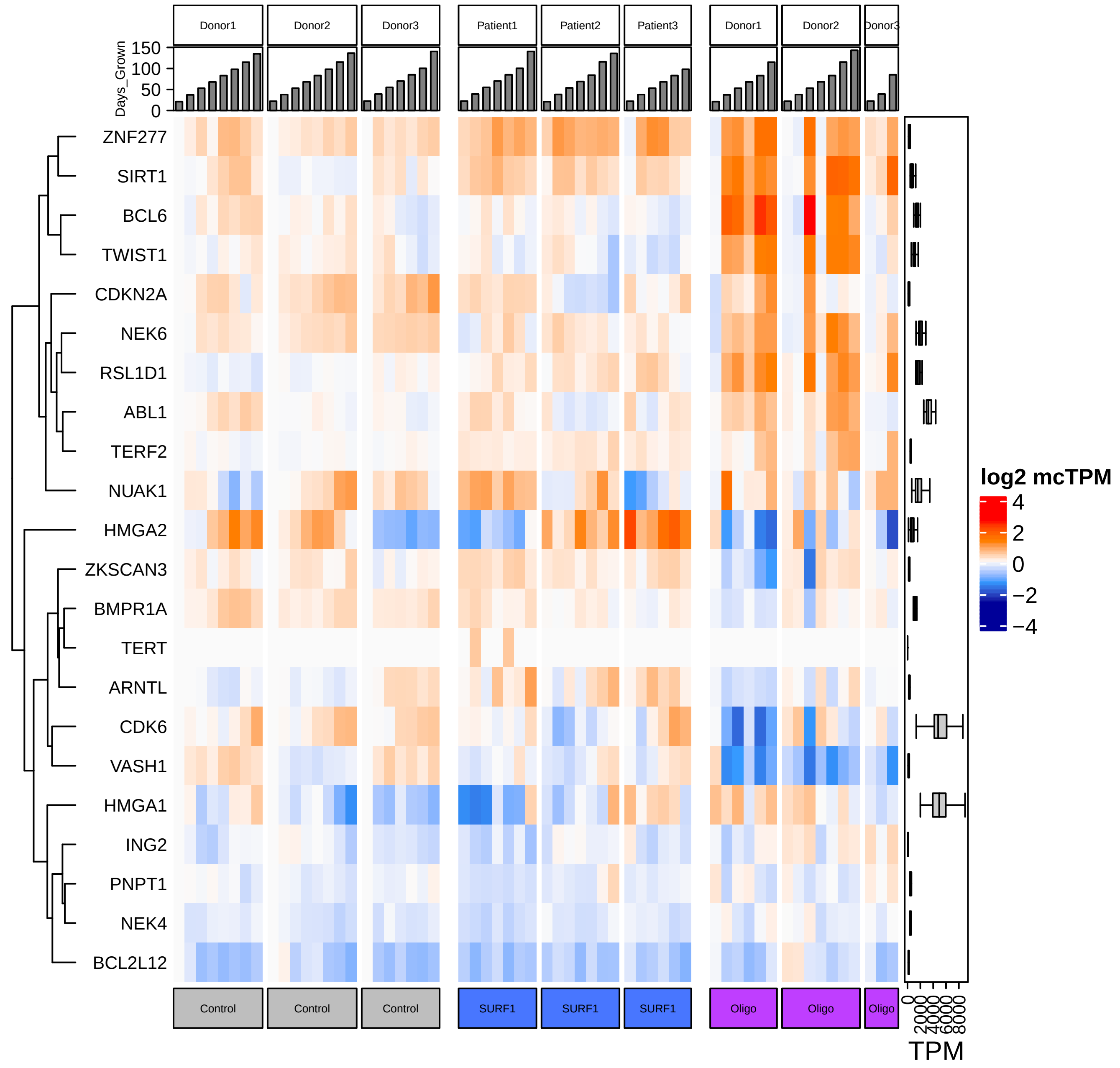

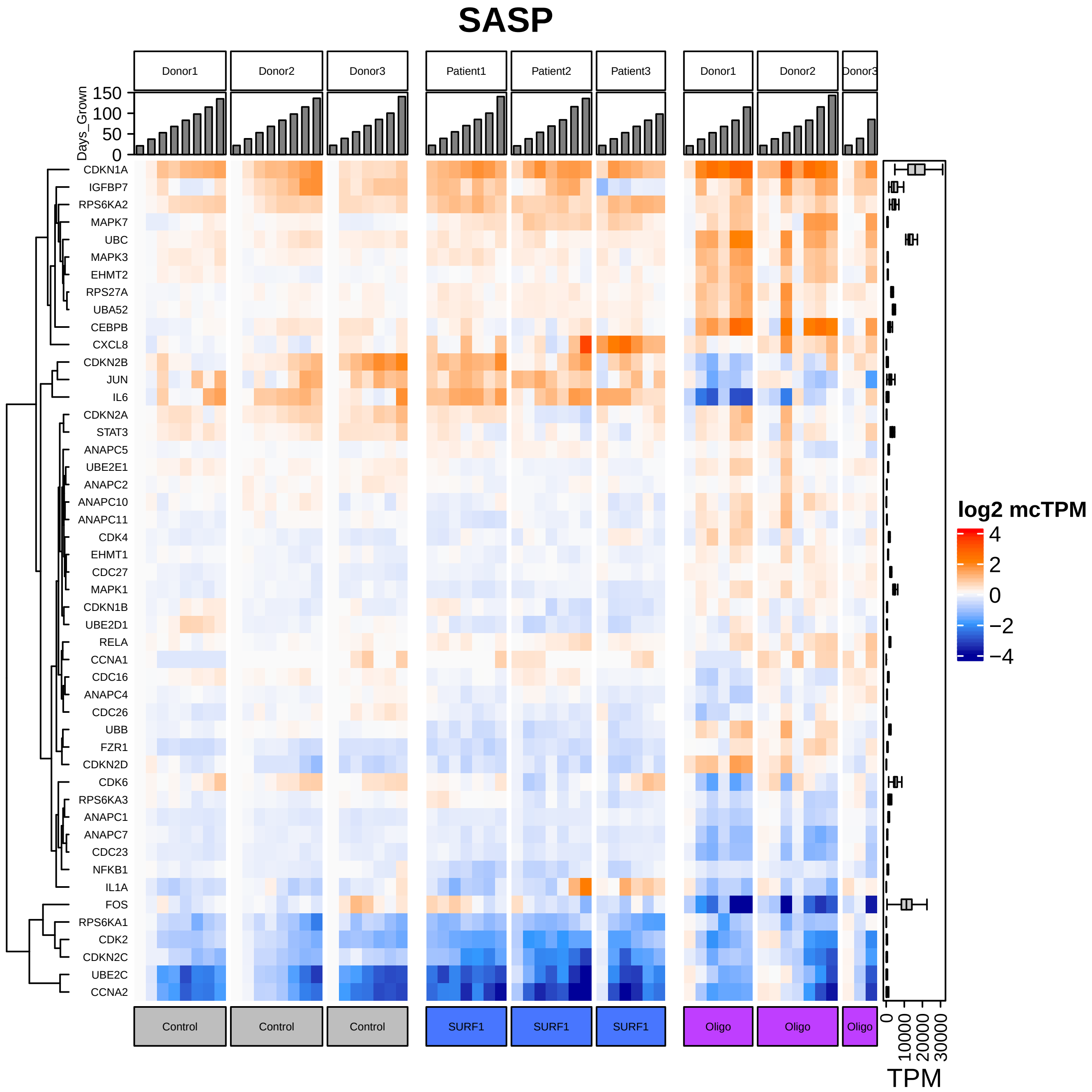

SASP
Donor1
Donor2
Donor3
Patient1
Patient2
Patient3
Donor1
Donor2
Donor3
Days_Grown
150
100
50
0
CDKN1A
IGFBP7
RPS6KA2
MAPK7
UBC
MAPK3
EHMT2
RPS27A
UBA52
CEBPB
CXCL8
CDKN2B
JUN
IL6
CDKN2A
STAT3
ANAPC5
UBE2E1
ANAPC2
ANAPC10
ANAPC11
CDK4
EHMT1
CDC27
MAPK1
CDKN1B
UBE2D1
RELA
CCNA1
CDC16
ANAPC4
CDC26
UBB
FZR1
CDKN2D
CDK6
RPS6KA3
ANAPC1
ANAPC7
CDC23
NFKB1
IL1A
FOS
RPS6KA1
CDK2
CDKN2C
UBE2C
CCNA2
log2 mcTPM
4
2
0
−2
−4
Control
Control
Control
SURF1
SURF1
SURF1
Oligo
Oligo
Oligo
0
10000
20000
30000
TPM

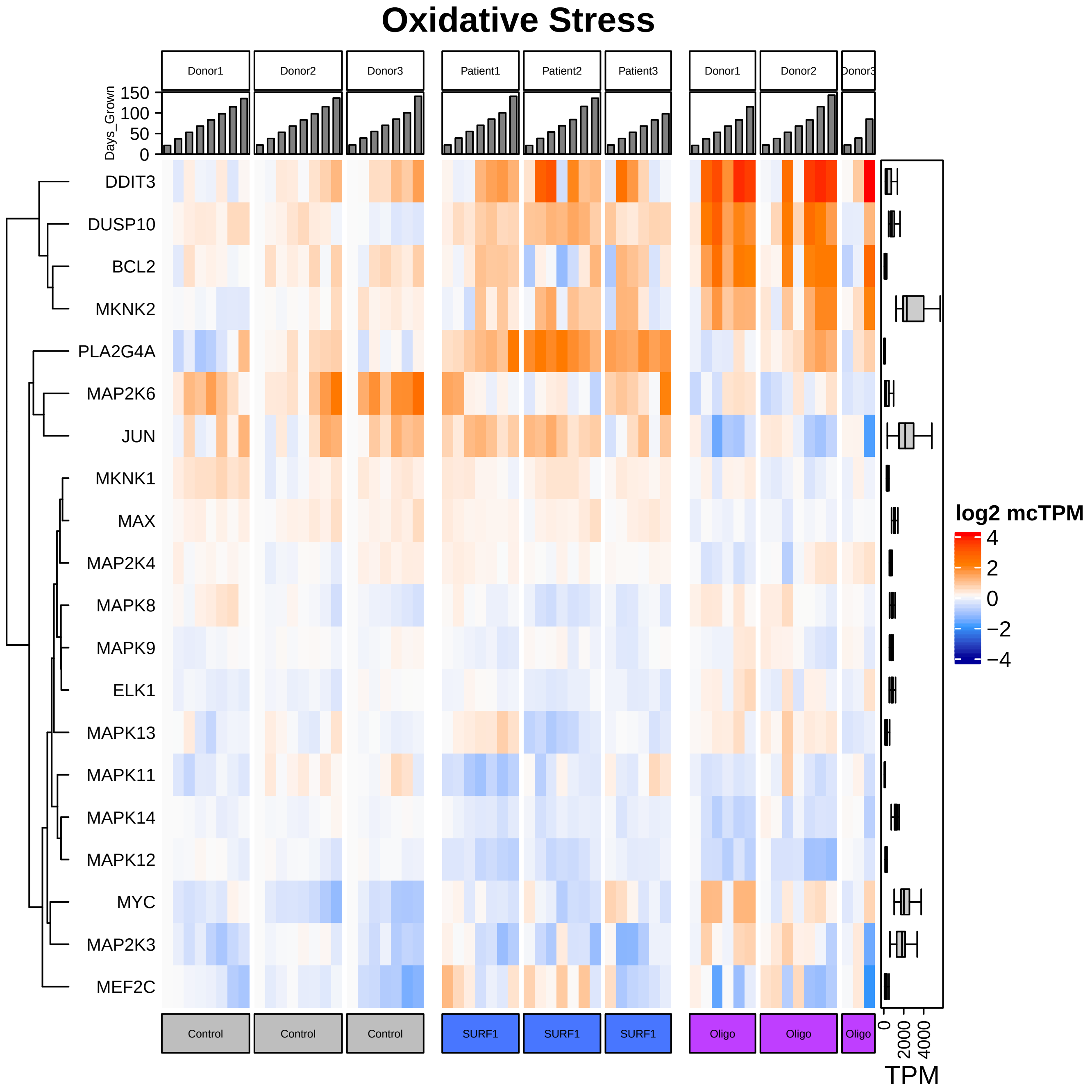

Oxidative Stress
Donor1
Donor2
Donor3
Patient1
Patient2
Patient3
Donor1
Donor2
Donor3
Days_Grown
150
100
50
0
DDIT3
DUSP10
BCL2
MKNK2
PLA2G4A
MAP2K6
JUN
MKNK1
MAX
MAP2K4
MAPK8
MAPK9
ELK1
MAPK13
MAPK11
MAPK14
MAPK12
MYC
MAP2K3
MEF2C
log2 mcTPM
4
2
0
−2
−4
Control
Control
Control
SURF1
SURF1
SURF1
Oligo
Oligo
Oligo
0
2000
4000
TPM

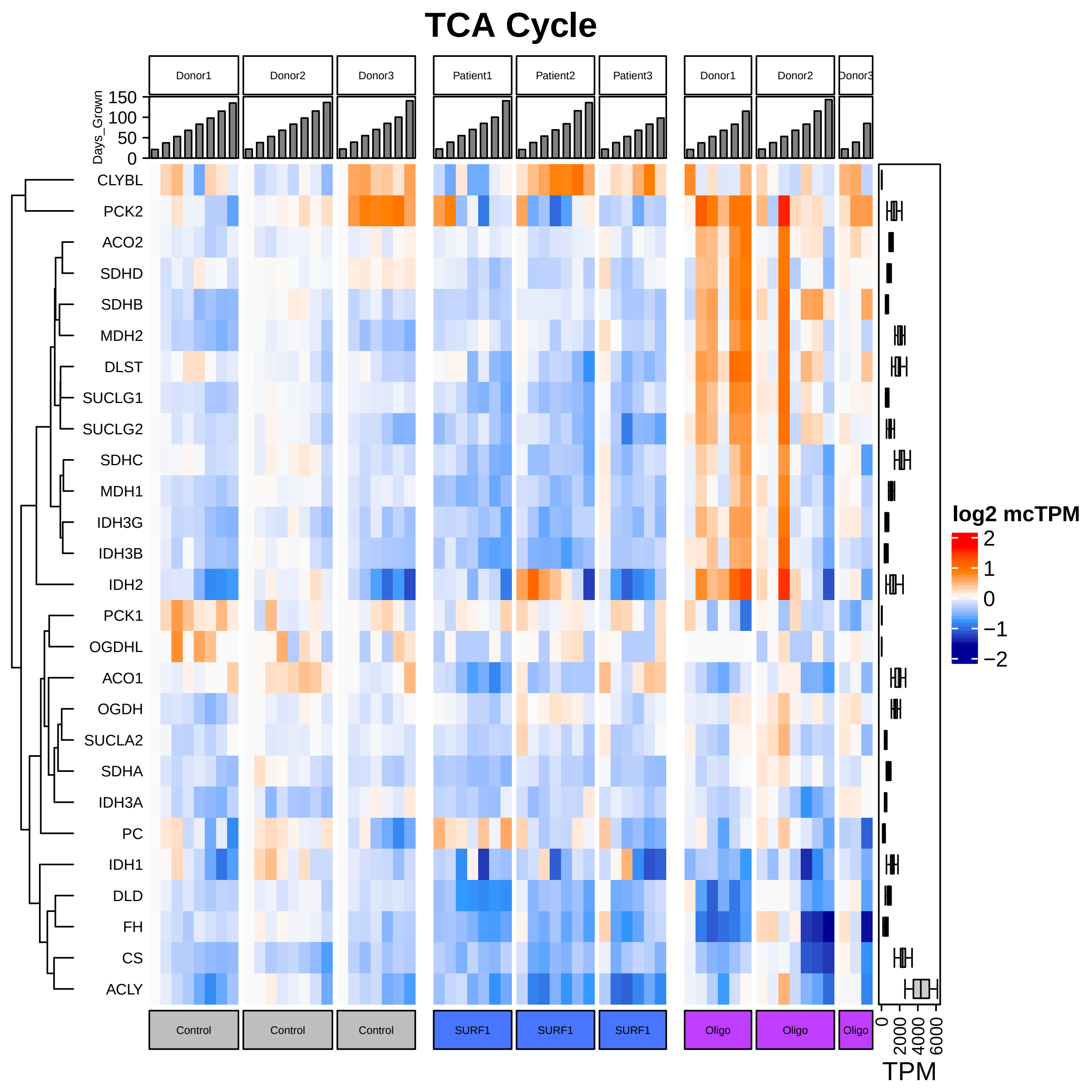
TCA Cycle
Donor1
Donor2
Donor3
Patient1
Patient2
Patient3
Donor1
Donor2
Donor3
Days_Grown
150
100
50
0
CLYBL
PCK2
ACO2
SDHD
SDHB
MDH2
DLST
SUCLG1
SUCLG2
SDHC
MDH1
IDH3G
IDH3B
IDH2
PCK1
OGDHL
ACO1
OGDH
SUCLA2
SDHA
IDH3A
PC
IDH1
DLD
FH
CS
ACLY
log2 mcTPM
2
1
0
−1
−2
Control
Control
Control
SURF1
SURF1
SURF1
Oligo
Oligo
Oligo
0
2000
4000
6000
TPM

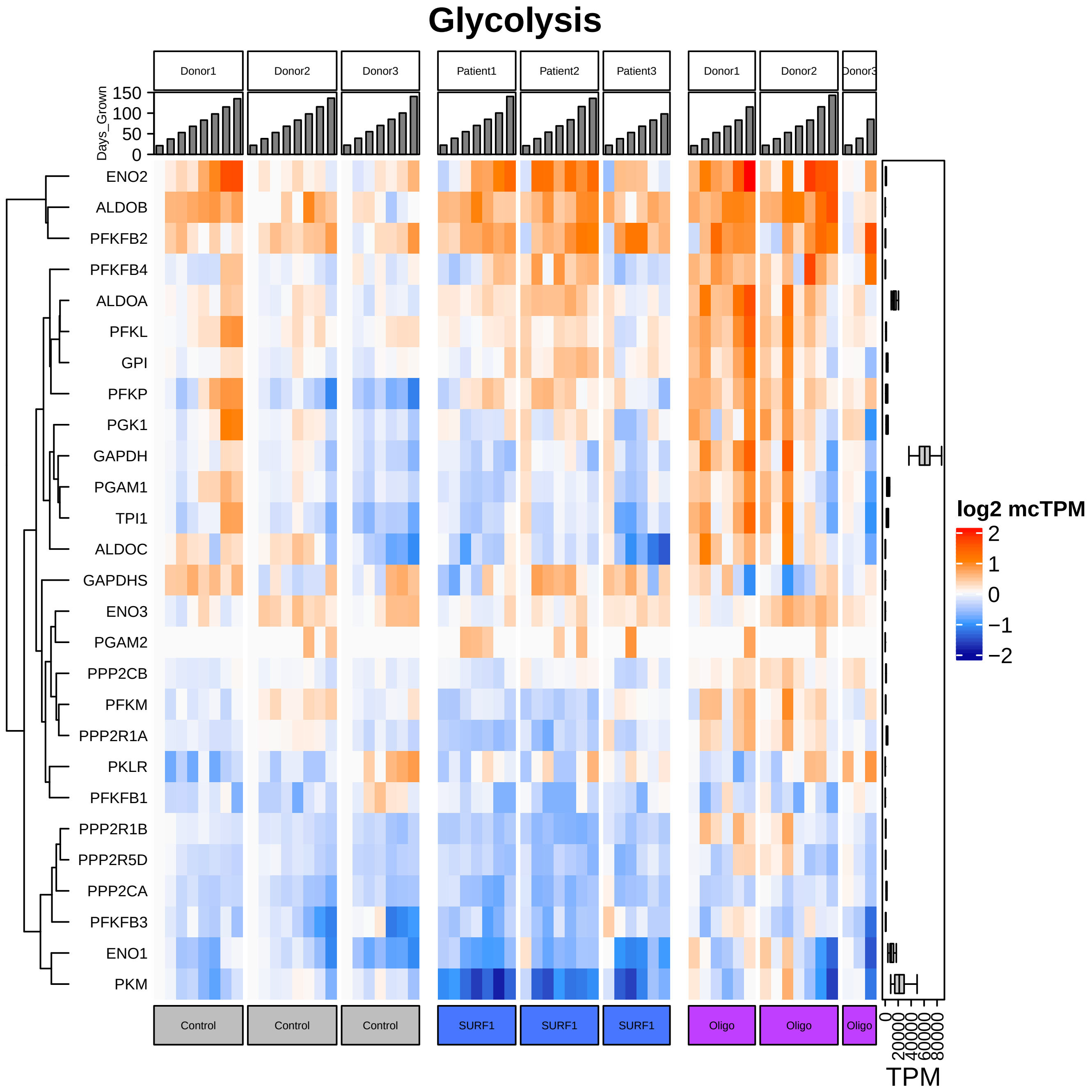

Glycolysis
Donor1
Donor2
Donor3
Patient1
Patient2
Patient3
Donor1
Donor2
Donor3
Days_Grown
150
100
50
0
ENO2
ALDOB
PFKFB2
PFKFB4
ALDOA
PFKL
GPI
PFKP
PGK1
GAPDH
PGAM1
TPI1
ALDOC
GAPDHS
ENO3
PGAM2
PPP2CB
PFKM
PPP2R1A
PKLR
PFKFB1
PPP2R1B
PPP2R5D
PPP2CA
PFKFB3
ENO1
PKM
Control
Control
Control
SURF1
SURF1
SURF1
Oligo
Oligo
Oligo
0
20000
40000
60000
80000
TPM
log2 mcTPM
2
1
0
−1
−2

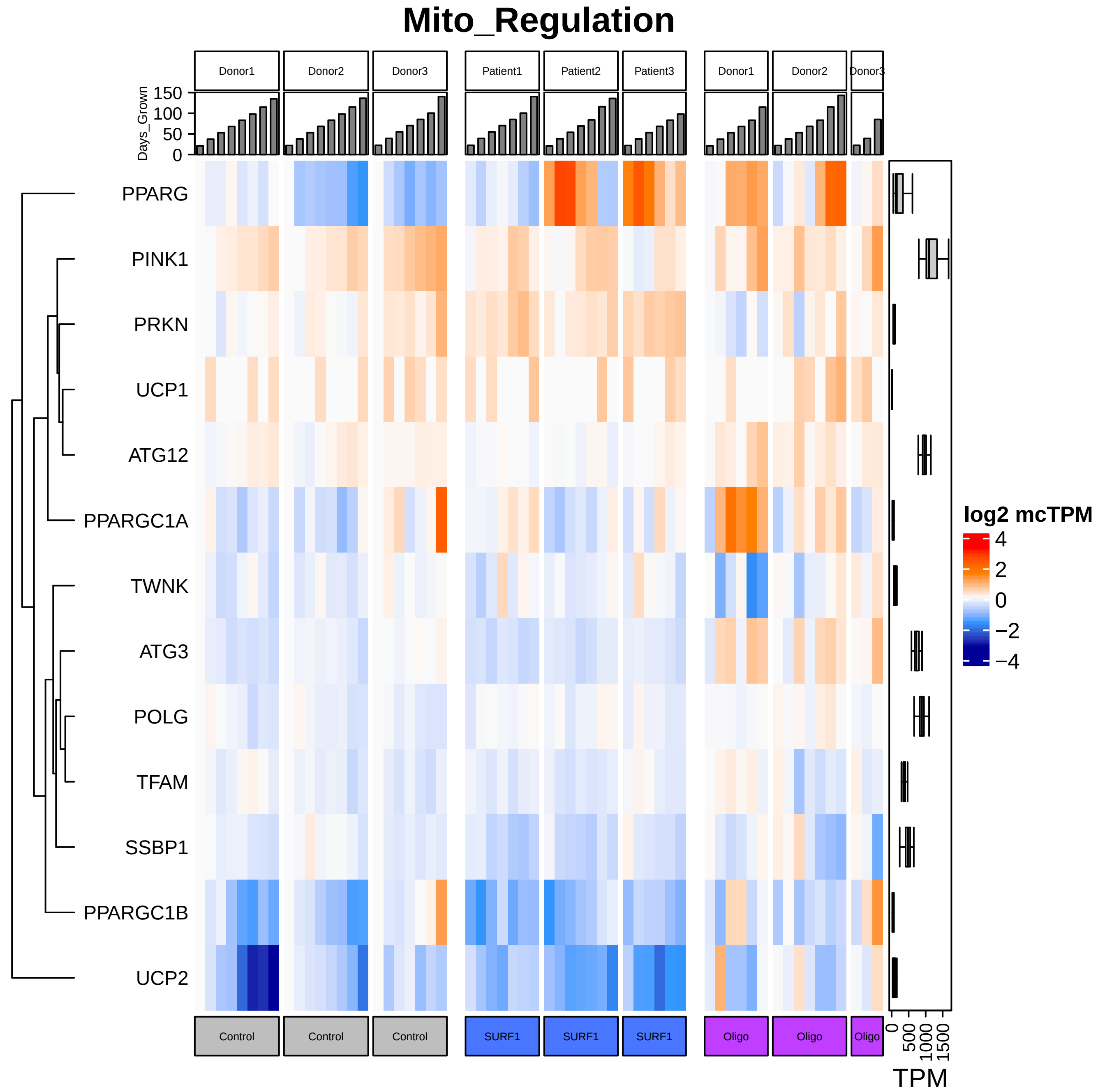

Mito_Regulation
Donor1
Donor2
Donor3
Patient1
Patient2
Patient3
Donor1
Donor2
Donor3
Days_Grown
150
100
50
0
PPARG
PINK1
PRKN
UCP1
ATG12
PPARGC1A
TWNK
ATG3
POLG
TFAM
SSBP1
PPARGC1B
UCP2
log2 mcTPM
4
2
0
−2
−4
Control
Control
Control
SURF1
SURF1
SURF1
Oligo
Oligo
Oligo
0
500
1000
1500
TPM

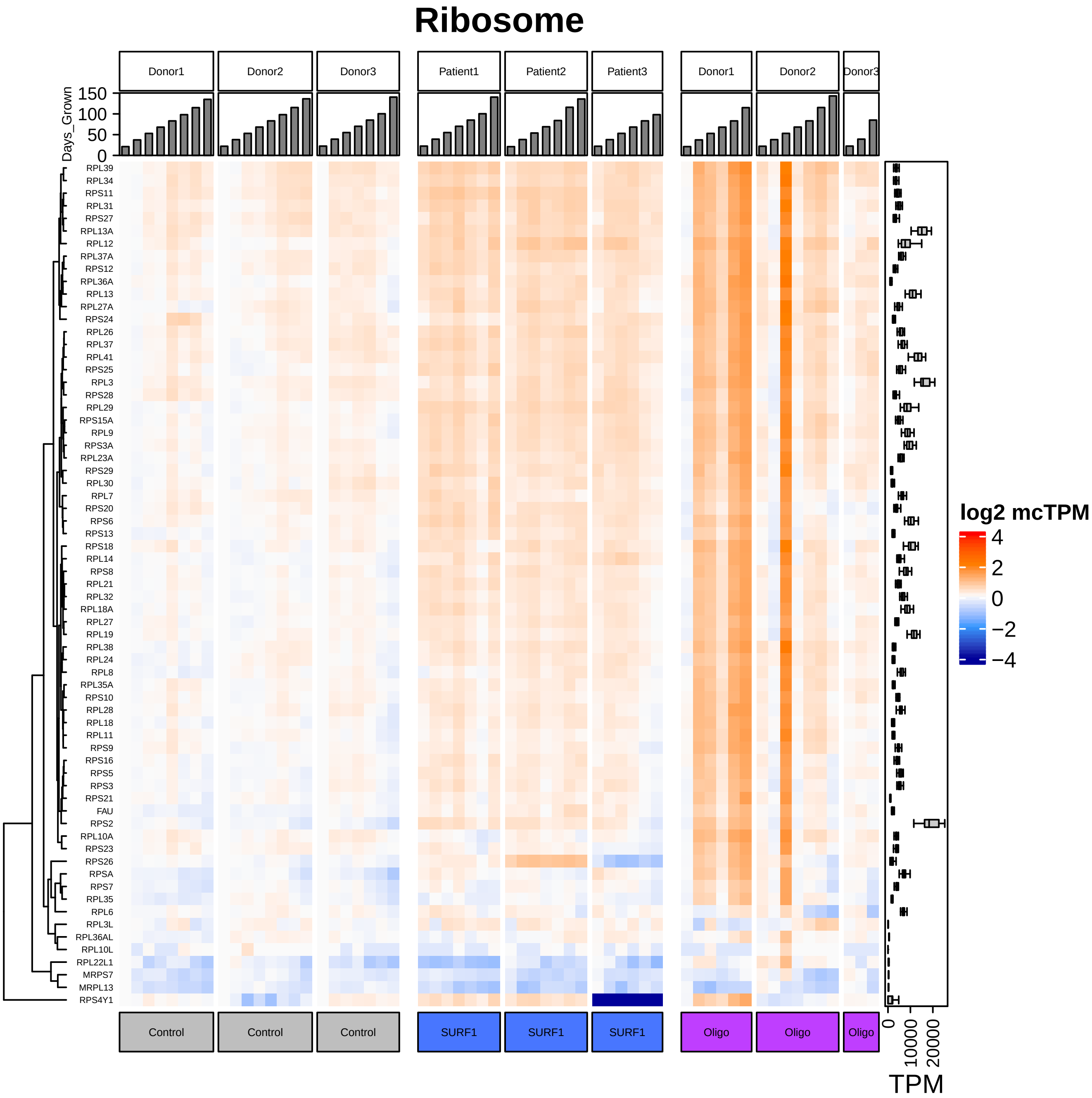

Ribosome
Donor1
Donor2
Donor3
Patient1
Patient2
Patient3
Donor1
Donor2
Donor3
Days_Grown
150
100
50
0
RPL39
RPL34
RPS11
RPL31
RPS27
RPL13A
RPL12
RPL37A
RPS12
RPL36A
RPL13
RPL27A
RPS24
RPL26
RPL37
RPL41
RPS25
RPL3
RPS28
RPL29
RPS15A
RPL9
RPS3A
RPL23A
RPS29
RPL30
RPL7
RPS20
RPS6
RPS13
RPS18
RPL14
RPS8
RPL21
RPL32
RPL18A
RPL27
RPL19
RPL38
RPL24
RPL8
RPL35A
RPS10
RPL28
RPL18
RPL11
RPS9
RPS16
RPS5
RPS3
RPS21
FAU
RPS2
RPL10A
RPS23
RPS26
RPSA
RPS7
RPL35
RPL6
RPL3L
RPL36AL
RPL10L
RPL22L1
MRPS7
MRPL13
RPS4Y1
Control
Control
Control
SURF1
SURF1
SURF1
Oligo
Oligo
Oligo
log2 mcTPM
4
2
0
−2
−4
0
10000
20000
TPM

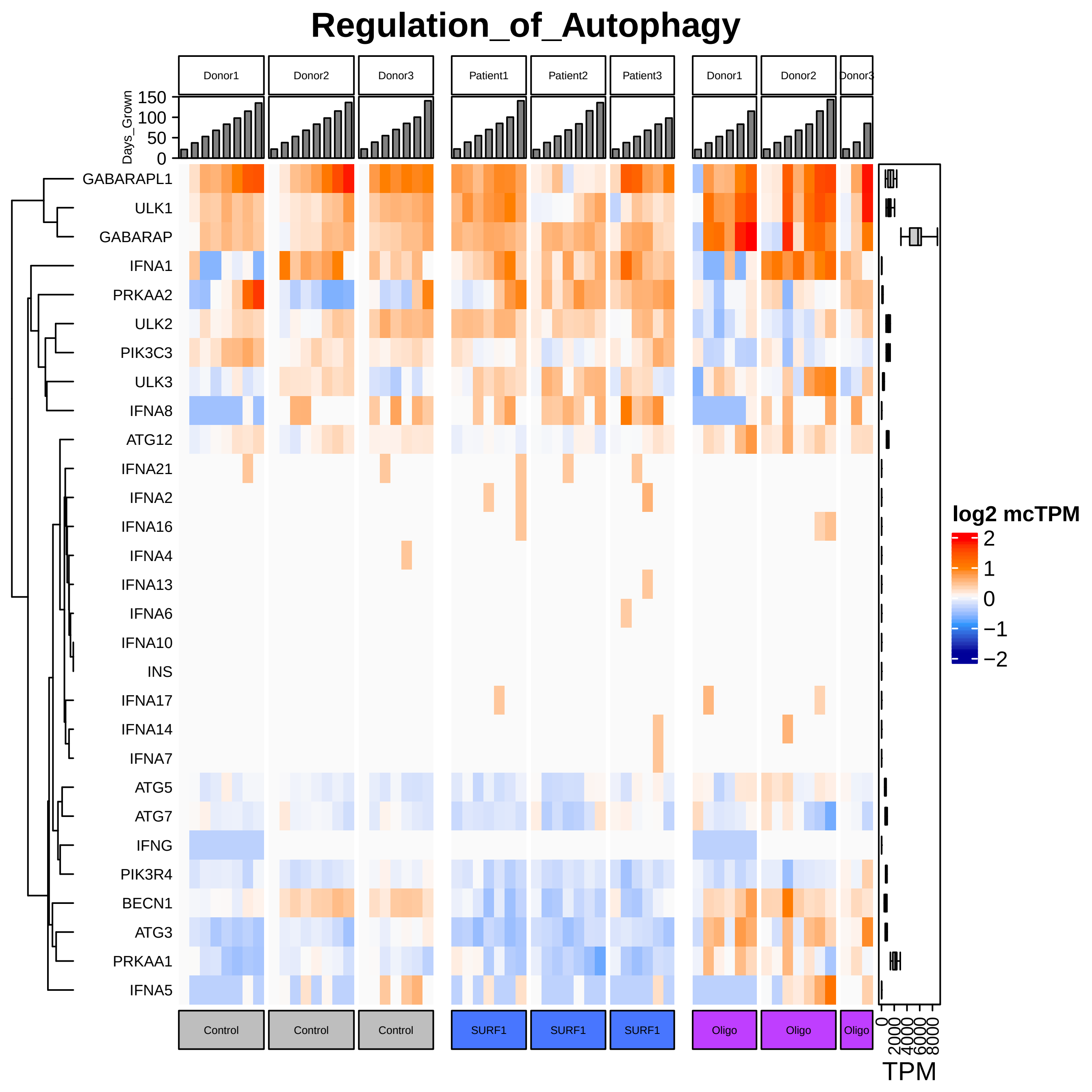
Regulation_of_Autophagy
Donor1
Donor2
Donor3
Patient1
Patient2
Patient3
Donor1
Donor2
Donor3
Days_Grown
150
100
50
0
GABARAPL1
ULK1
GABARAP
IFNA1
PRKAA2
ULK2
PIK3C3
ULK3
IFNA8
ATG12
IFNA21
IFNA2
IFNA16
IFNA4
IFNA13
IFNA6
IFNA10
INS
IFNA17
IFNA14
IFNA7
ATG5
ATG7
IFNG
PIK3R4
BECN1
ATG3
PRKAA1
IFNA5
log2 mcTPM
2
1
0
−1
−2
Control
Control
Control
SURF1
SURF1
SURF1
Oligo
Oligo
Oligo
0
2000
4000
6000
8000
TPM

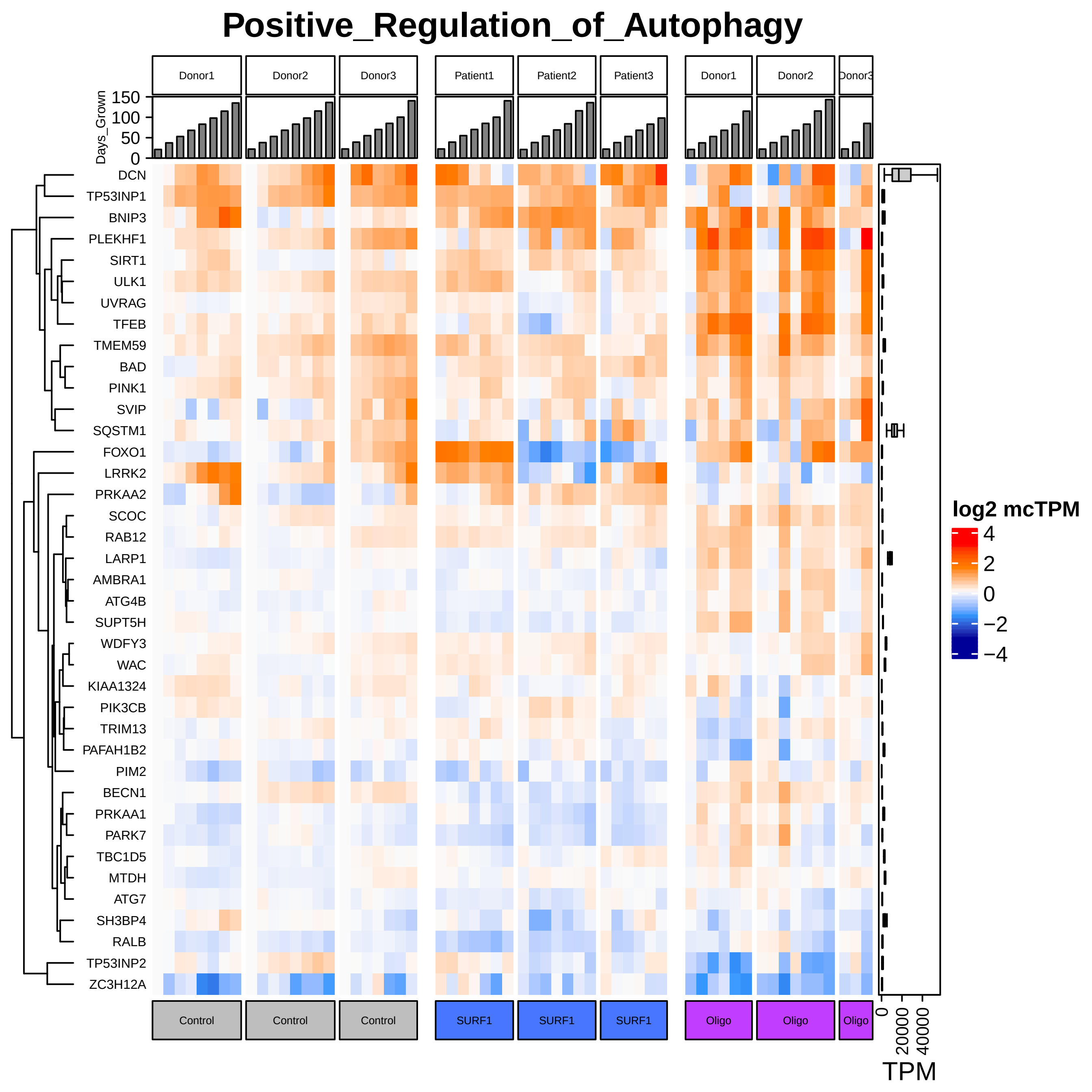

Positive_Regulation_of_Autophagy
Donor1
Donor2
Donor3
Patient1
Patient2
Patient3
Donor1
Donor2
Donor3
Days_Grown
150
100
50
0
DCN
TP53INP1
BNIP3
PLEKHF1
SIRT1
ULK1
UVRAG
TFEB
TMEM59
BAD
PINK1
SVIP
SQSTM1
FOXO1
LRRK2
PRKAA2
SCOC
RAB12
LARP1
AMBRA1
ATG4B
SUPT5H
WDFY3
WAC
KIAA1324
PIK3CB
TRIM13
PAFAH1B2
PIM2
BECN1
PRKAA1
PARK7
TBC1D5
MTDH
ATG7
SH3BP4
RALB
TP53INP2
ZC3H12A
log2 mcTPM
4
2
0
−2
−4
Control
Control
Control
SURF1
SURF1
SURF1
Oligo
Oligo
Oligo
0
20000
40000
TPM

### Negative_Regulation_of_Autophagy

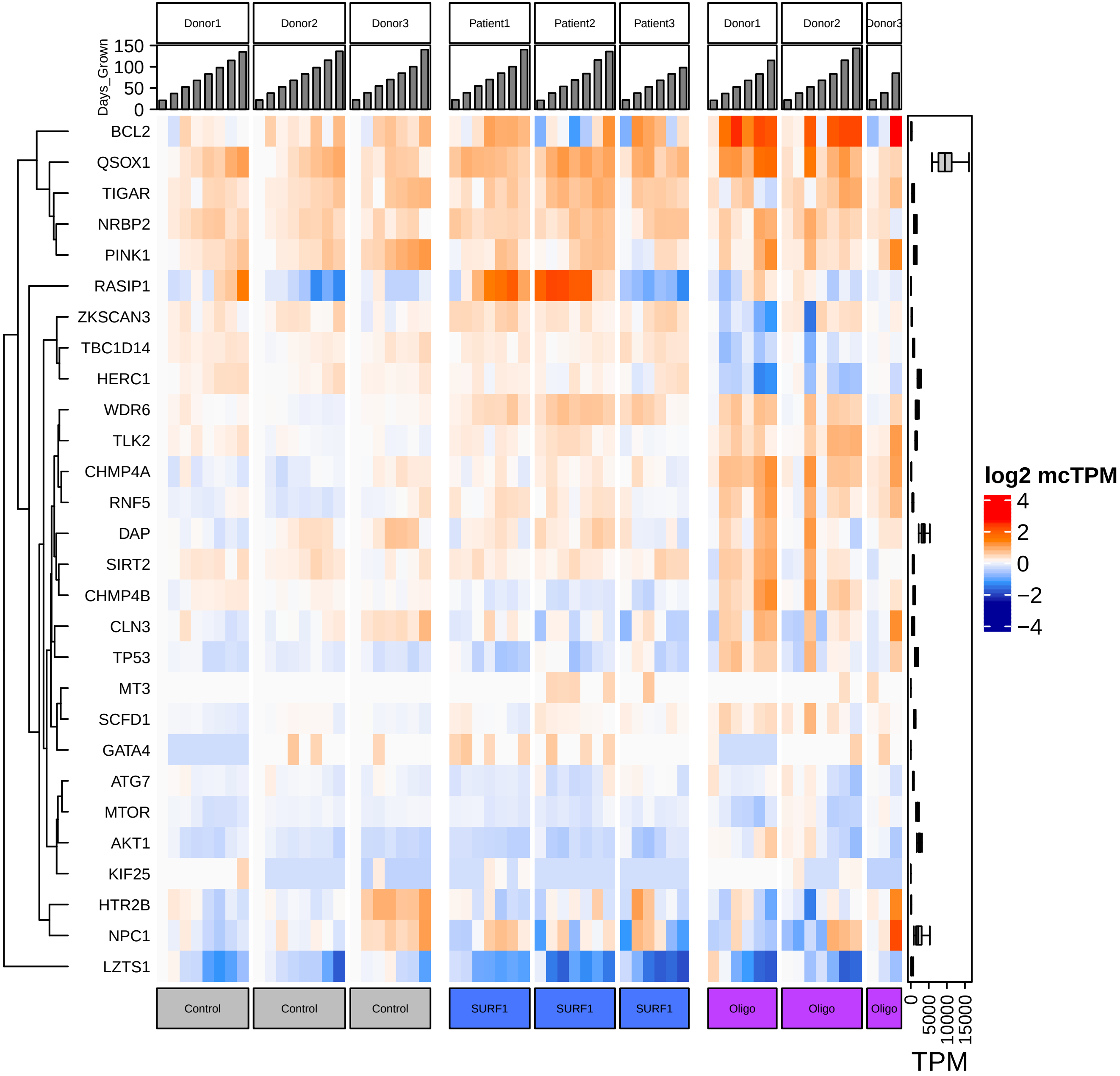

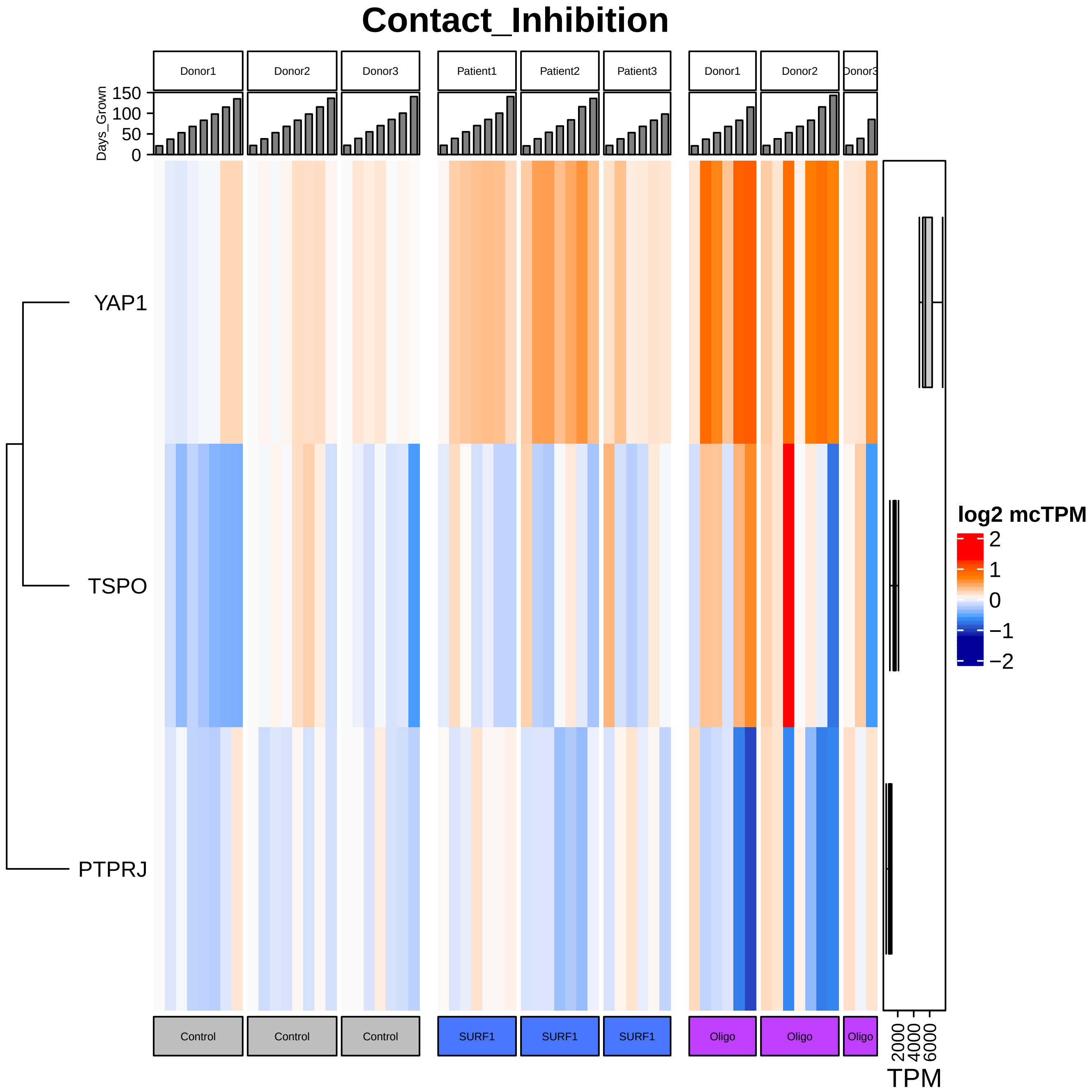

Contact_Inhibition
Donor1
Donor2
Donor3
Patient1
Patient2
Patient3
Donor1
Donor2
Donor3
Days_Grown
150
100
50
0
YAP1
TSPO
PTPRJ
log2 mcTPM
2
1
0
−1
−2
Control
Control
Control
SURF1
SURF1
SURF1
Oligo
Oligo
Oligo
2000
4000
6000
TPM

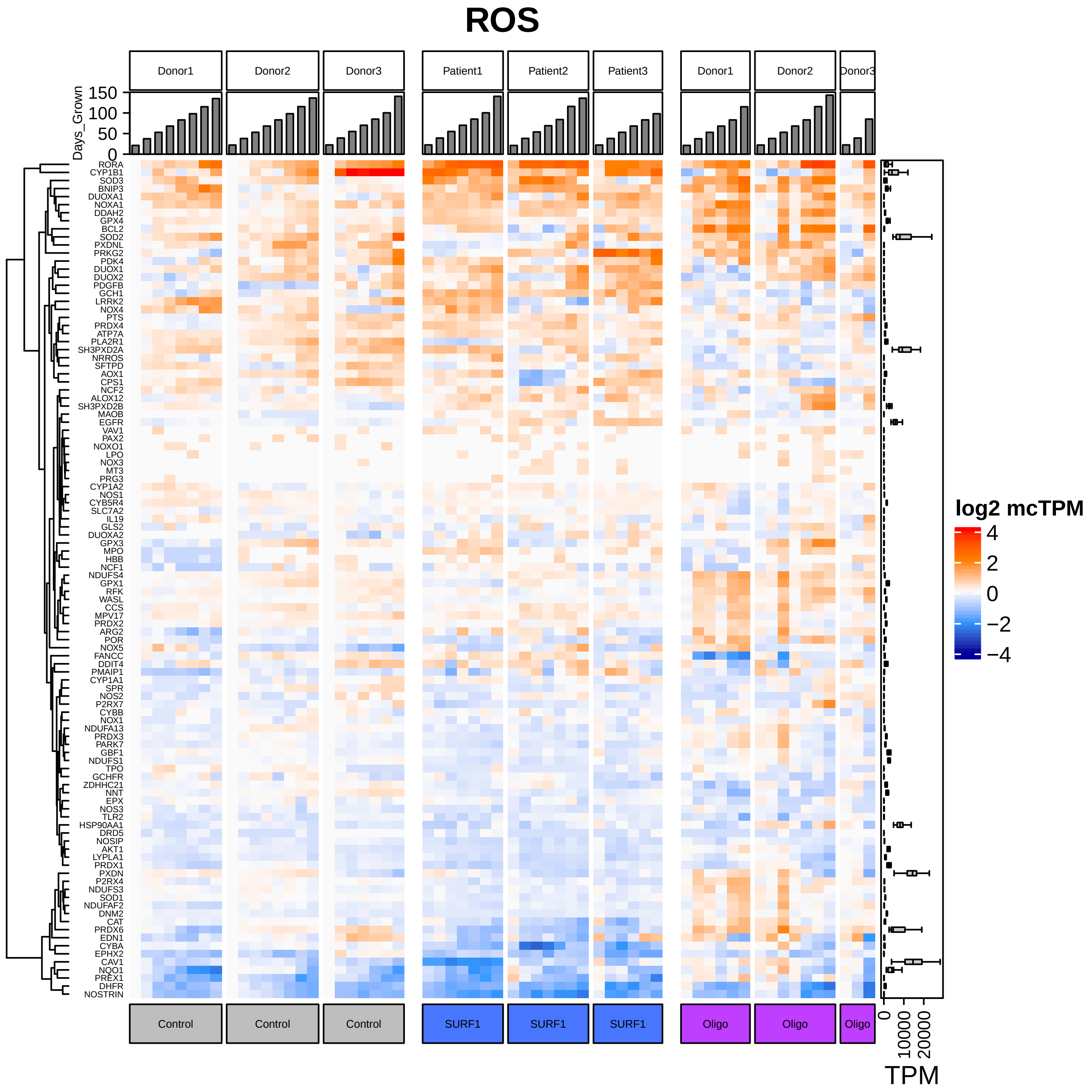
ROS
Donor1
Donor2
Donor3
Patient1
Patient2
Patient3
Donor1
Donor2
Donor3
Days_Grown
150
100
50
0
RORA
CYP1B1
SOD3
BNIP3
DUOXA1
NOXA1
DDAH2
GPX4
BCL2
SOD2
PXDNL
PRKG2
PDK4
DUOX1
DUOX2
PDGFB
GCH1
LRRK2
NOX4
PTS
PRDX4
ATP7A
PLA2R1
SH3PXD2A
NRROS
SFTPD
AOX1
CPS1
NCF2
ALOX12
SH3PXD2B
MAOB
EGFR
VAV1
PAX2
NOXO1
LPO
NOX3
MT3
PRG3
CYP1A2
NOS1
CYB5R4
SLC7A2
IL19
GLS2
DUOXA2
GPX3
MPO
HBB
NCF1
NDUFS4
GPX1
RFK
WASL
CCS
MPV17
PRDX2
ARG2
POR
NOX5
FANCC
DDIT4
PMAIP1
CYP1A1
SPR
NOS2
P2RX7
CYBB
NOX1
NDUFA13
PRDX3
PARK7
GBF1
NDUFS1
TPO
GCHFR
ZDHHC21
NNT
EPX
NOS3
TLR2
HSP90AA1
DRD5
NOSIP
AKT1
LYPLA1
PRDX1
PXDN
P2RX4
NDUFS3
SOD1
NDUFAF2
DNM2
CAT
PRDX6
EDN1
CYBA
EPHX2
CAV1
NQO1
PREX1
DHFR
NOSTRIN
log2 mcTPM
4
2
0
−2
−4
Control
Control
Control
SURF1
SURF1
SURF1
Oligo
Oligo
Oligo
0
10000
20000
TPM

### Pentose_Phosphate

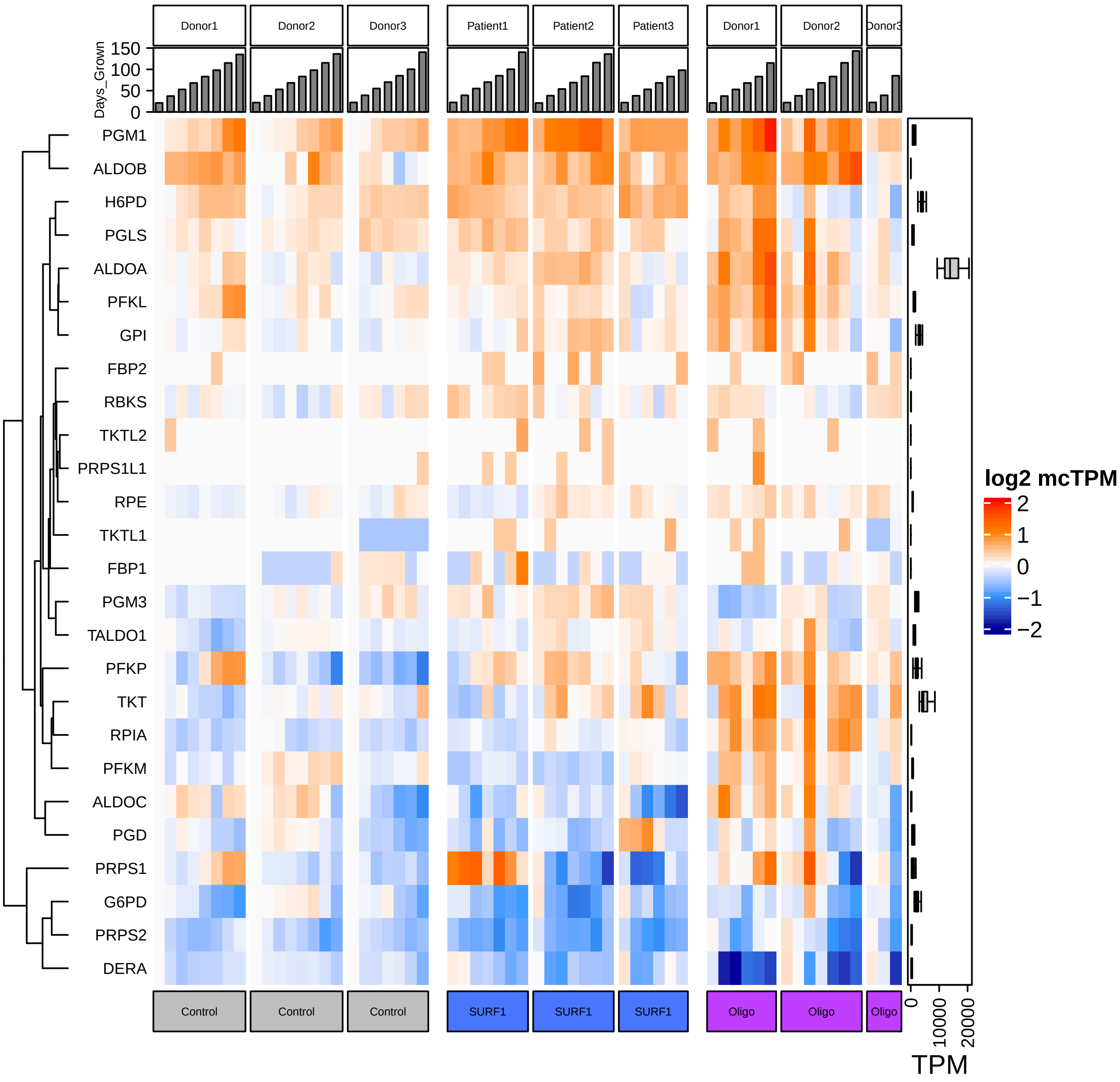

### Mitophagy

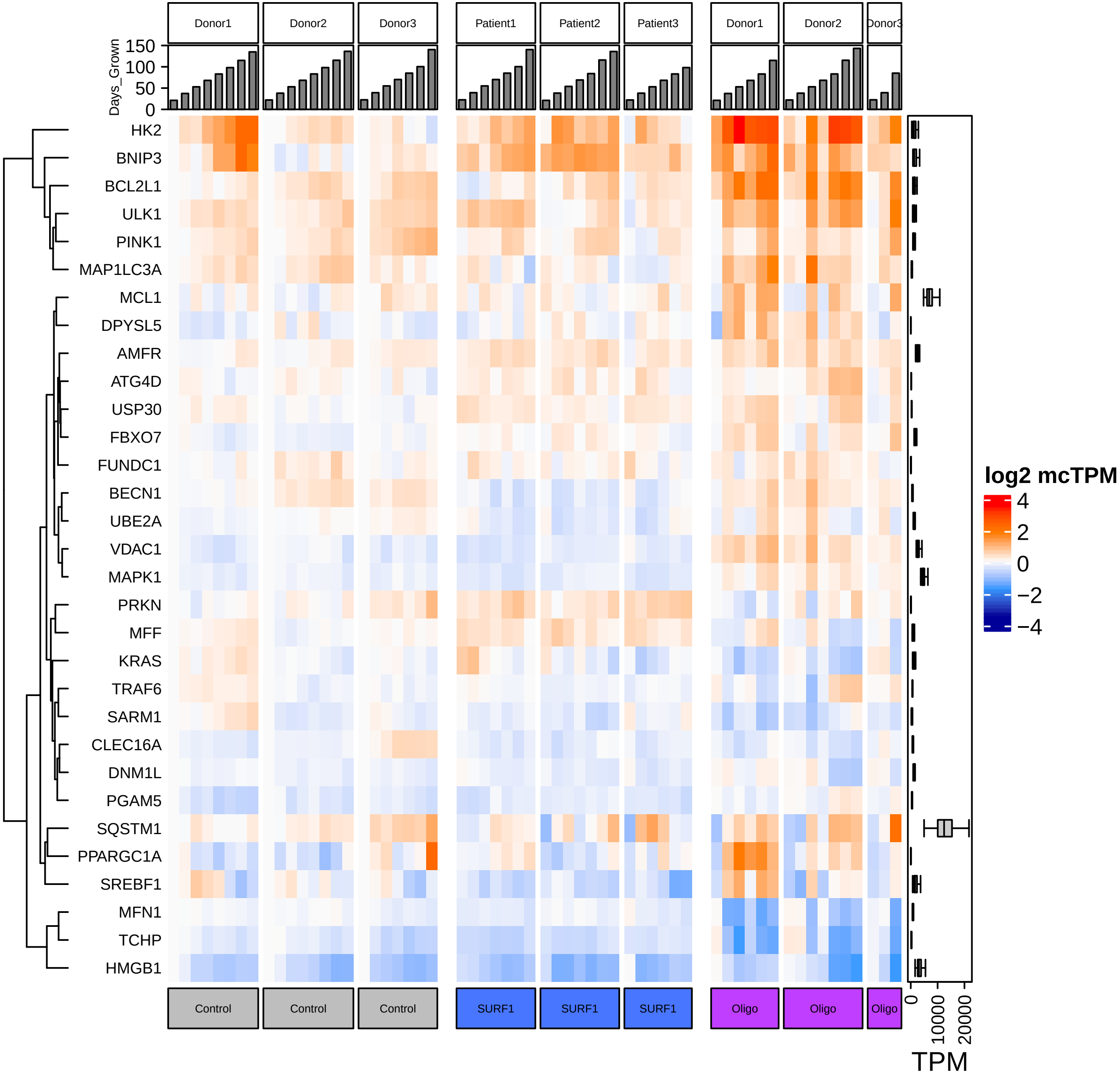

Donor1
Donor2
Donor3
Patient1
Patient2
Patient3
Donor1
Donor2
Donor3
Days_Grown
150
100
50
0
HK2
BNIP3
BCL2L1
ULK1
PINK1
MAP1LC3A
MCL1
DPYSL5
AMFR
ATG4D
USP30
FBXO7
FUNDC1
BECN1
UBE2A
VDAC1
MAPK1
PRKN
MFF
KRAS
TRAF6
SARM1
CLEC16A
DNM1L
PGAM5
SQSTM1
PPARGC1A
SREBF1
MFN1
TCHP
HMGB1
Control
Control
Control
SURF1
SURF1
SURF1
Oligo
Oligo
Oligo
0
10000
20000
TPM
log2 mcTPM
4
2
0
−2
−4

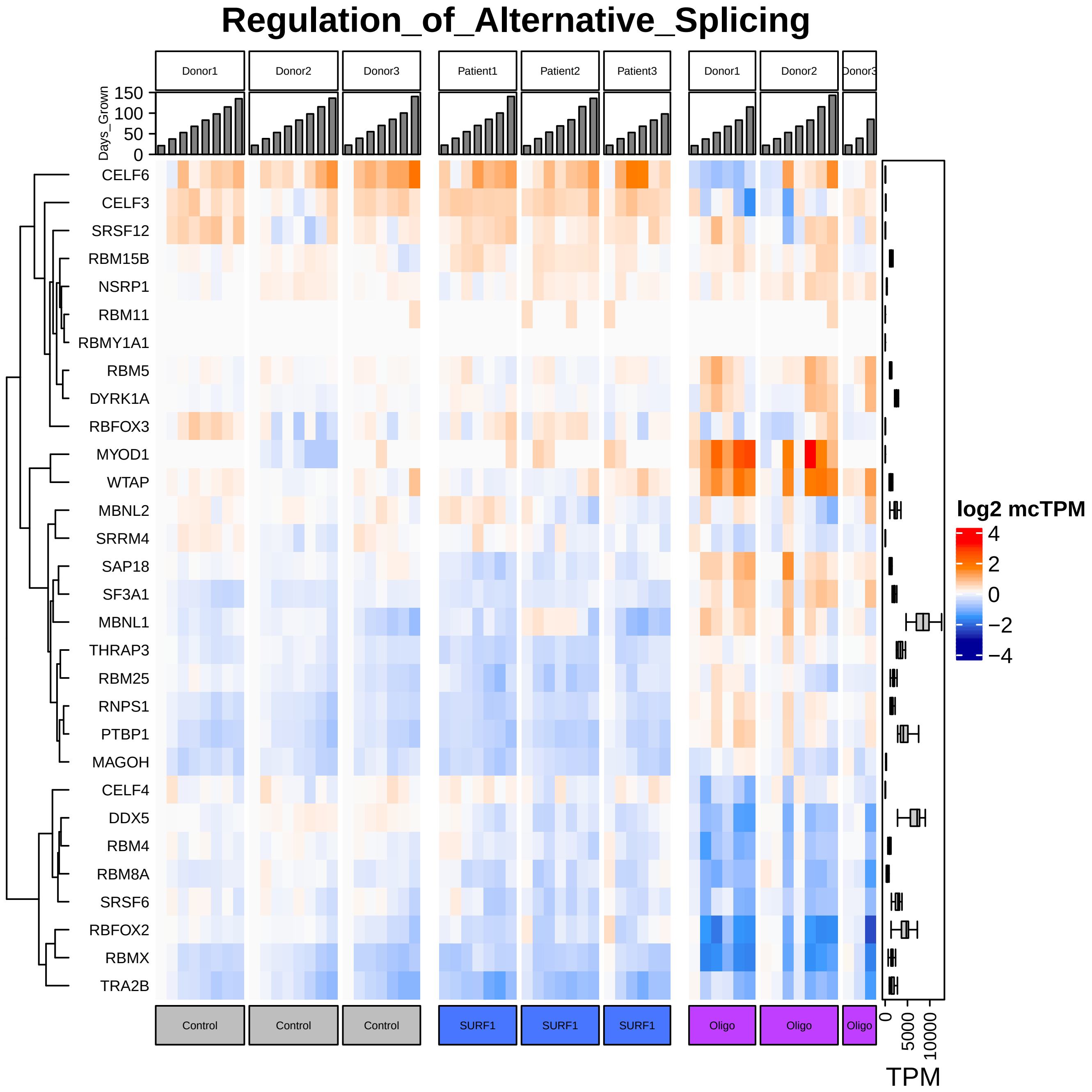

Regulation_of_Alternative_Splicing
Donor1
Donor2
Donor3
Patient1
Patient2
Patient3
Donor1
Donor2
Donor3
Days_Grown
150
100
50
0
CELF6
CELF3
SRSF12
RBM15B
NSRP1
RBM11
RBMY1A1
RBM5
DYRK1A
RBFOX3
MYOD1
WTAP
MBNL2
SRRM4
SAP18
SF3A1
MBNL1
THRAP3
RBM25
RNPS1
PTBP1
MAGOH
CELF4
DDX5
RBM4
RBM8A
SRSF6
RBFOX2
RBMX
TRA2B
log2 mcTPM
4
2
0
−2
−4
Control
Control
Control
SURF1
SURF1
SURF1
Oligo
Oligo
Oligo
0
5000
10000
TPM

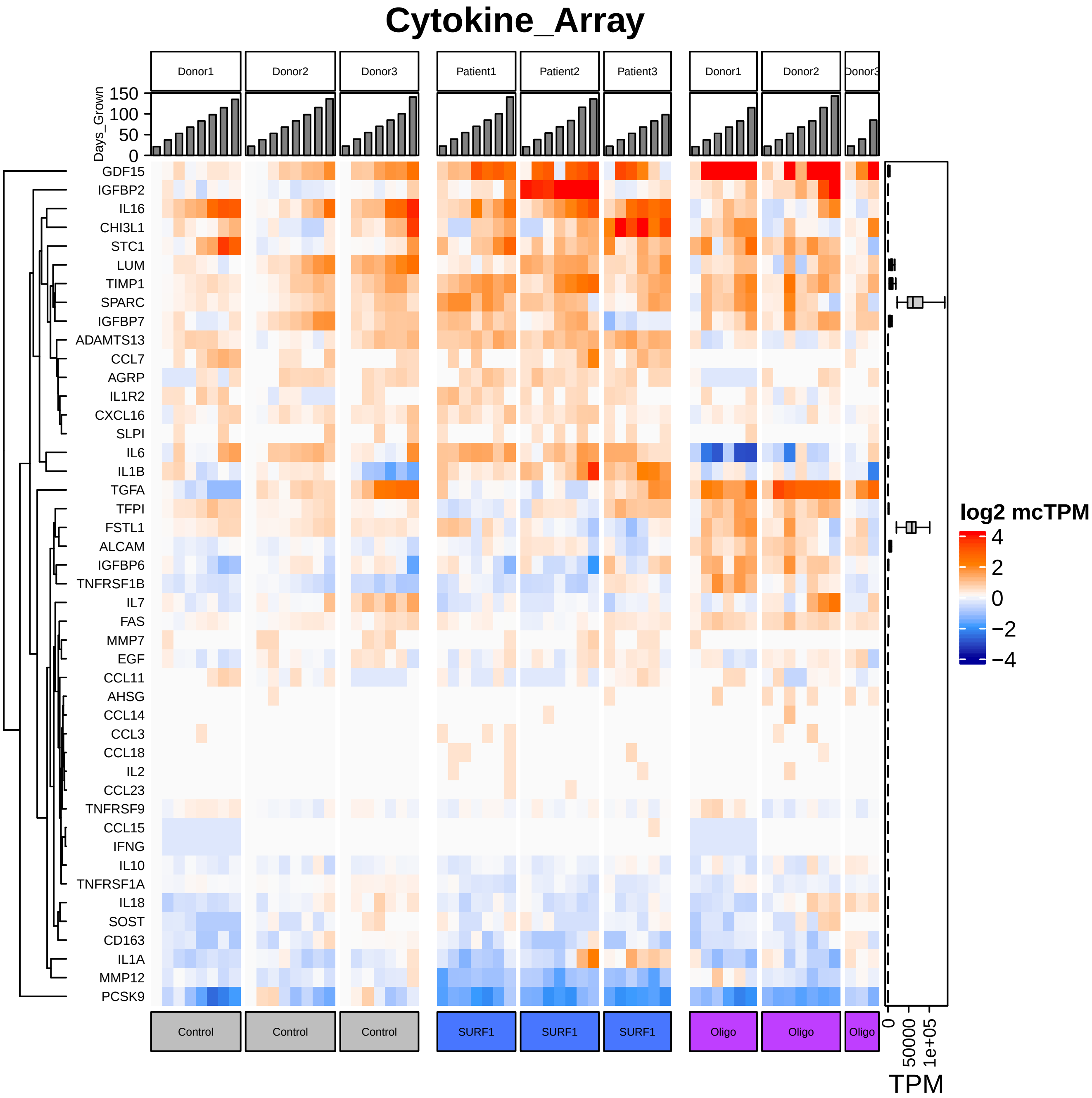

Cytokine_Array
Donor1
Donor2
Donor3
Patient1
Patient2
Patient3
Donor1
Donor2
Donor3
Days_Grown
150
100
50
0
GDF15
IGFBP2
IL16
CHI3L1
STC1
LUM
TIMP1
SPARC
IGFBP7
ADAMTS13
CCL7
AGRP
IL1R2
CXCL16
SLPI
IL6
IL1B
TGFA
TFPI
FSTL1
ALCAM
IGFBP6
TNFRSF1B
IL7
FAS
MMP7
EGF
CCL11
AHSG
CCL14
CCL3
CCL18
IL2
CCL23
TNFRSF9
CCL15
IFNG
IL10
TNFRSF1A
IL18
SOST
CD163
IL1A
MMP12
PCSK9
log2 mcTPM
4
2
0
−2
−4
Control
Control
Control
SURF1
SURF1
SURF1
Oligo
Oligo
Oligo
0
50000
1e+05
TPM

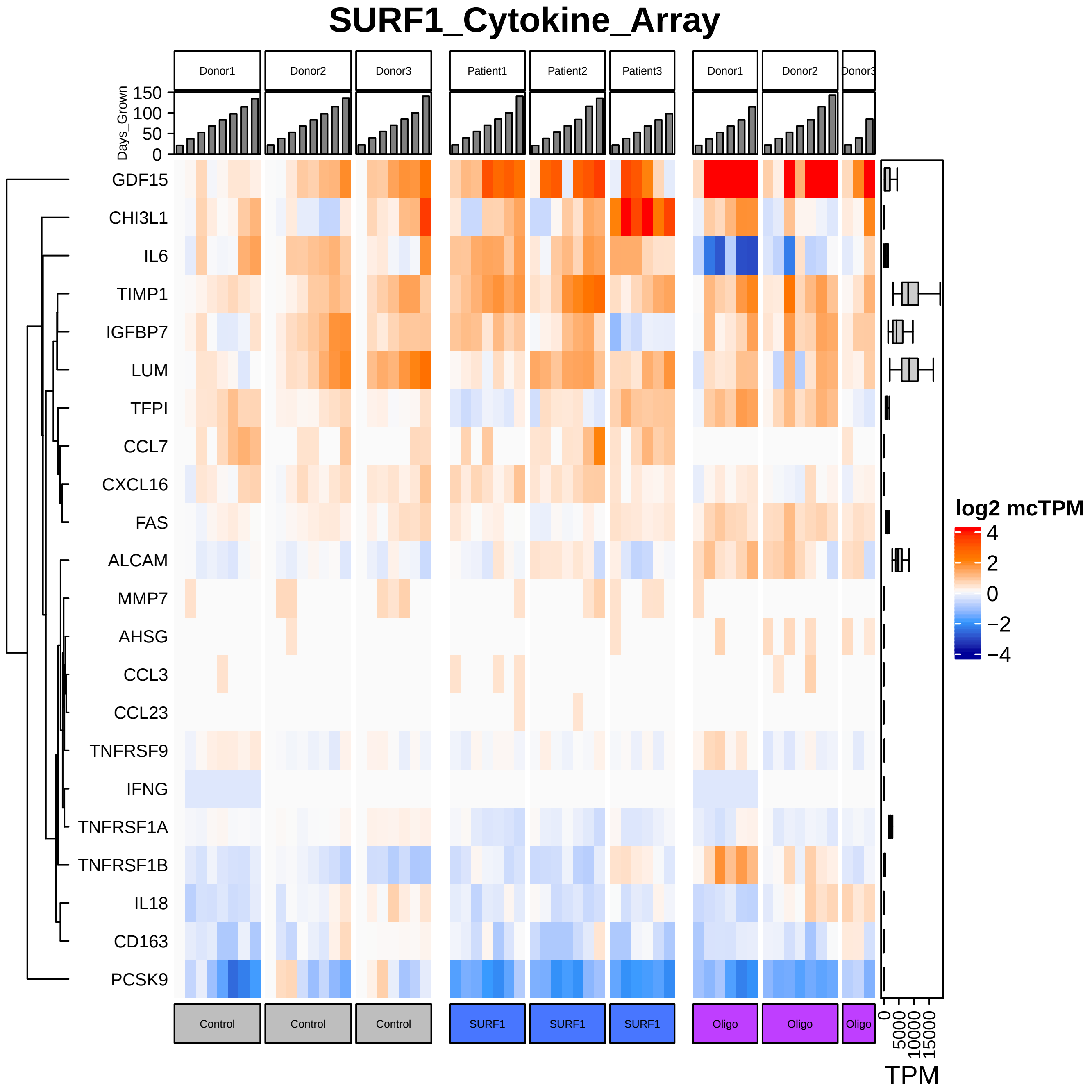
SURF1_Cytokine_Array
Donor1
Donor2
Donor3
Patient1
Patient2
Patient3
Donor1
Donor2
Donor3
Days_Grown
150
100
50
0
GDF15
CHI3L1
IL6
TIMP1
IGFBP7
LUM
TFPI
CCL7
CXCL16
FAS
ALCAM
MMP7
AHSG
CCL3
CCL23
TNFRSF9
IFNG
TNFRSF1A
TNFRSF1B
IL18
CD163
PCSK9
Control
Control
Control
SURF1
SURF1
SURF1
Oligo
Oligo
Oligo
0
5000
10000
15000
TPM
log2 mcTPM
4
2
0
−2
−4

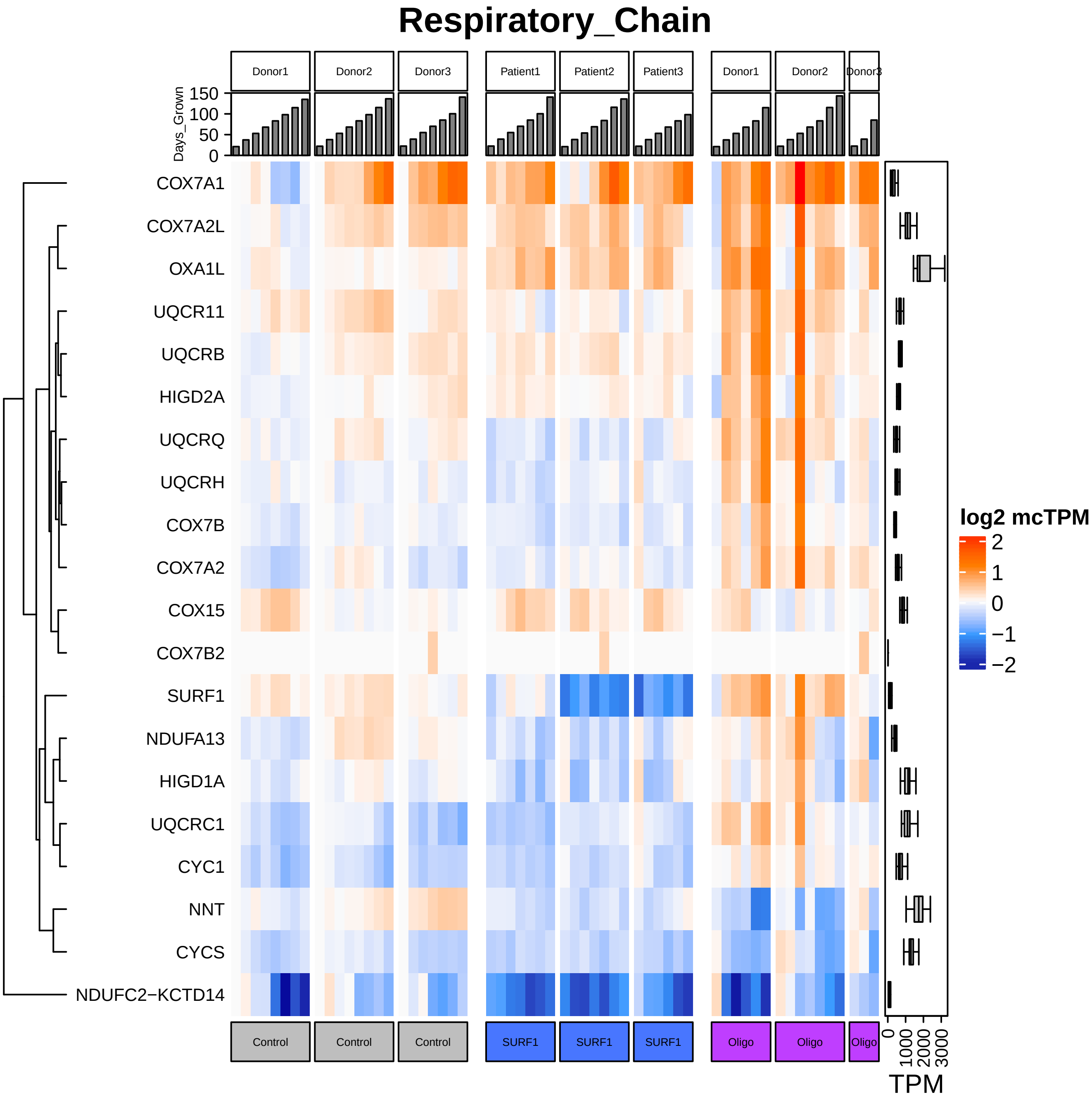

Respiratory_Chain
Donor1
Donor2
Donor3
Patient1
Patient2
Patient3
Donor1
Donor2
Donor3
Days_Grown
150
100
50
0
COX7A1
COX7A2L
OXA1L
UQCR11
UQCRB
HIGD2A
UQCRQ
UQCRH
COX7B
COX7A2
COX15
COX7B2
SURF1
NDUFA13
HIGD1A
UQCRC1
CYC1
NNT
CYCS
NDUFC2−KCTD14
log2 mcTPM
2
1
0
−1
−2
Control
Control
Control
SURF1
SURF1
SURF1
Oligo
Oligo
Oligo
0
1000
2000
3000
TPM

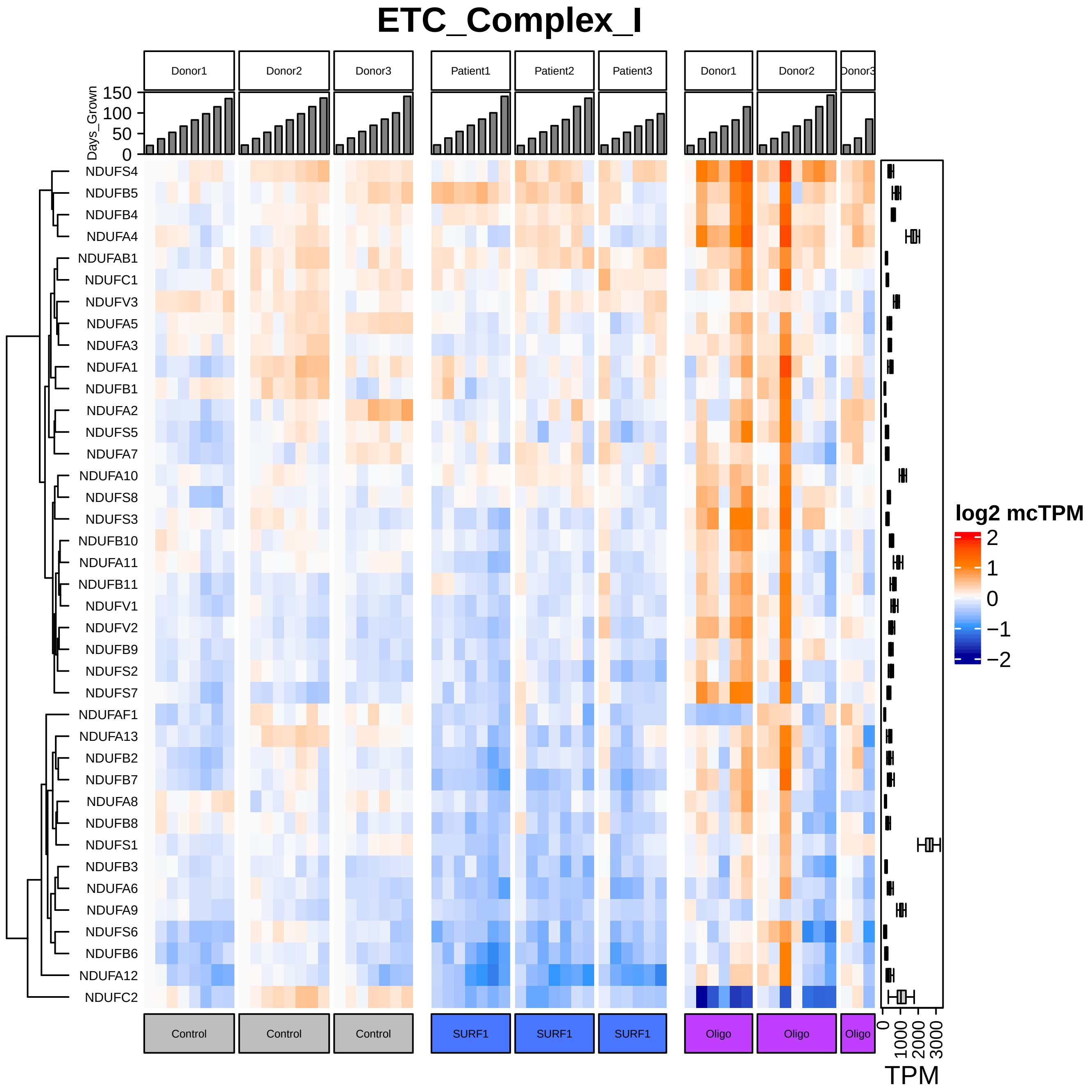

ETC_Complex_I
Donor1
Donor2
Donor3
Patient1
Patient2
Patient3
Donor1
Donor2
Donor3
Days_Grown
150
100
50
0
NDUFS4
NDUFB5
NDUFB4
NDUFA4
NDUFAB1
NDUFC1
NDUFV3
NDUFA5
NDUFA3
NDUFA1
NDUFB1
NDUFA2
NDUFS5
NDUFA7
NDUFA10
NDUFS8
NDUFS3
NDUFB10
NDUFA11
NDUFB11
NDUFV1
NDUFV2
NDUFB9
NDUFS2
NDUFS7
NDUFAF1
NDUFA13
NDUFB2
NDUFB7
NDUFA8
NDUFB8
NDUFS1
NDUFB3
NDUFA6
NDUFA9
NDUFS6
NDUFB6
NDUFA12
NDUFC2
log2 mcTPM
2
1
0
−1
−2
Control
Control
Control
SURF1
SURF1
SURF1
Oligo
Oligo
Oligo
0
1000
2000
3000
TPM

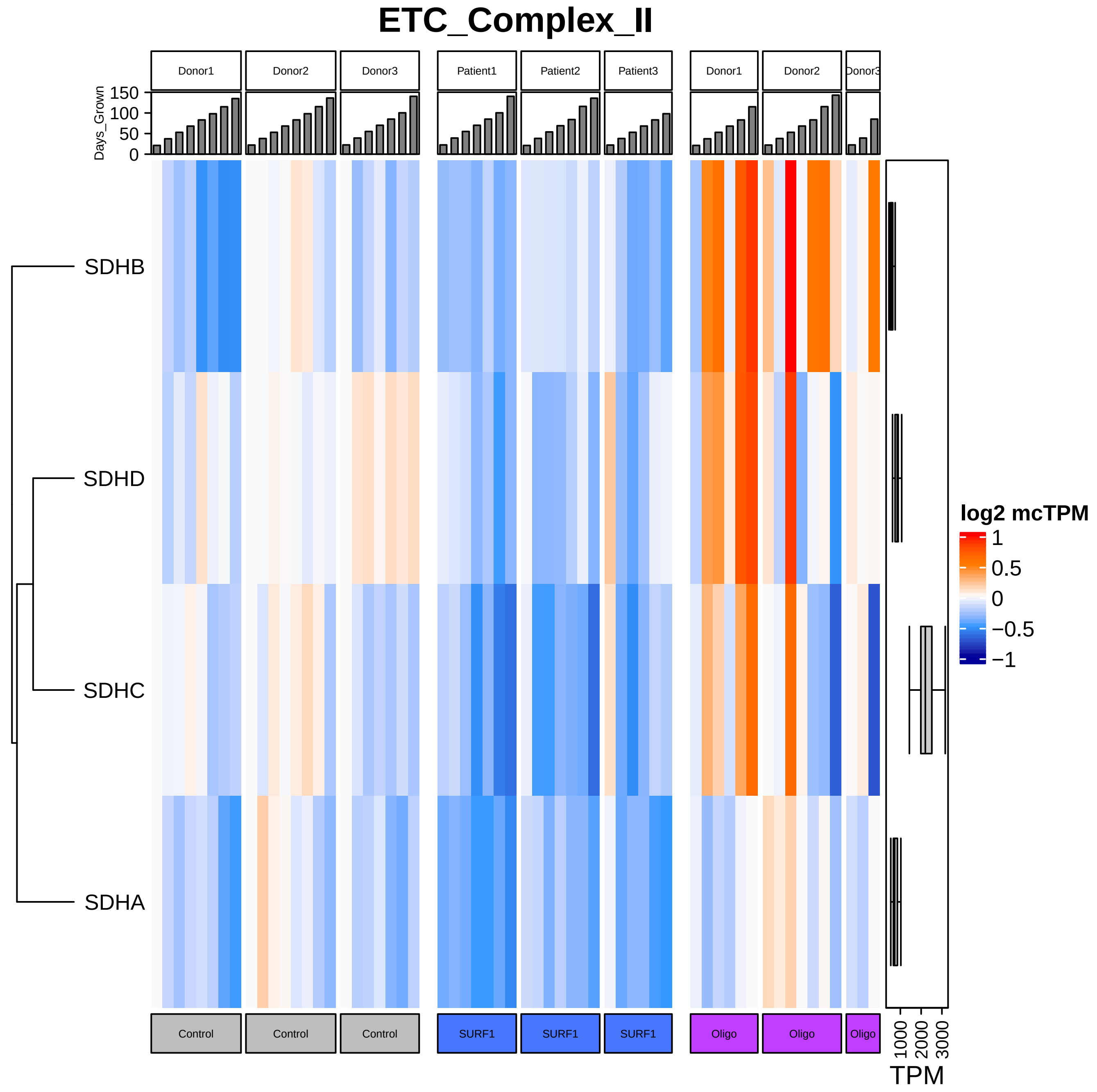

ETC_Complex_II
Donor1
Donor2
Donor3
Patient1
Patient2
Patient3
Donor1
Donor2
Donor3
Days_Grown
150
100
50
0
SDHB
SDHD
SDHC
SDHA
log2 mcTPM
1
0.5
0
−0.5
−1
Control
Control
Control
SURF1
SURF1
SURF1
Oligo
Oligo
Oligo
1000
2000
3000
TPM

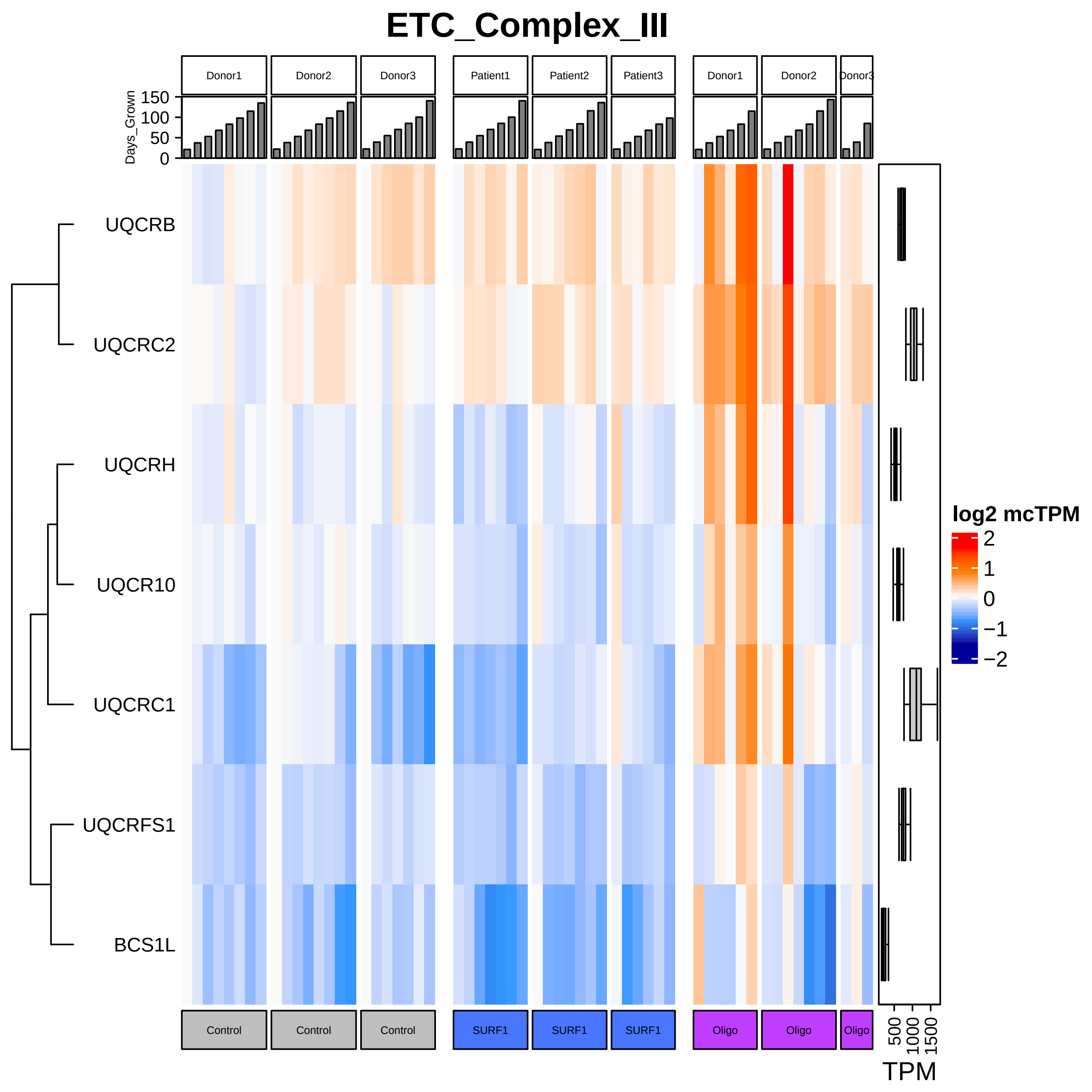

ETC_Complex_III
Donor1
Donor2
Donor3
Patient1
Patient2
Patient3
Donor1
Donor2
Donor3
Days_Grown
150
100
50
0
UQCRB
UQCRC2
UQCRH
UQCR10
UQCRC1
UQCRFS1
BCS1L
log2 mcTPM
2
1
0
−1
−2
Control
Control
Control
SURF1
SURF1
SURF1
Oligo
Oligo
Oligo
500
1000
1500
TPM

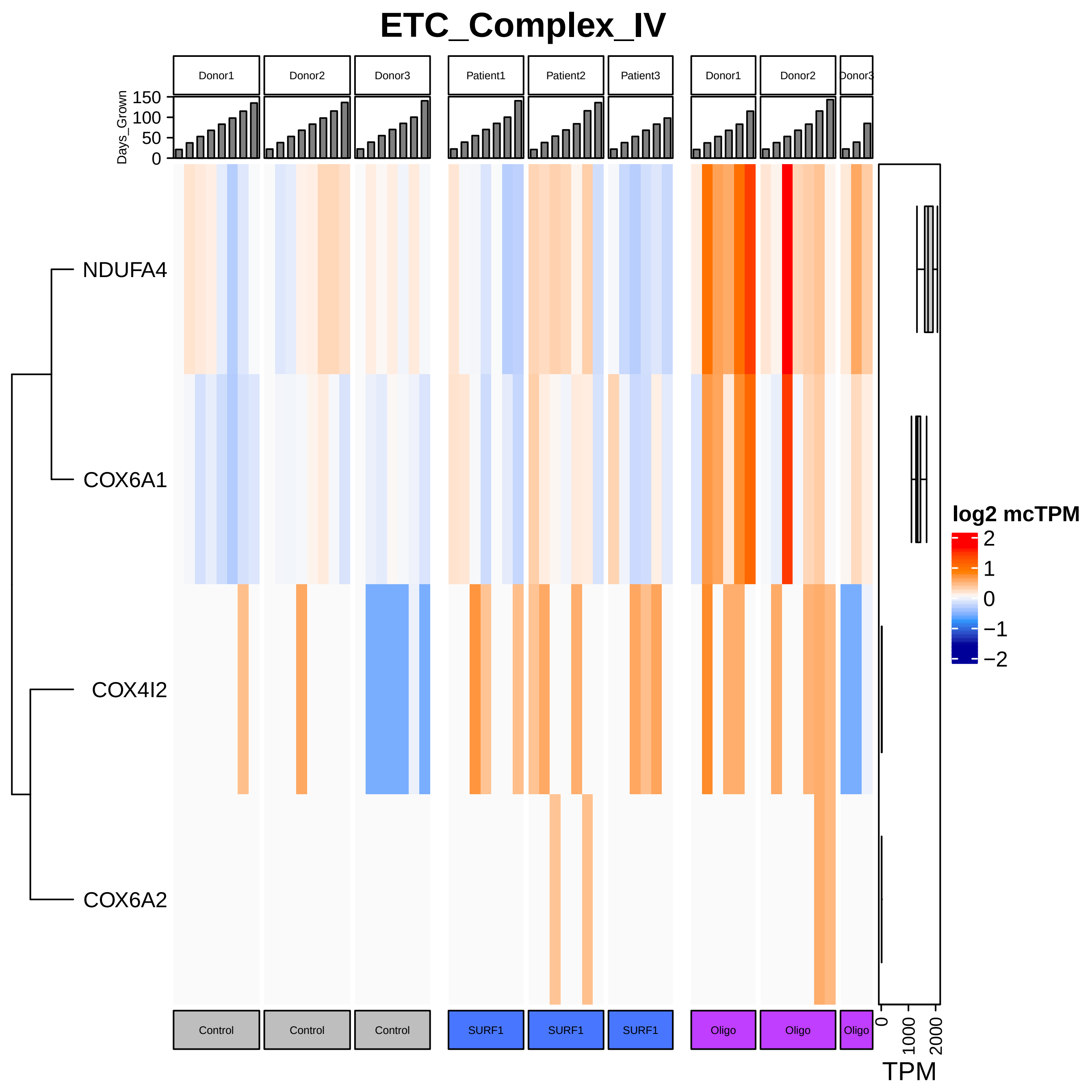

ETC_Complex_IV
Donor1
Donor2
Donor3
Patient1
Patient2
Patient3
Donor1
Donor2
Donor3
Days_Grown
150
100
50
0
NDUFA4
COX6A1
COX4I2
COX6A2
log2 mcTPM
2
1
0
−1
−2
Control
Control
Control
SURF1
SURF1
SURF1
Oligo
Oligo
Oligo
0
1000
2000
TPM

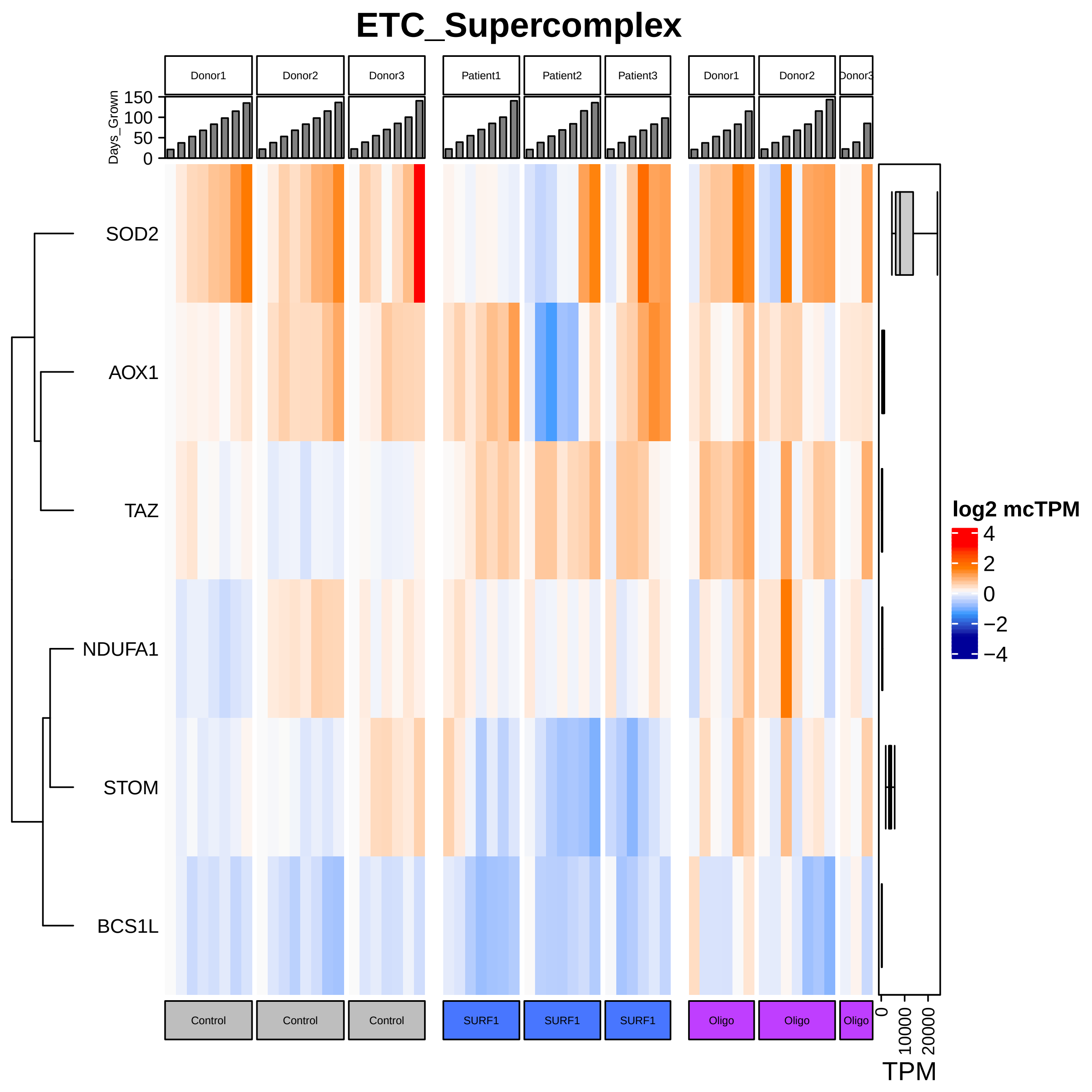
ETC_Supercomplex
Donor1
Donor2
Donor3
Patient1
Patient2
Patient3
Donor1
Donor2
Donor3
Days_Grown
150
100
50
0
SOD2
AOX1
TAZ
NDUFA1
STOM
BCS1L
log2 mcTPM
4
2
0
−2
−4
Control
Control
Control
SURF1
SURF1
SURF1
Oligo
Oligo
Oligo
0
10000
20000
TPM

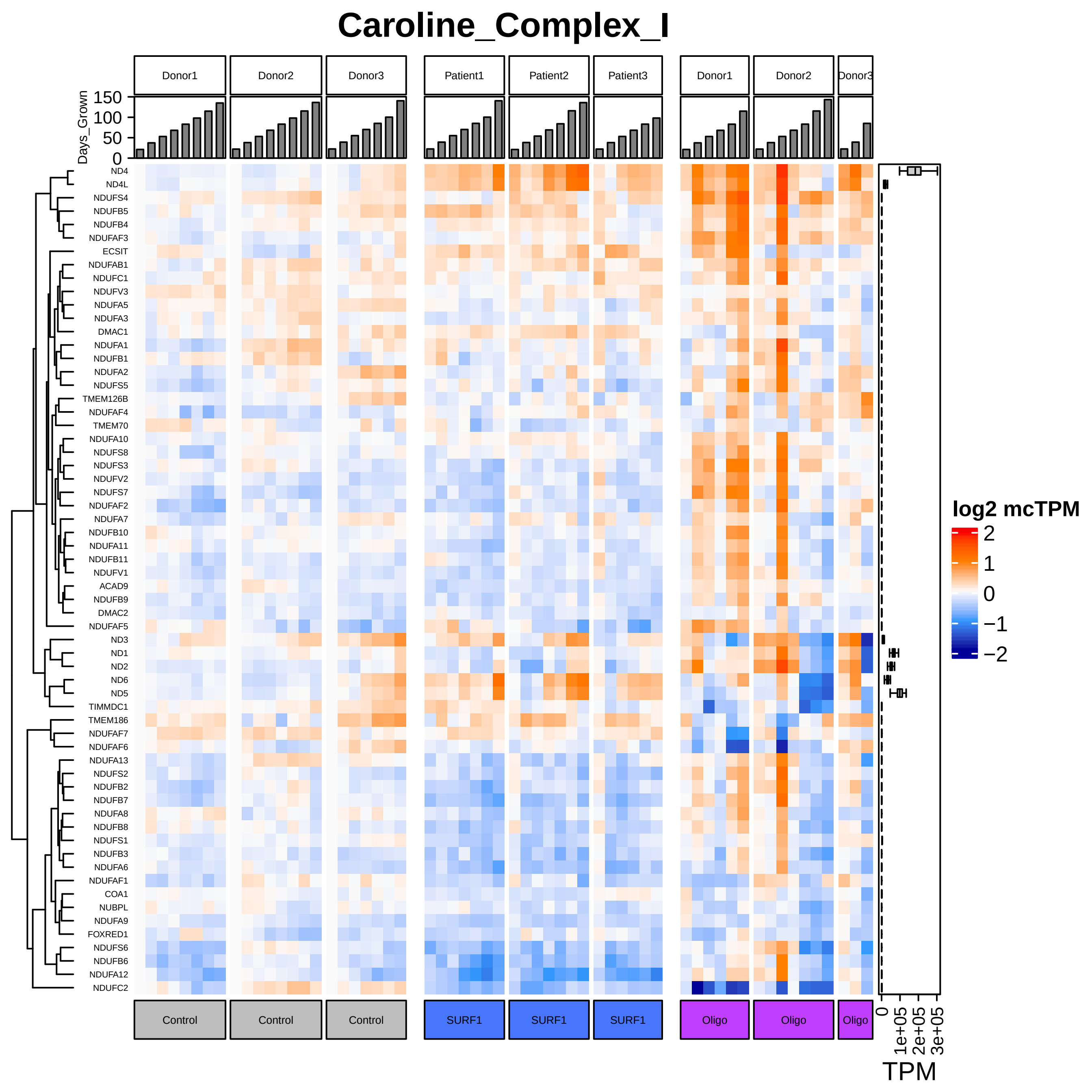
Caroline_Complex_I
Donor1
Donor2
Donor3
Patient1
Patient2
Patient3
Donor1
Donor2
Donor3
Days_Grown
150
100
50
0
ND4
ND4L
NDUFS4
NDUFB5
NDUFB4
NDUFAF3
ECSIT
NDUFAB1
NDUFC1
NDUFV3
NDUFA5
NDUFA3
DMAC1
NDUFA1
NDUFB1
NDUFA2
NDUFS5
TMEM126B
NDUFAF4
TMEM70
NDUFA10
NDUFS8
NDUFS3
NDUFV2
NDUFS7
NDUFAF2
NDUFA7
NDUFB10
NDUFA11
NDUFB11
NDUFV1
ACAD9
NDUFB9
DMAC2
NDUFAF5
ND3
ND1
ND2
ND6
ND5
TIMMDC1
TMEM186
NDUFAF7
NDUFAF6
NDUFA13
NDUFS2
NDUFB2
NDUFB7
NDUFA8
NDUFB8
NDUFS1
NDUFB3
NDUFA6
NDUFAF1
COA1
NUBPL
NDUFA9
FOXRED1
NDUFS6
NDUFB6
NDUFA12
NDUFC2
log2 mcTPM
2
1
0
−1
−2
Control
Control
Control
SURF1
SURF1
SURF1
Oligo
Oligo
Oligo
0
1e+05
2e+05
3e+05
TPM

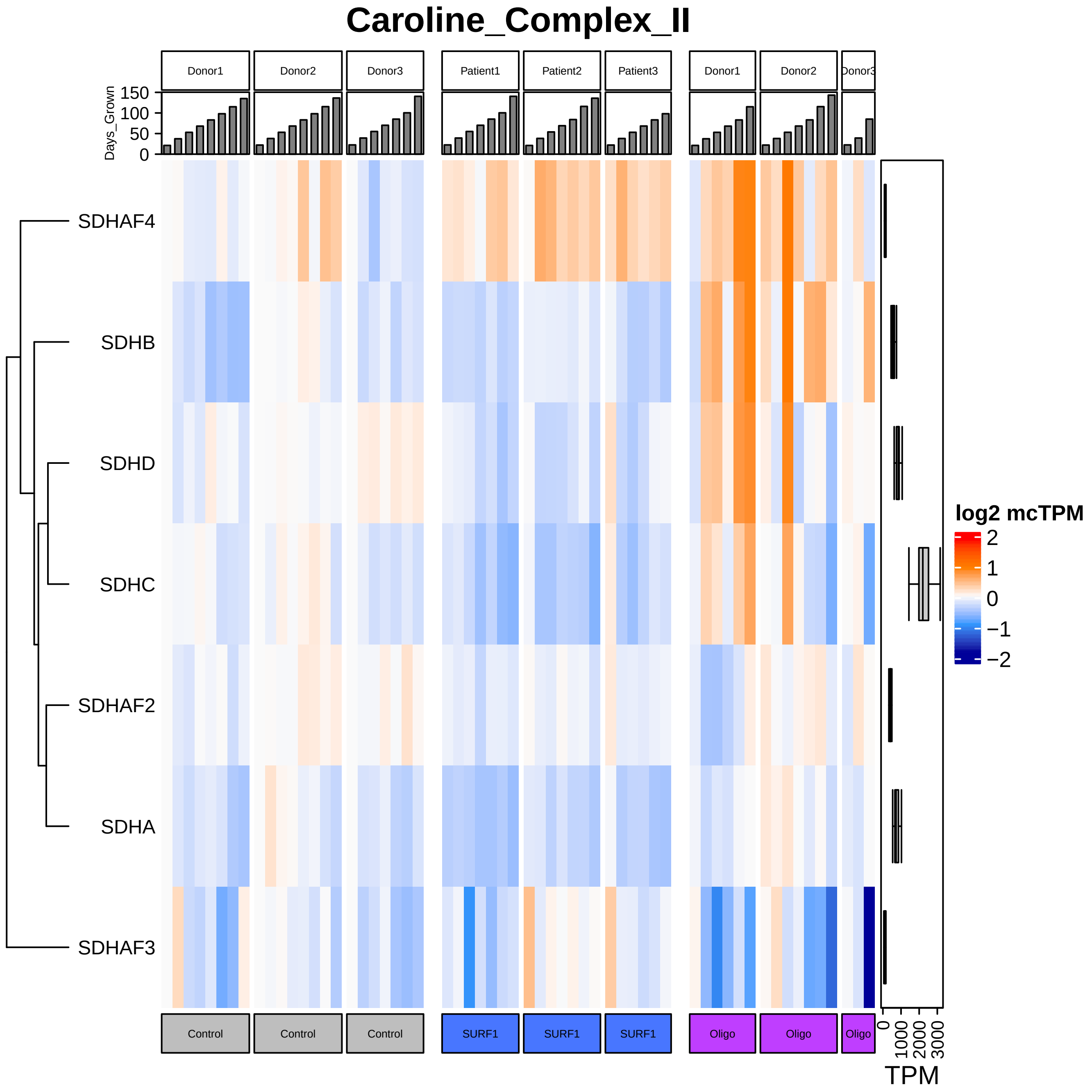

Caroline_Complex_II
Donor1
Donor2
Donor3
Patient1
Patient2
Patient3
Donor1
Donor2
Donor3
Days_Grown
150
100
50
0
SDHAF4
SDHB
SDHD
SDHC
SDHAF2
SDHA
SDHAF3
log2 mcTPM
2
1
0
−1
−2
Control
Control
Control
SURF1
SURF1
SURF1
Oligo
Oligo
Oligo
0
1000
2000
3000
TPM

### Caroline_Complex_III

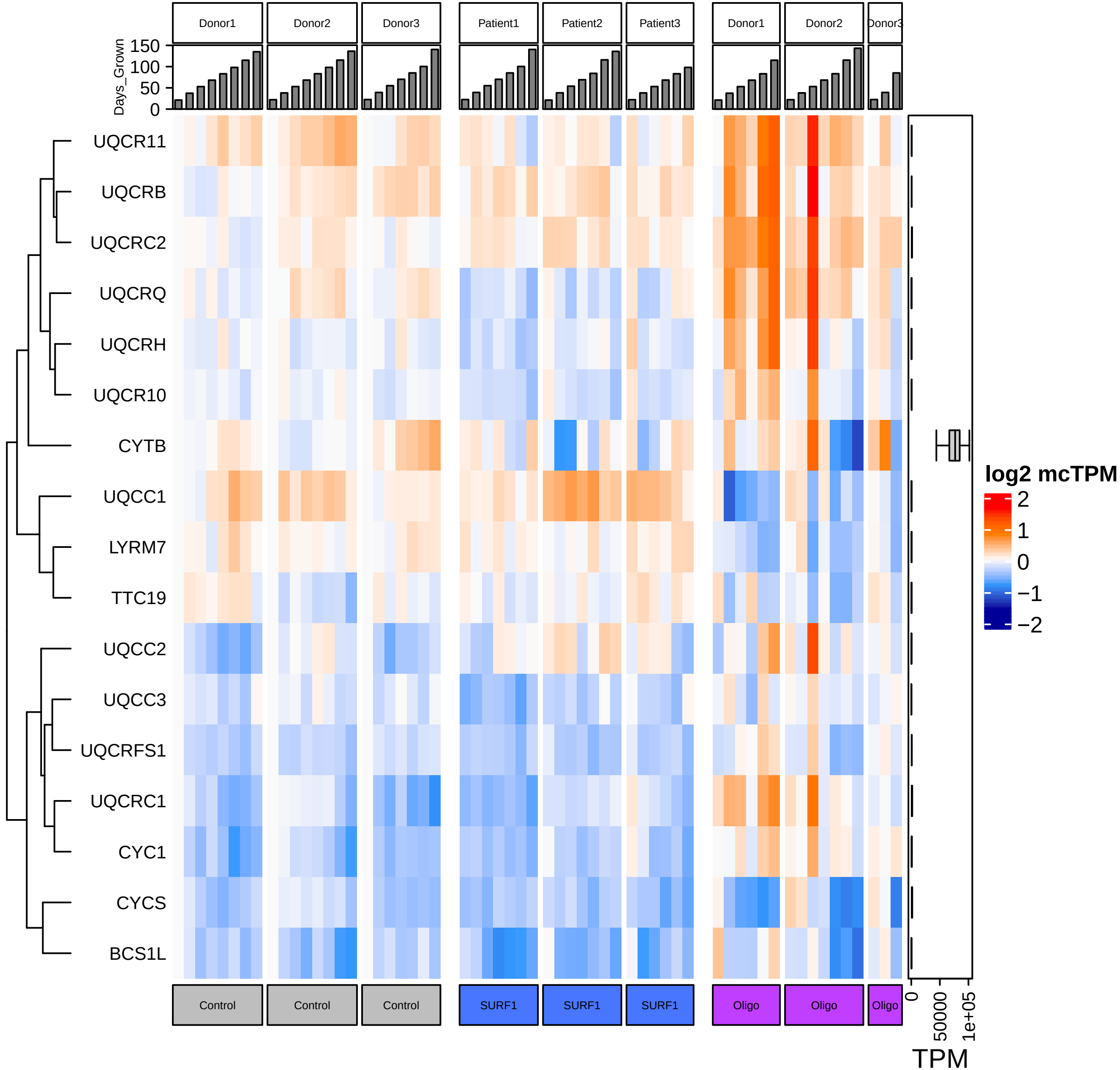

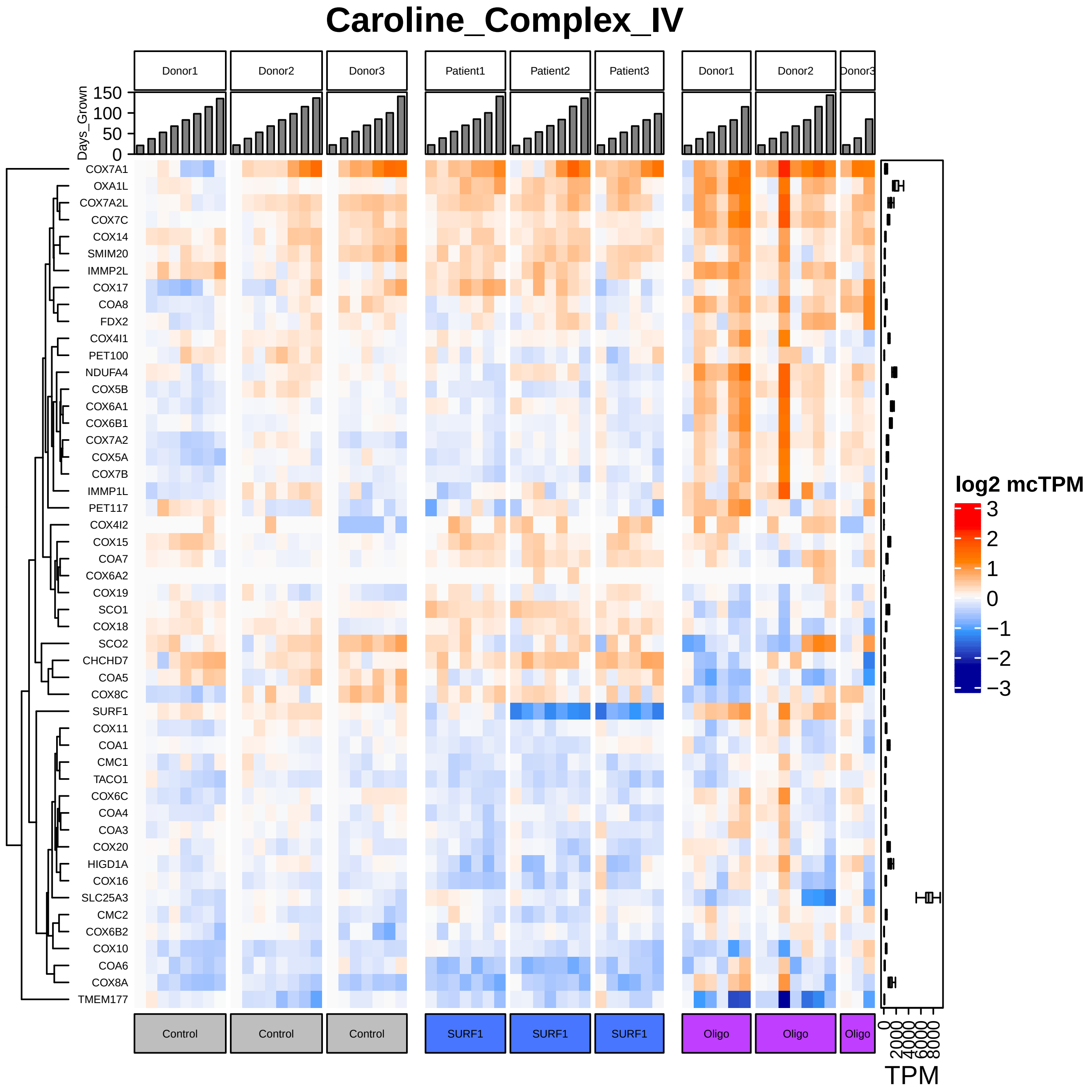

Caroline_Complex_IV
Donor1
Donor2
Donor3
Patient1
Patient2
Patient3
Donor1
Donor2
Donor3
Days_Grown
150
100
50
0
COX7A1
OXA1L
COX7A2L
COX7C
COX14
SMIM20
IMMP2L
COX17
COA8
FDX2
COX4I1
PET100
NDUFA4
COX5B
COX6A1
COX6B1
COX7A2
COX5A
COX7B
IMMP1L
PET117
COX4I2
COX15
COA7
COX6A2
COX19
SCO1
COX18
SCO2
CHCHD7
COA5
COX8C
SURF1
COX11
COA1
CMC1
TACO1
COX6C
COA4
COA3
COX20
HIGD1A
COX16
SLC25A3
CMC2
COX6B2
COX10
COA6
COX8A
TMEM177
Control
Control
Control
SURF1
SURF1
SURF1
Oligo
Oligo
Oligo
log2 mcTPM
3
2
1
0
−1
−2
−3
0
2000
4000
6000
8000
TPM

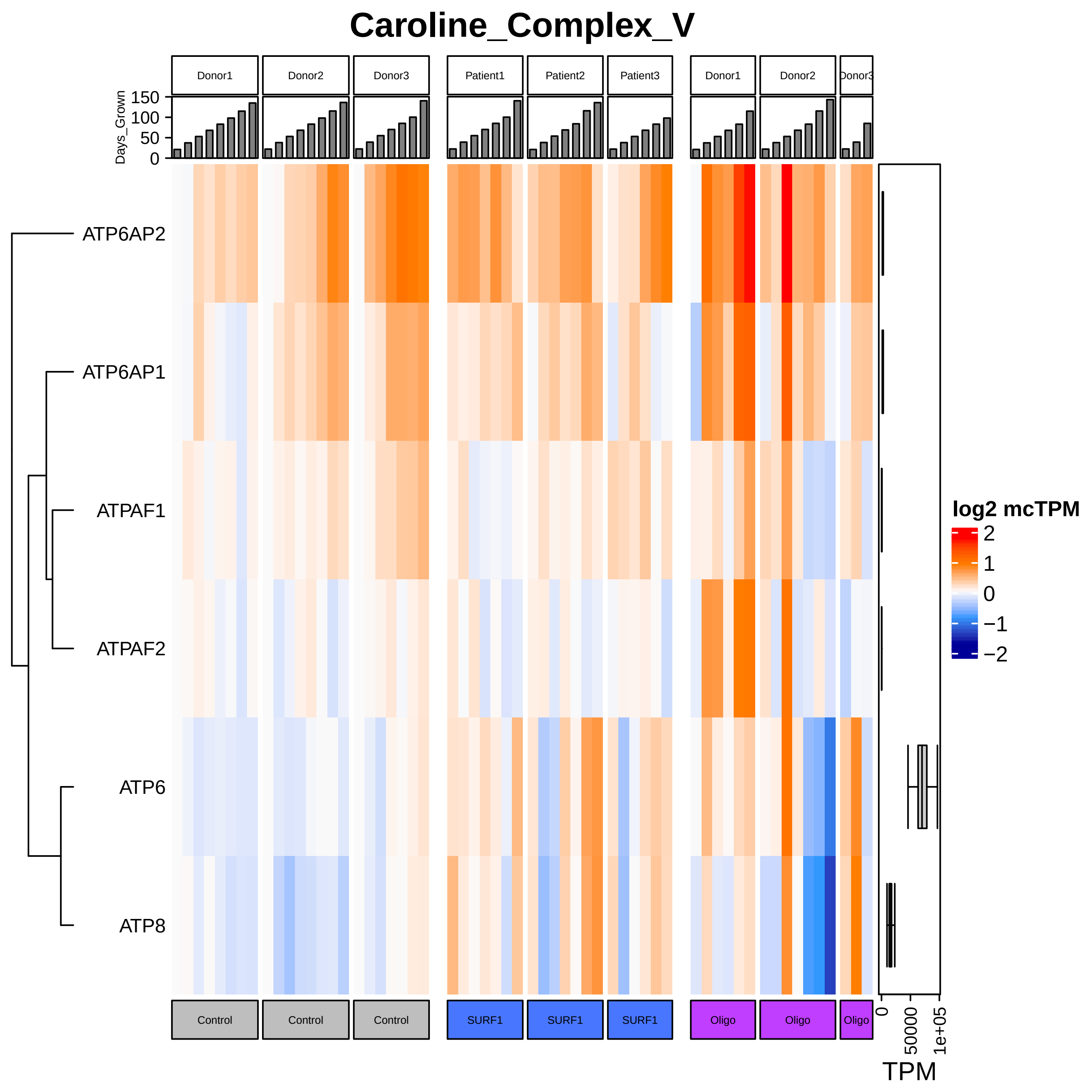

Caroline_Complex_V
Donor1
Donor2
Donor3
Patient1
Patient2
Patient3
Donor1
Donor2
Donor3
Days_Grown
150
100
50
0
ATP6AP2
ATP6AP1
ATPAF1
ATPAF2
ATP6
ATP8
log2 mcTPM
2
1
0
−1
−2
Control
Control
Control
SURF1
SURF1
SURF1
Oligo
Oligo
Oligo
0
50000
1e+05
TPM

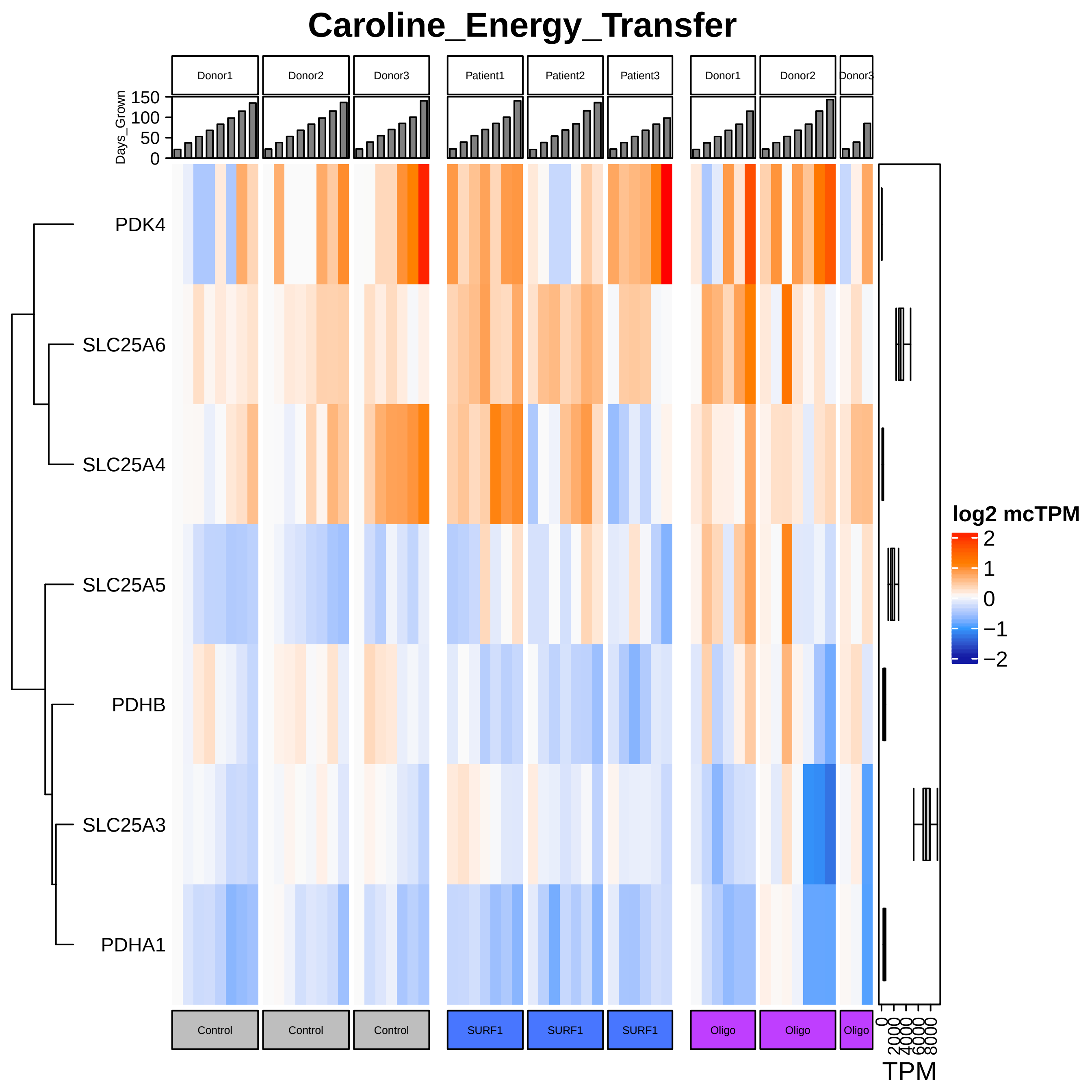

Caroline_Energy_Transfer
Donor1
Donor2
Donor3
Patient1
Patient2
Patient3
Donor1
Donor2
Donor3
Days_Grown
150
100
50
0
PDK4
SLC25A6
SLC25A4
SLC25A5
PDHB
SLC25A3
PDHA1
log2 mcTPM
2
1
0
−1
−2
Control
Control
Control
SURF1
SURF1
SURF1
Oligo
Oligo
Oligo
0
2000
4000
6000
8000
TPM

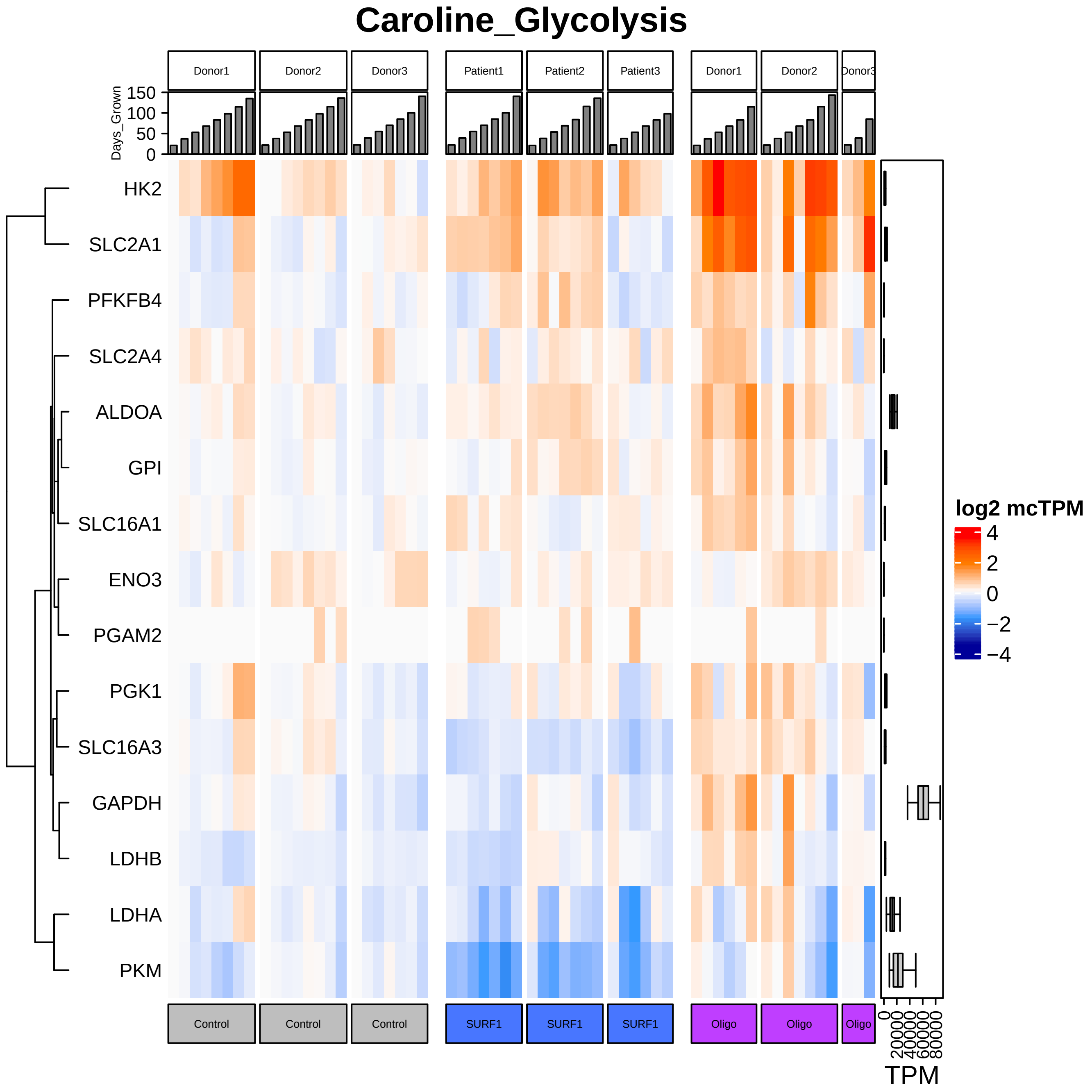

Caroline_Glycolysis
Donor1
Donor2
Donor3
Patient1
Patient2
Patient3
Donor1
Donor2
Donor3
Days_Grown
150
100
50
0
HK2
SLC2A1
PFKFB4
SLC2A4
ALDOA
GPI
SLC16A1
ENO3
PGAM2
PGK1
SLC16A3
GAPDH
LDHB
LDHA
PKM
Control
Control
Control
SURF1
SURF1
SURF1
Oligo
Oligo
Oligo
0
20000
40000
60000
80000
TPM
log2 mcTPM
4
2
0
−2
−4

### Caroline_Metabolic_Sensing

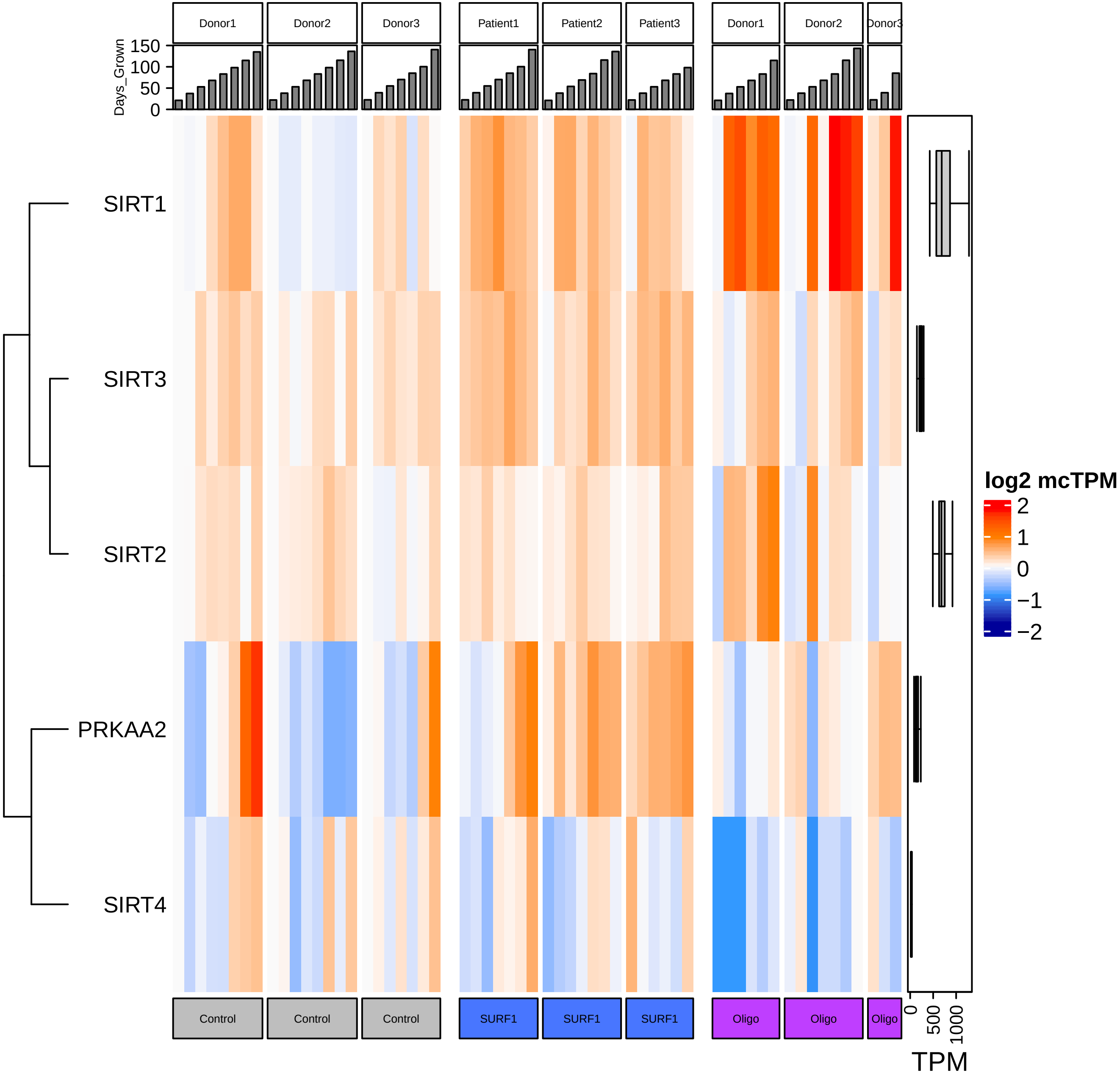

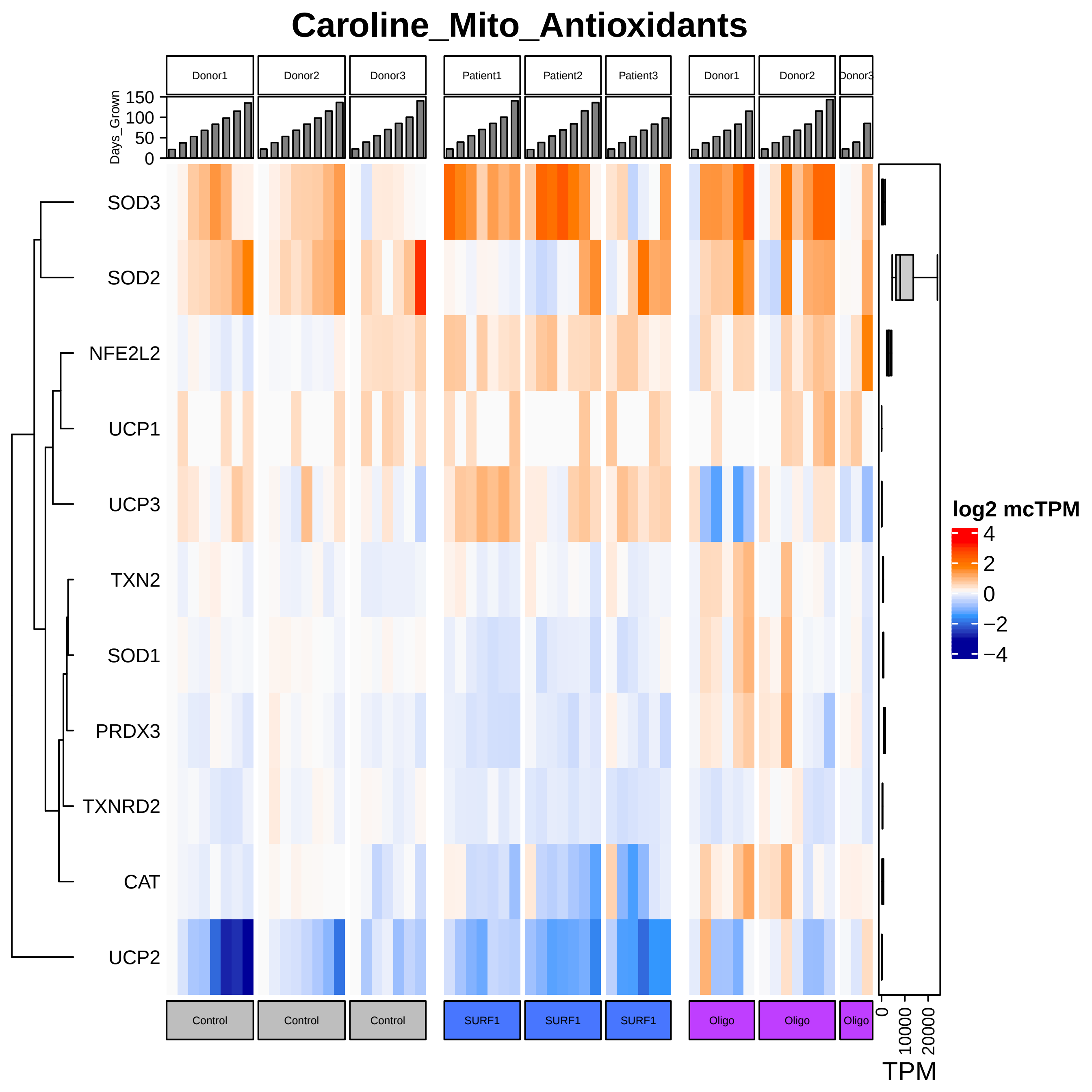

Caroline_Mito_Antioxidants
Donor1
Donor2
Donor3
Patient1
Patient2
Patient3
Donor1
Donor2
Donor3
Days_Grown
150
100
50
0
SOD3
SOD2
NFE2L2
UCP1
UCP3
TXN2
SOD1
PRDX3
TXNRD2
CAT
UCP2
log2 mcTPM
4
2
0
−2
−4
Control
Control
Control
SURF1
SURF1
SURF1
Oligo
Oligo
Oligo
0
10000
20000
TPM

### Caroline_Mito_Axonal_Transport

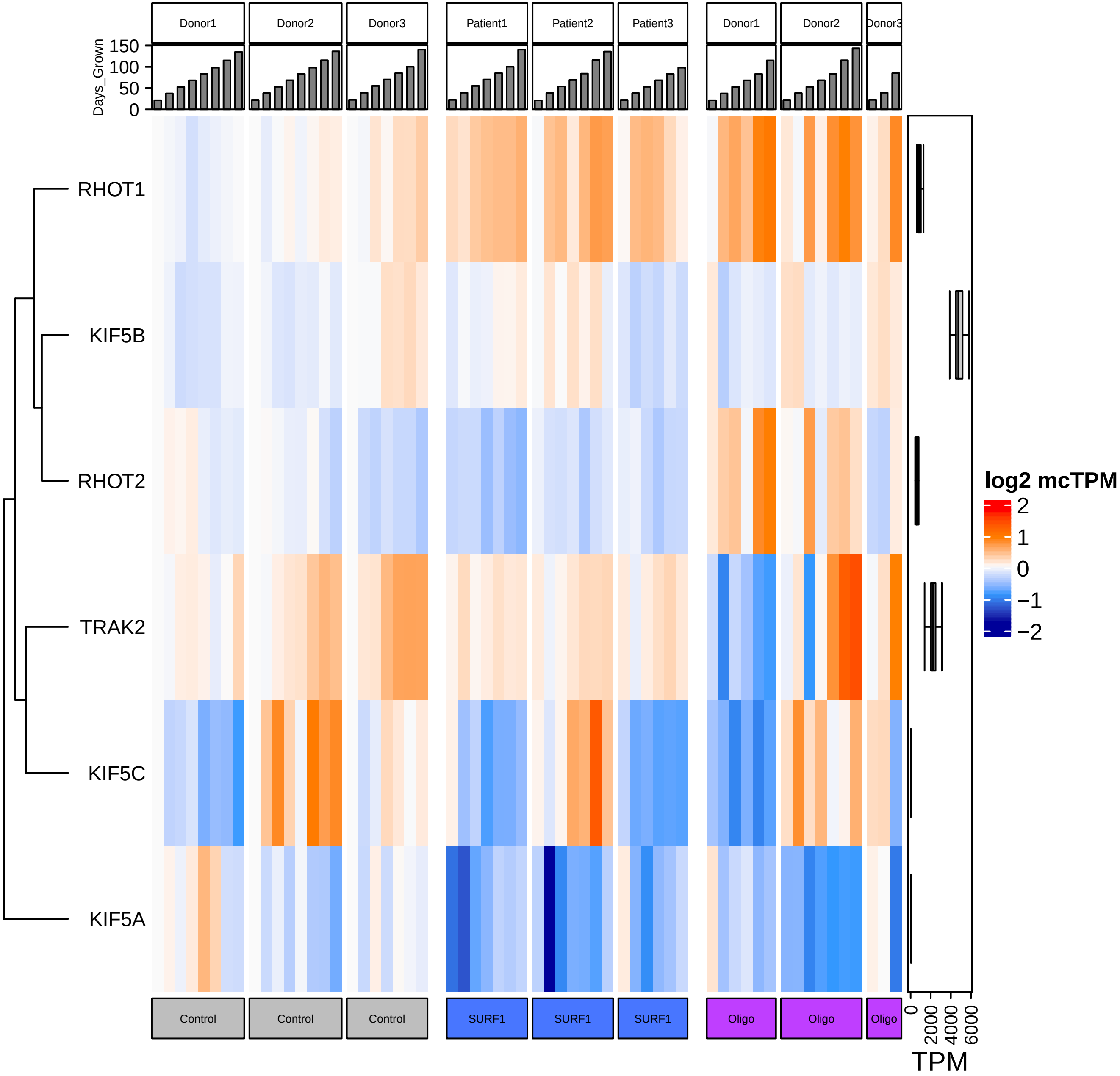

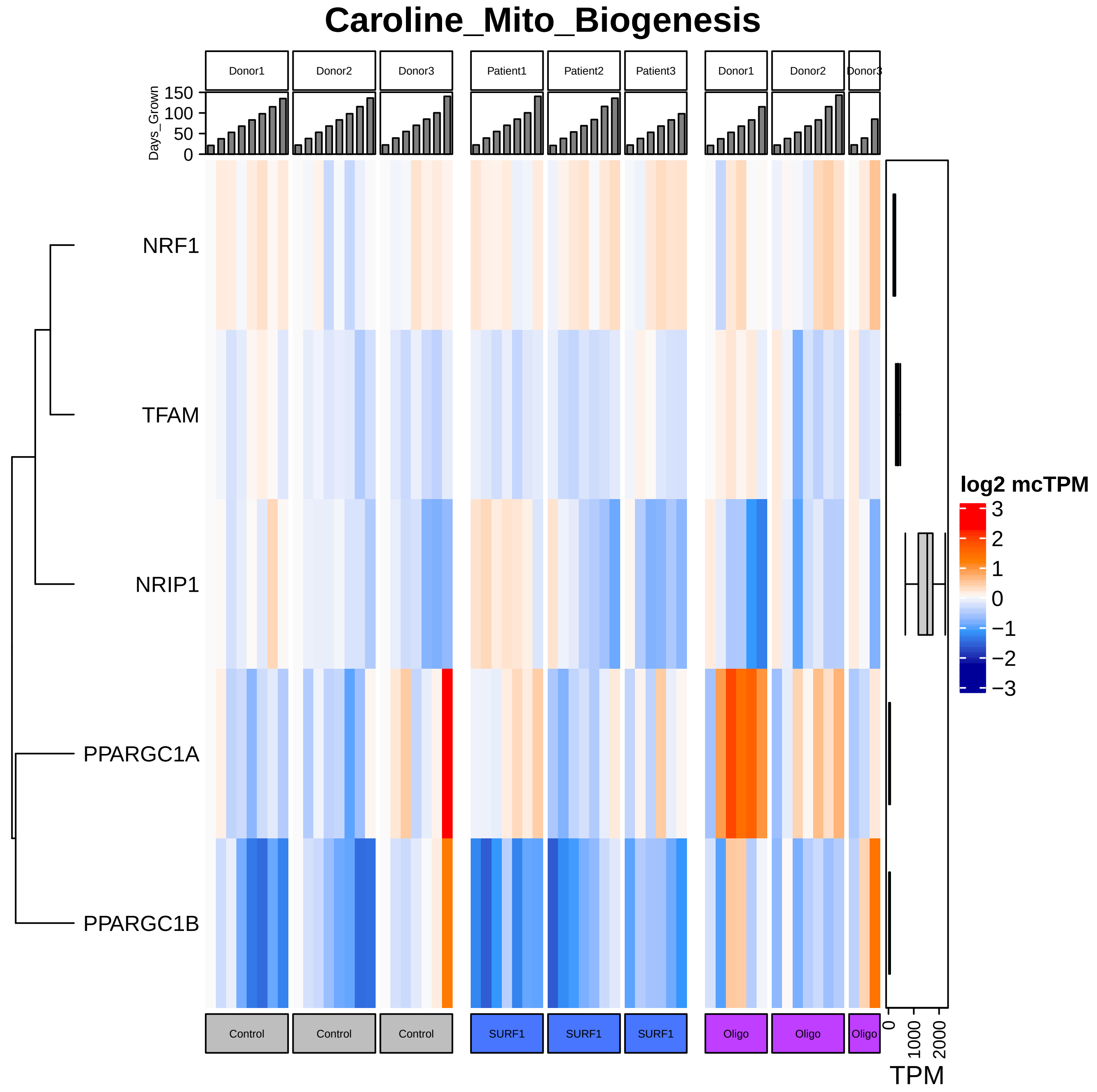
Caroline_Mito_Biogenesis
Donor1
Donor2
Donor3
Patient1
Patient2
Patient3
Donor1
Donor2
Donor3
Days_Grown
150
100
50
0
NRF1
TFAM
NRIP1
PPARGC1A
PPARGC1B
log2 mcTPM
3
2
1
0
−1
−2
−3
Control
Control
Control
SURF1
SURF1
SURF1
Oligo
Oligo
Oligo
0
1000
2000
TPM

### Caroline_Mito_Calcium_Handling

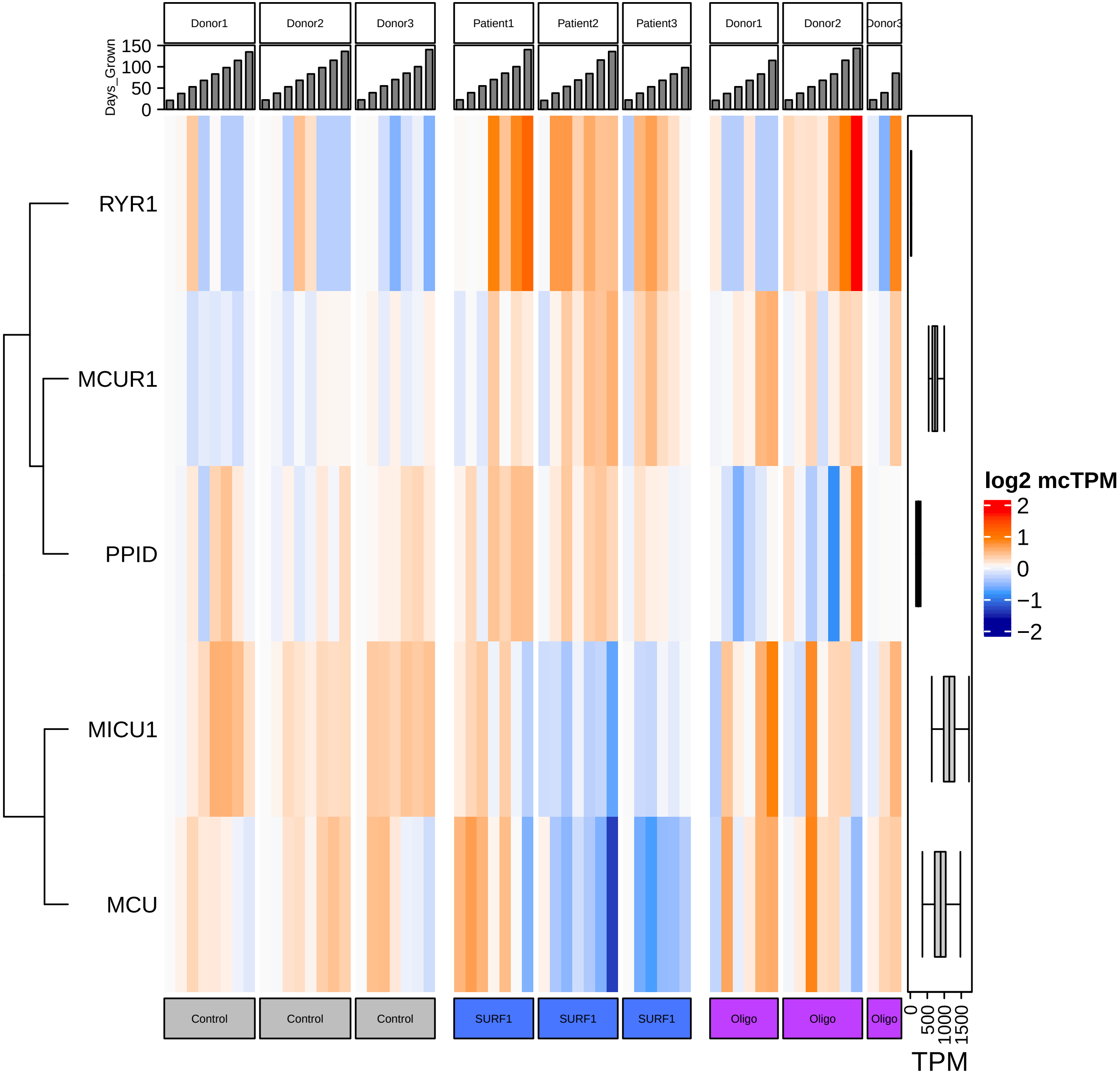

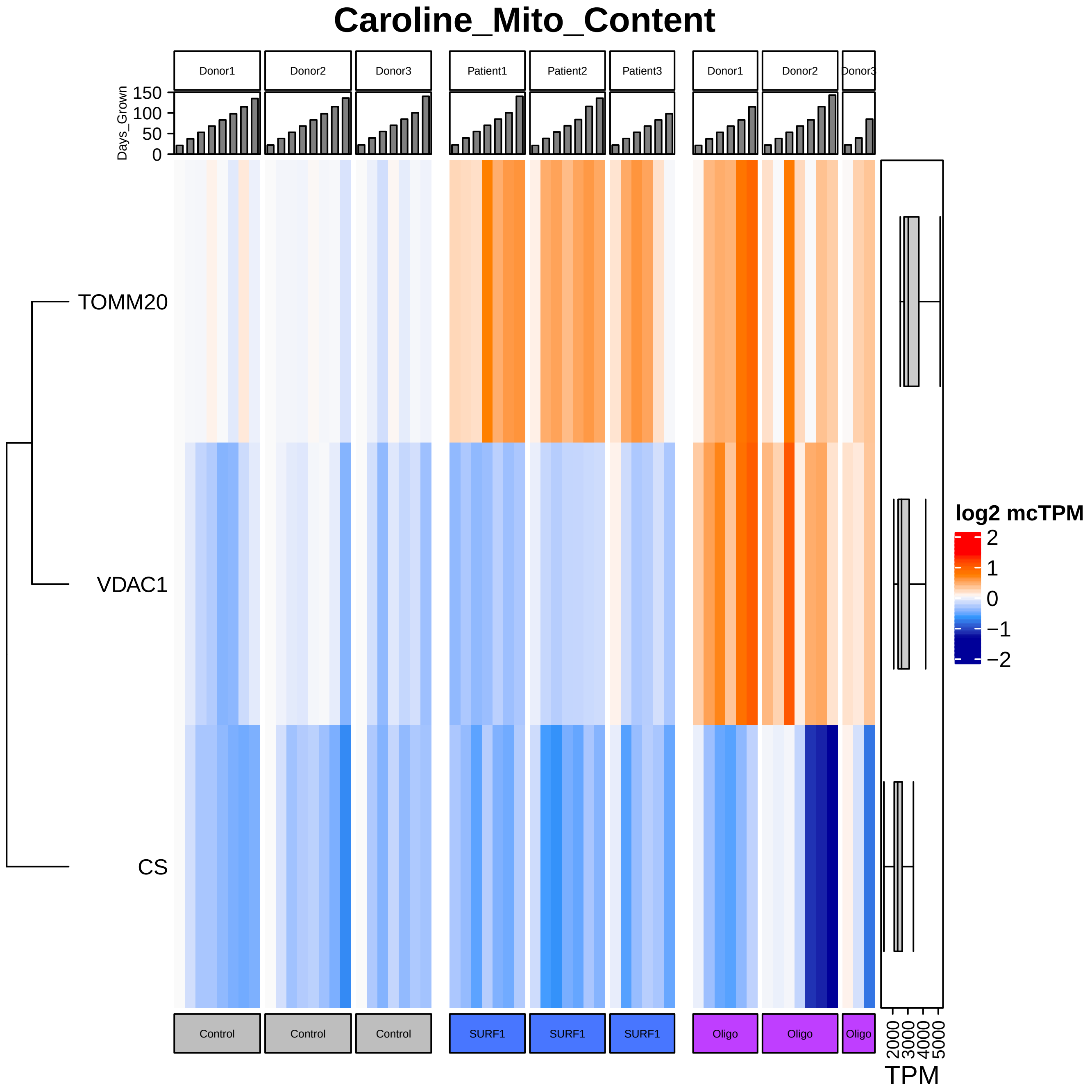

Caroline_Mito_Content
Donor1
Donor2
Donor3
Patient1
Patient2
Patient3
Donor1
Donor2
Donor3
Days_Grown
150
100
50
0
TOMM20
VDAC1
CS
log2 mcTPM
2
1
0
−1
−2
Control
Control
Control
SURF1
SURF1
SURF1
Oligo
Oligo
Oligo
2000
3000
4000
5000
TPM

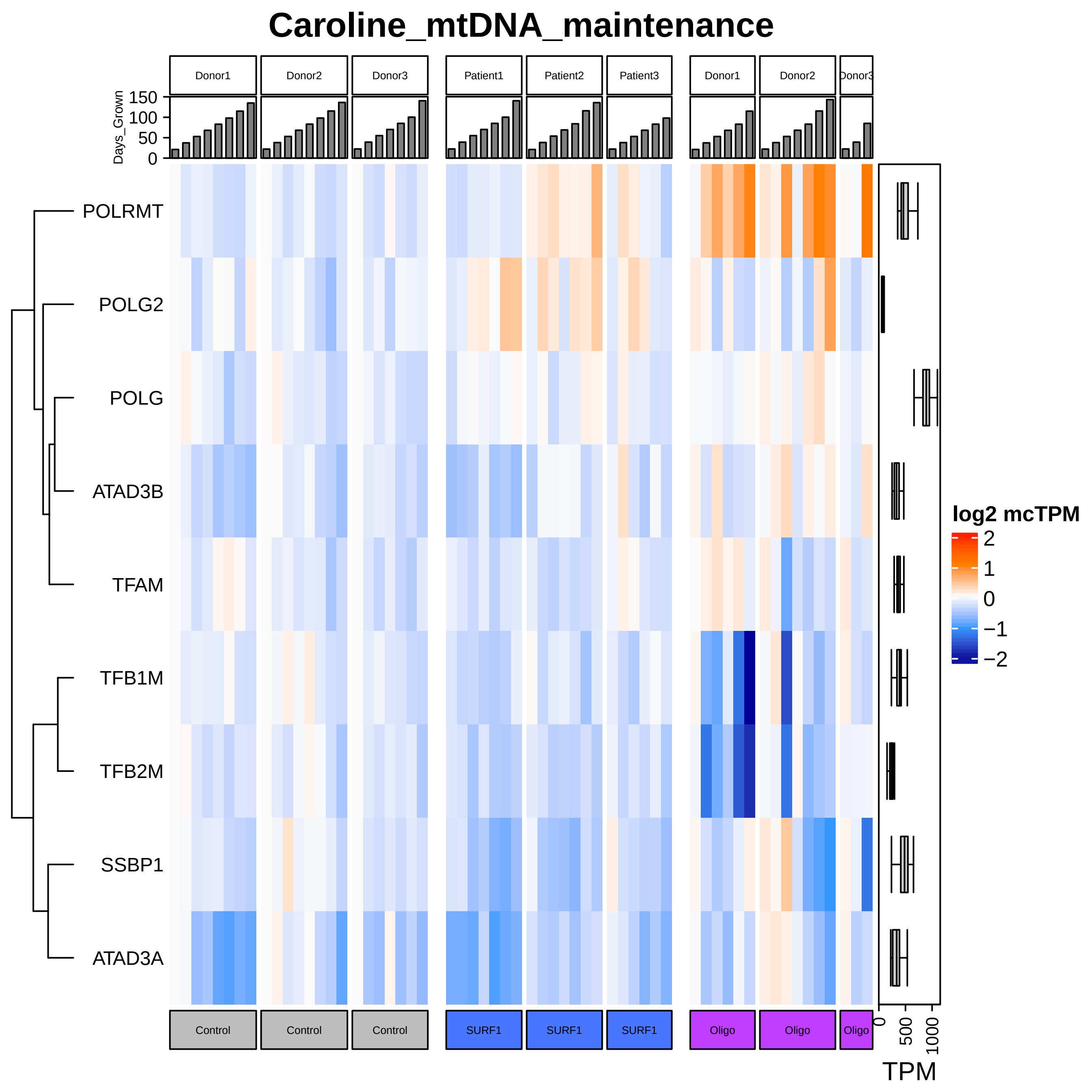
Caroline_mtDNA_maintenance
Donor1
Donor2
Donor3
Patient1
Patient2
Patient3
Donor1
Donor2
Donor3
Days_Grown
150
100
50
0
POLRMT
POLG2
POLG
ATAD3B
TFAM
TFB1M
TFB2M
SSBP1
ATAD3A
log2 mcTPM
2
1
0
−1
−2
Control
Control
Control
SURF1
SURF1
SURF1
Oligo
Oligo
Oligo
0
500
1000
TPM

### Caroline_Mito_Dynamics

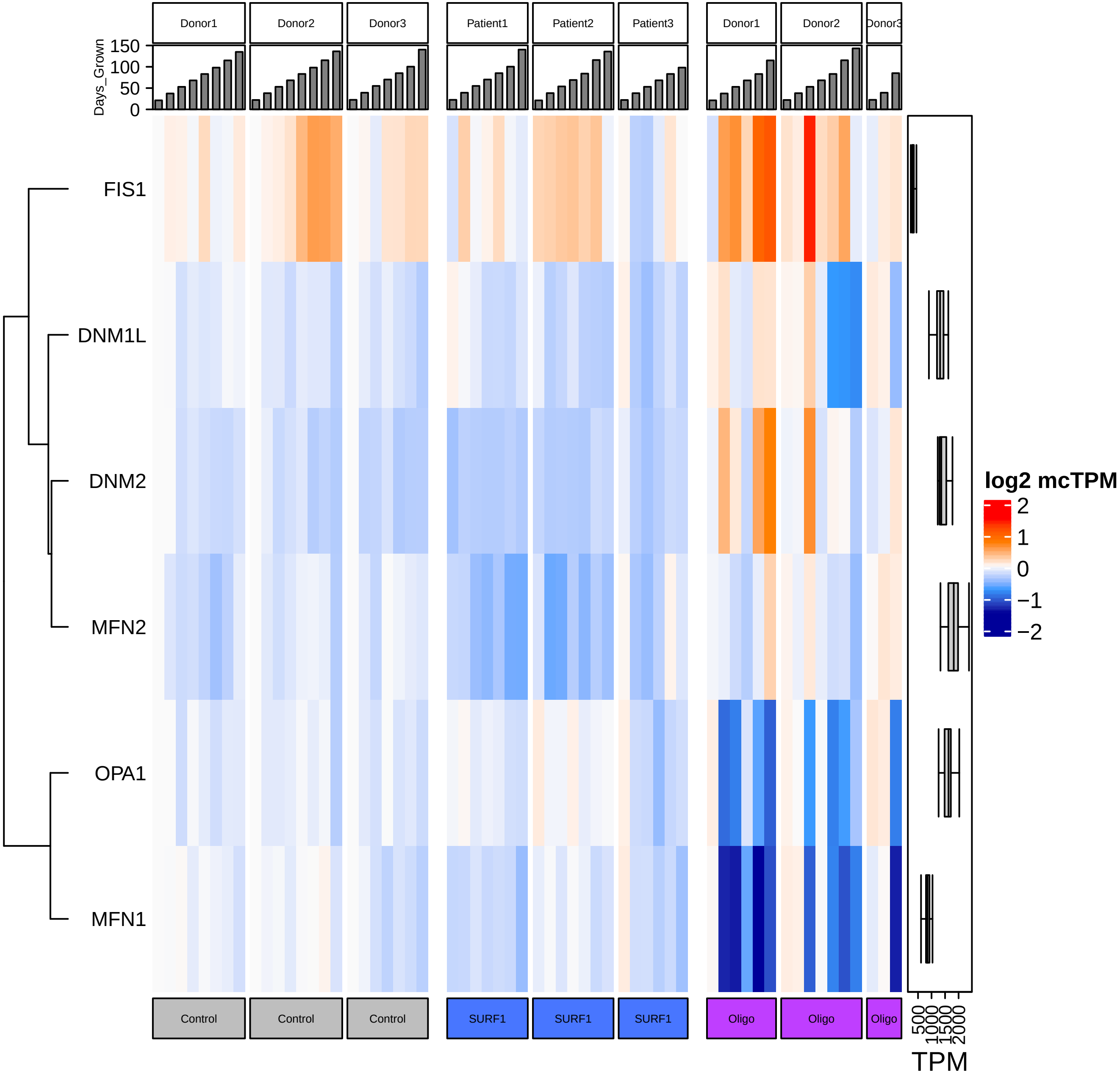

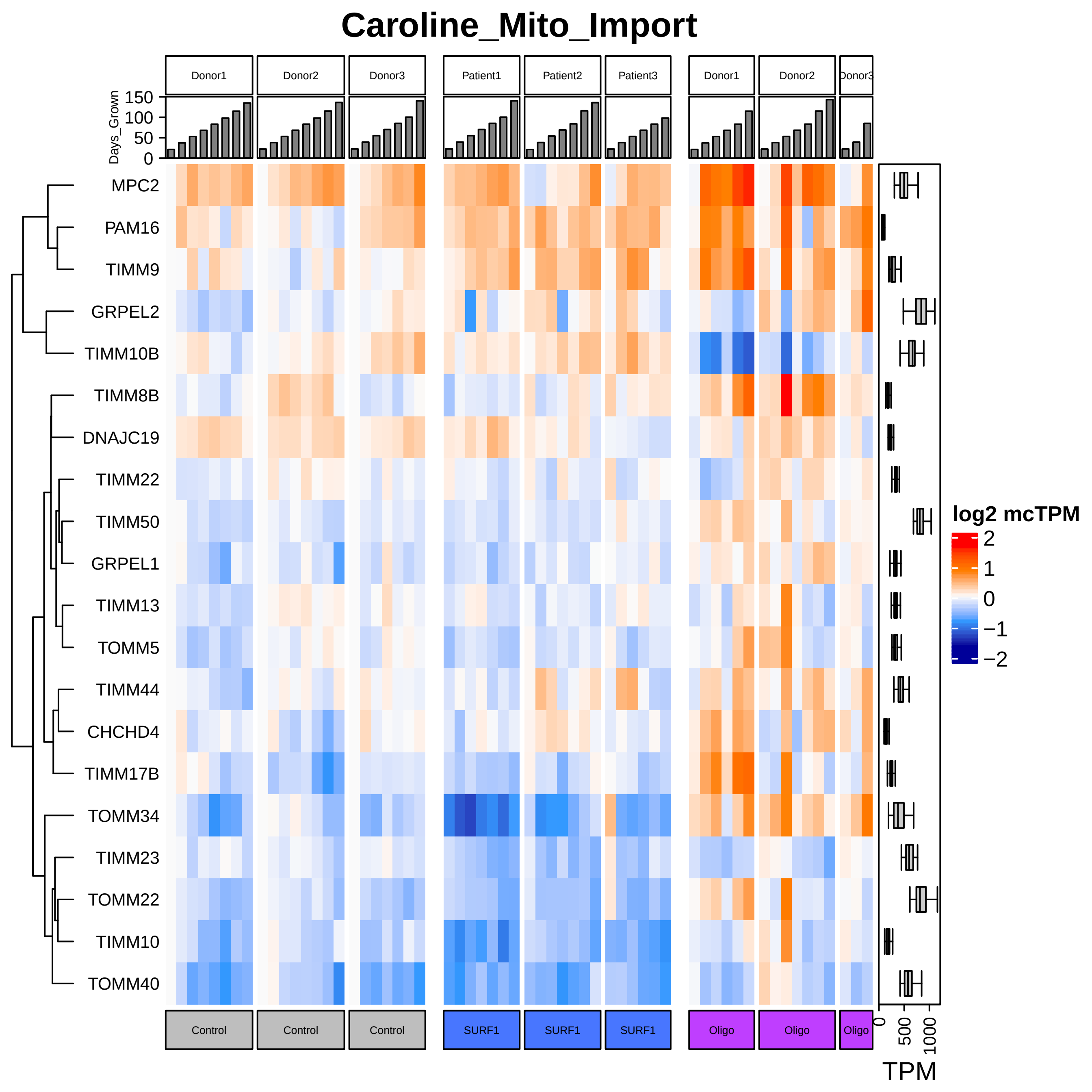

Caroline_Mito_Import
Donor1
Donor2
Donor3
Patient1
Patient2
Patient3
Donor1
Donor2
Donor3
Days_Grown
150
100
50
0
MPC2
PAM16
TIMM9
GRPEL2
TIMM10B
TIMM8B
DNAJC19
TIMM22
TIMM50
GRPEL1
TIMM13
TOMM5
TIMM44
CHCHD4
TIMM17B
TOMM34
TIMM23
TOMM22
TIMM10
TOMM40
log2 mcTPM
2
1
0
−1
−2
Control
Control
Control
SURF1
SURF1
SURF1
Oligo
Oligo
Oligo
0
500
1000
TPM

### Caroline_Mito_Ribosome

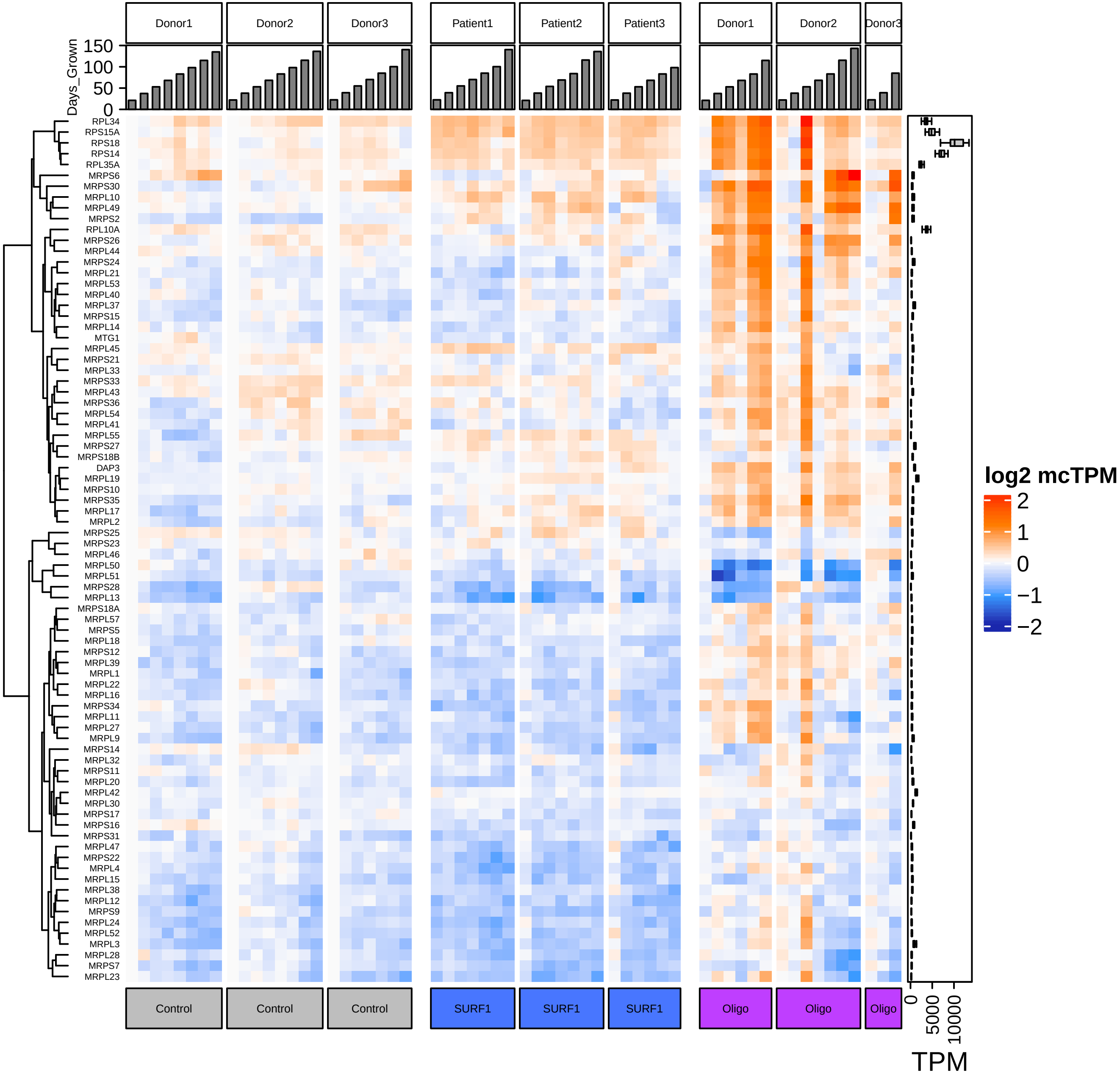

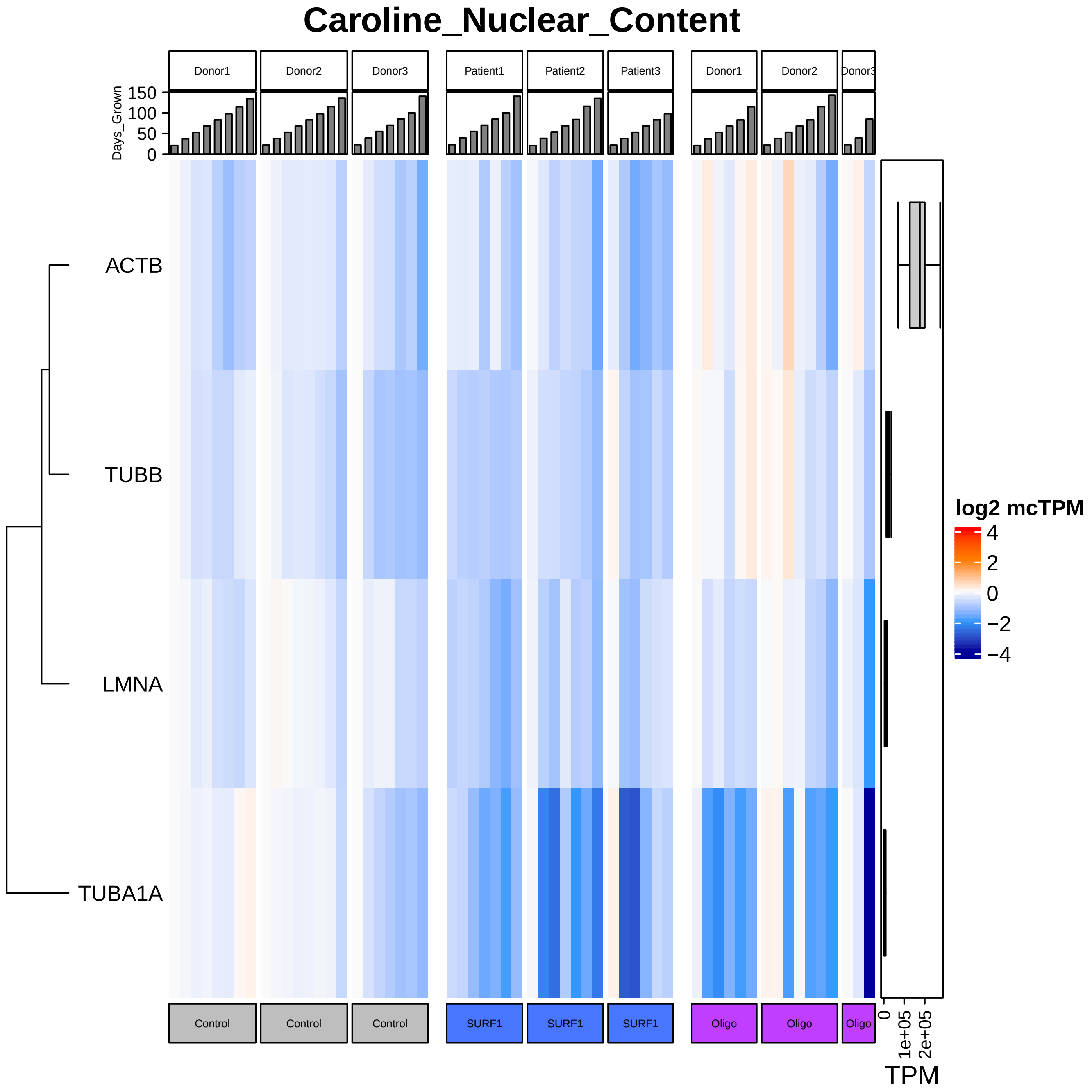

Caroline_Nuclear_Content
Donor1
Donor2
Donor3
Patient1
Patient2
Patient3
Donor1
Donor2
Donor3
Days_Grown
150
100
50
0
ACTB
TUBB
LMNA
TUBA1A
log2 mcTPM
4
2
0
−2
−4
Control
Control
Control
SURF1
SURF1
SURF1
Oligo
Oligo
Oligo
0
1e+05
2e+05
TPM

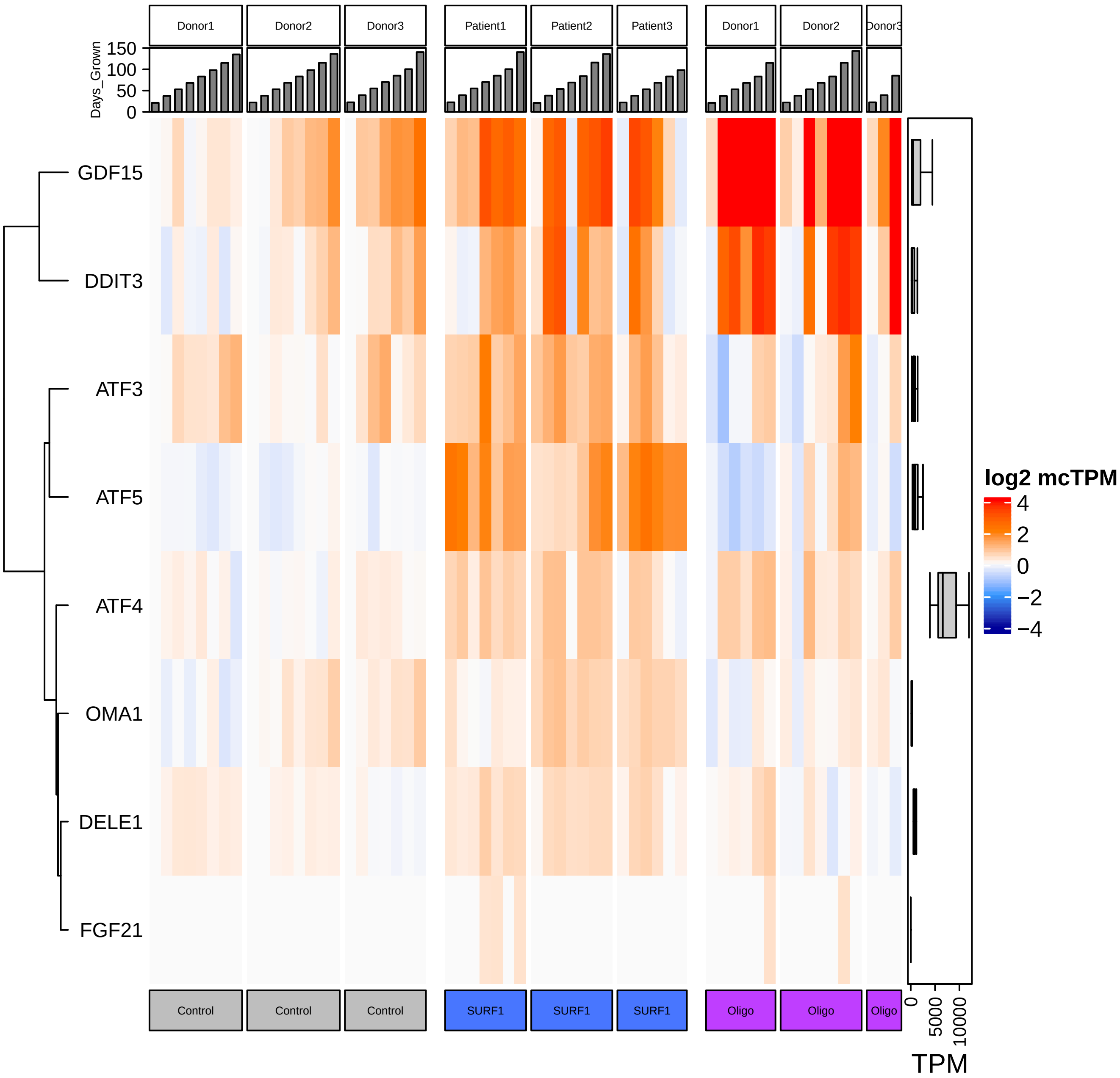

Integrated_Stress_Response
Donor1
Donor2
Donor3
Patient1
Patient2
Patient3
Donor1
Donor2
Donor3
Days_Grown
150
100
50
0
GDF15
DDIT3
ATF3
ATF5
ATF4
OMA1
DELE1
FGF21
log2 mcTPM
4
2
0
−2
−4
Control
Control
Control
SURF1
SURF1
SURF1
Oligo
Oligo
Oligo
0
5000
10000
TPM

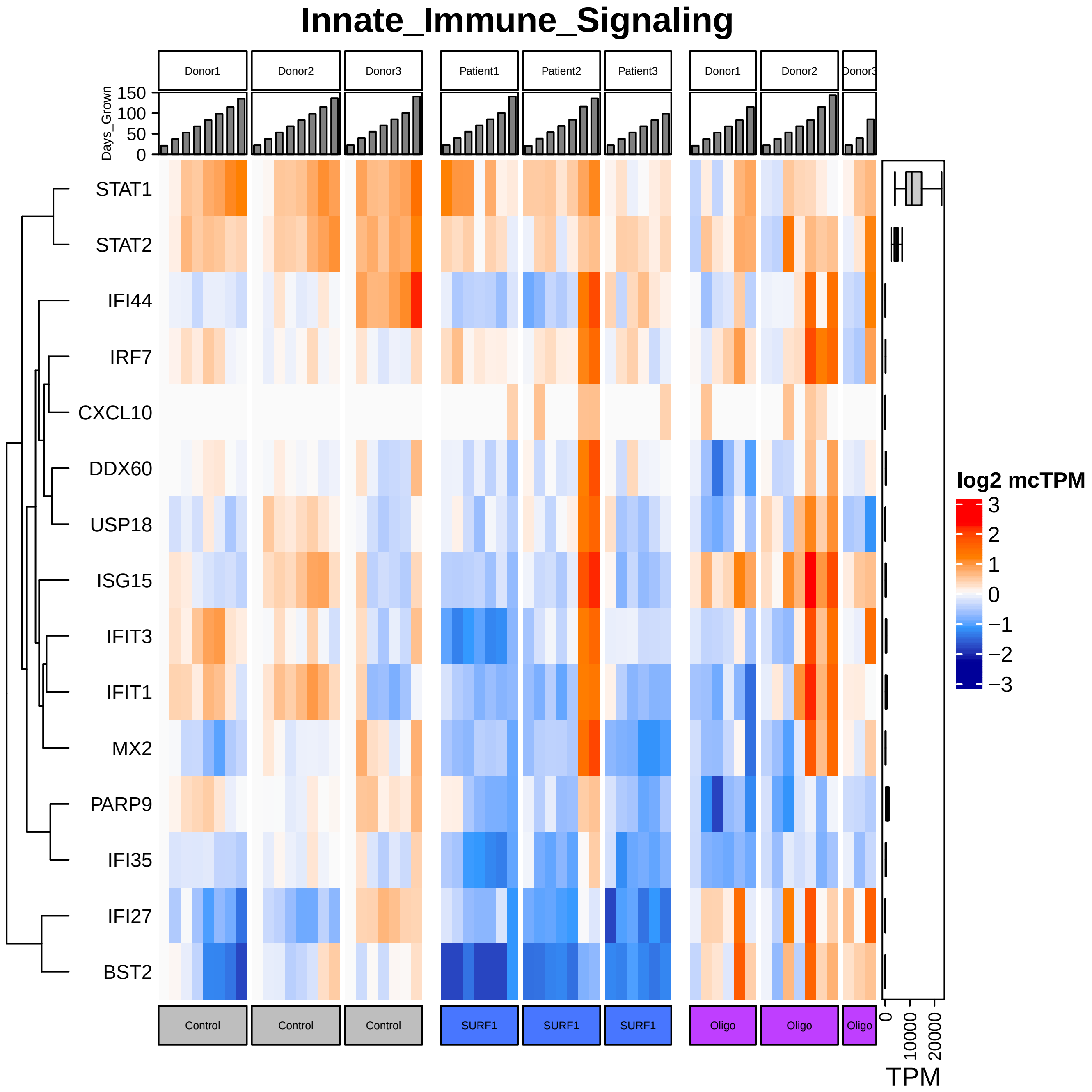
Innate_Immune_Signaling
Donor1
Donor2
Donor3
Patient1
Patient2
Patient3
Donor1
Donor2
Donor3
Days_Grown
150
100
50
0
STAT1
STAT2
IFI44
IRF7
CXCL10
DDX60
USP18
ISG15
IFIT3
IFIT1
MX2
PARP9
IFI35
IFI27
BST2
log2 mcTPM
3
2
1
0
−1
−2
−3
Control
Control
Control
SURF1
SURF1
SURF1
Oligo
Oligo
Oligo
0
10000
20000
TPM

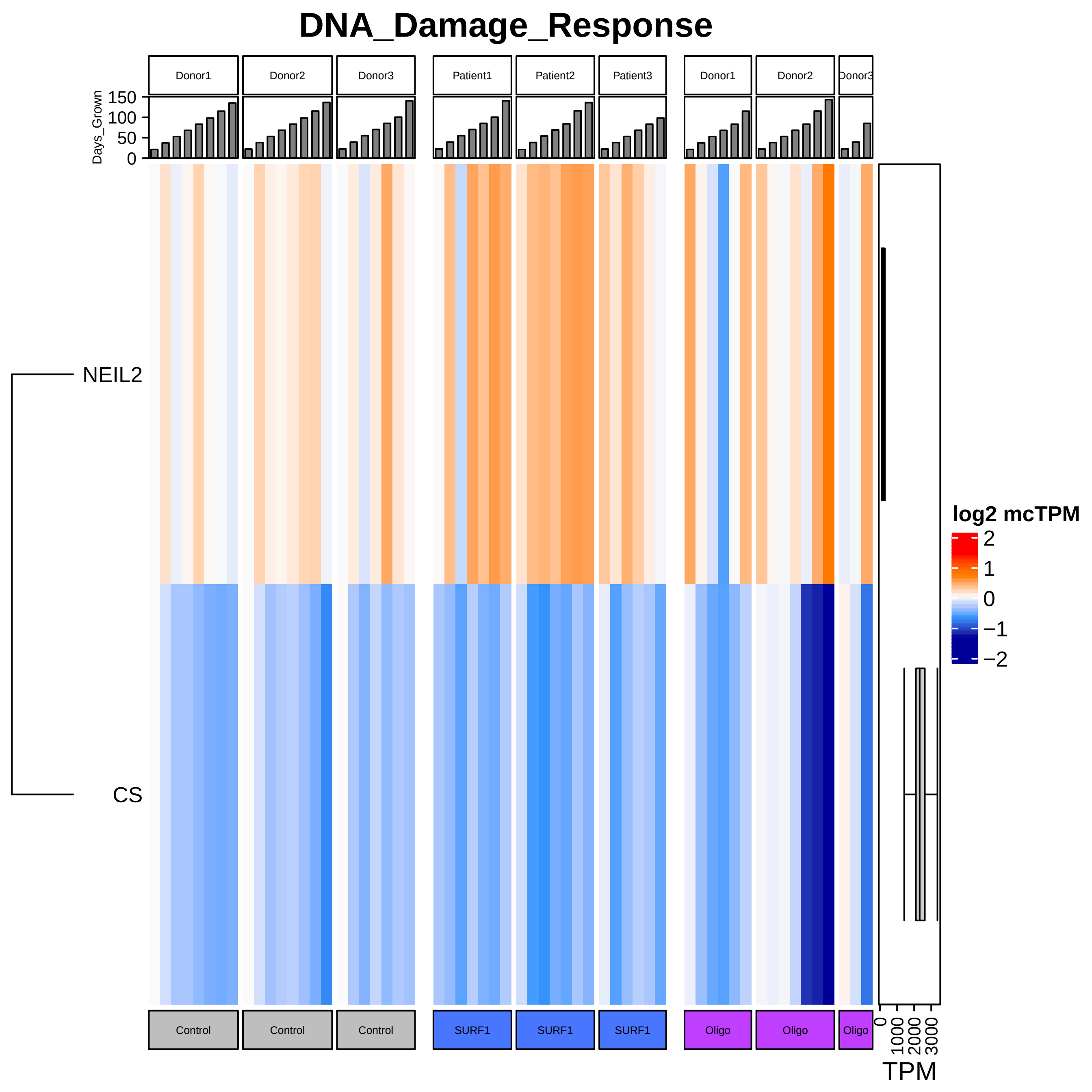
DNA_Damage_Response
Donor1
Donor2
Donor3
Patient1
Patient2
Patient3
Donor1
Donor2
Donor3
Days_Grown
150
100
50
0
NEIL2
CS
log2 mcTPM
2
1
0
−1
−2
Control
Control
Control
SURF1
SURF1
SURF1
Oligo
Oligo
Oligo
0
1000
2000
3000
TPM

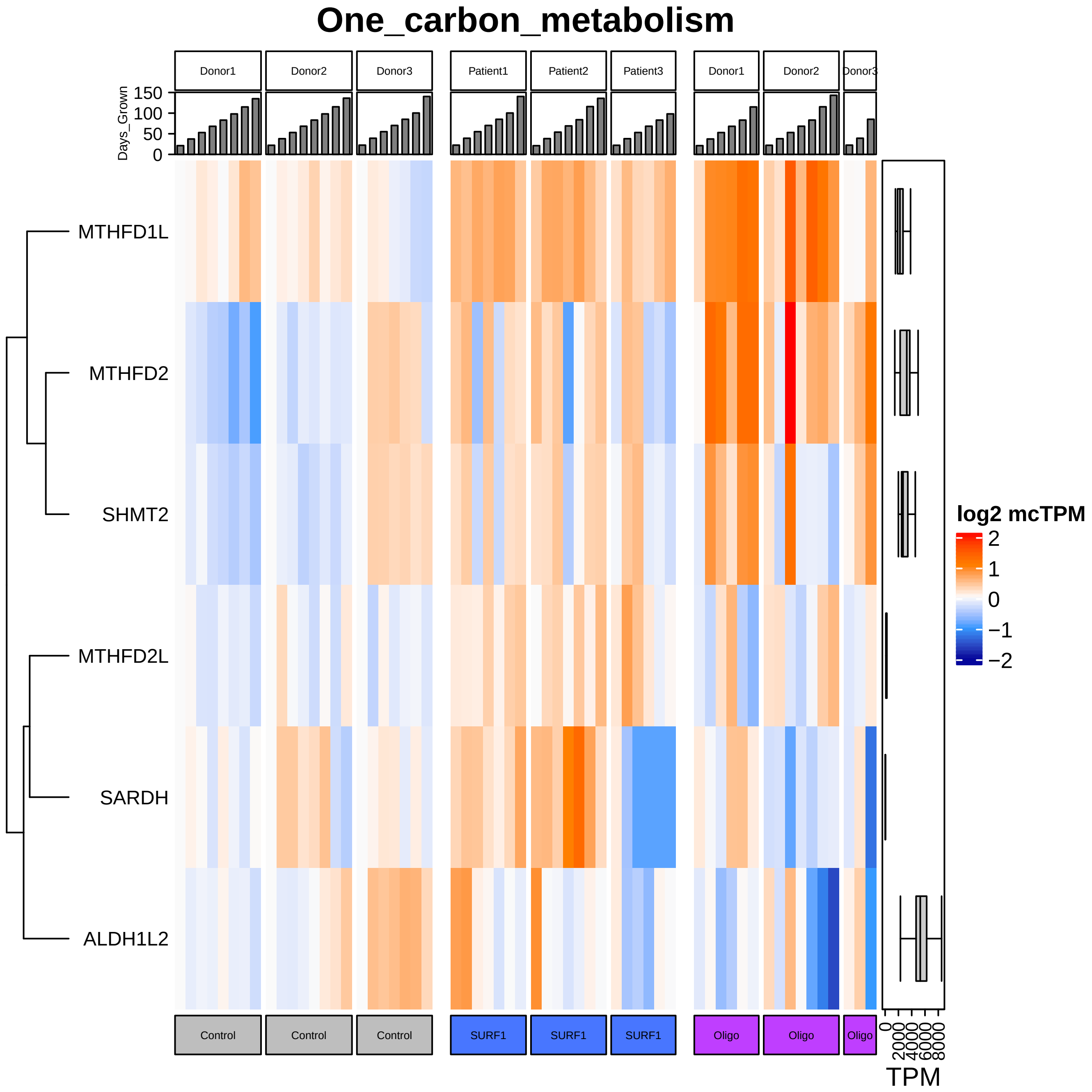

One_carbon_metabolism
Donor1
Donor2
Donor3
Patient1
Patient2
Patient3
Donor1
Donor2
Donor3
Days_Grown
150
100
50
0
MTHFD1L
MTHFD2
SHMT2
MTHFD2L
SARDH
ALDH1L2
log2 mcTPM
2
1
0
−1
−2
Control
Control
Control
SURF1
SURF1
SURF1
Oligo
Oligo
Oligo
0
2000
4000
6000
8000
TPM

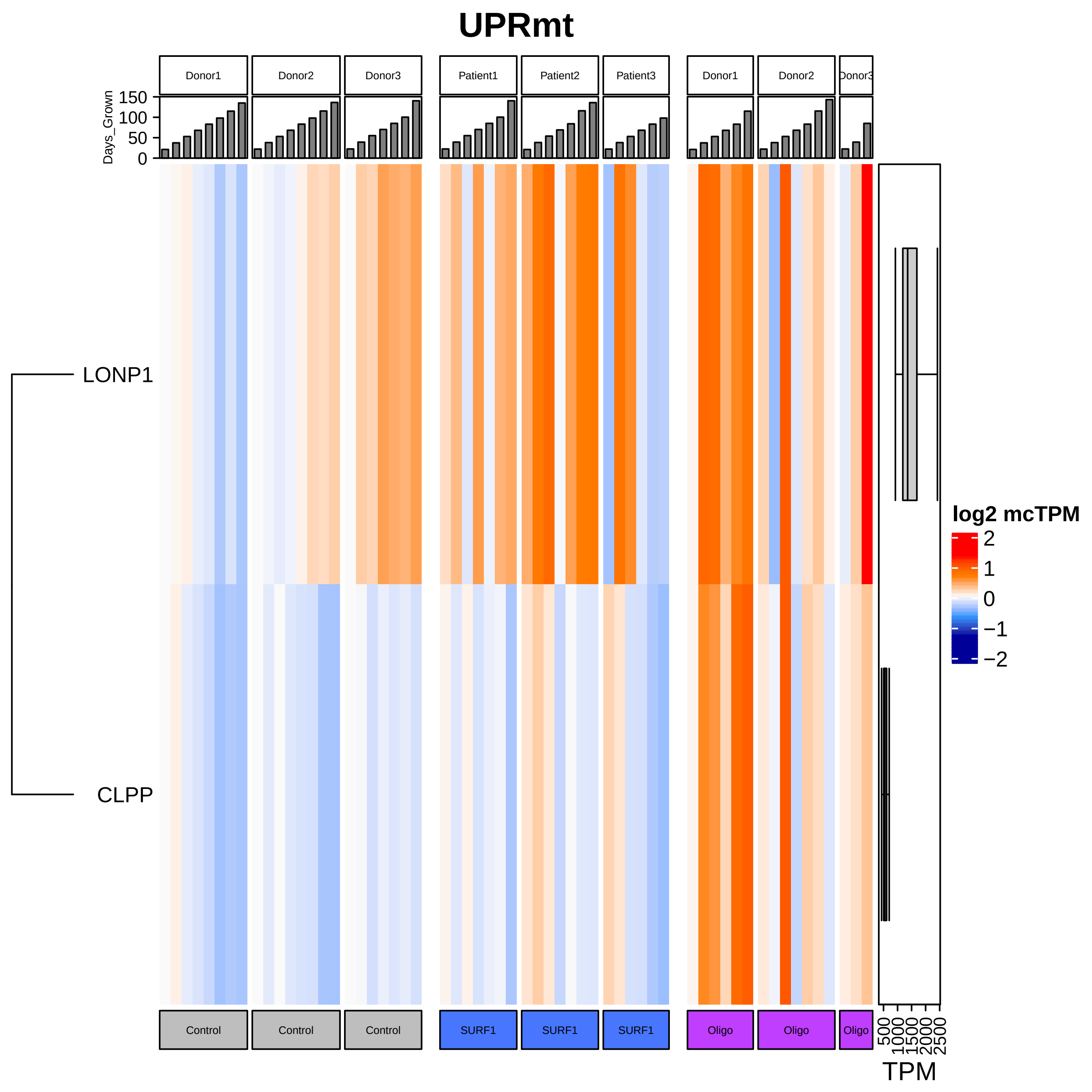

UPRmt
Donor1
Donor2
Donor3
Patient1
Patient2
Patient3
Donor1
Donor2
Donor3
Days_Grown
150
100
50
0
LONP1
CLPP
log2 mcTPM
2
1
0
−1
−2
Control
Control
Control
SURF1
SURF1
SURF1
Oligo
Oligo
Oligo
500
1000
1500
2000
2500
TPM

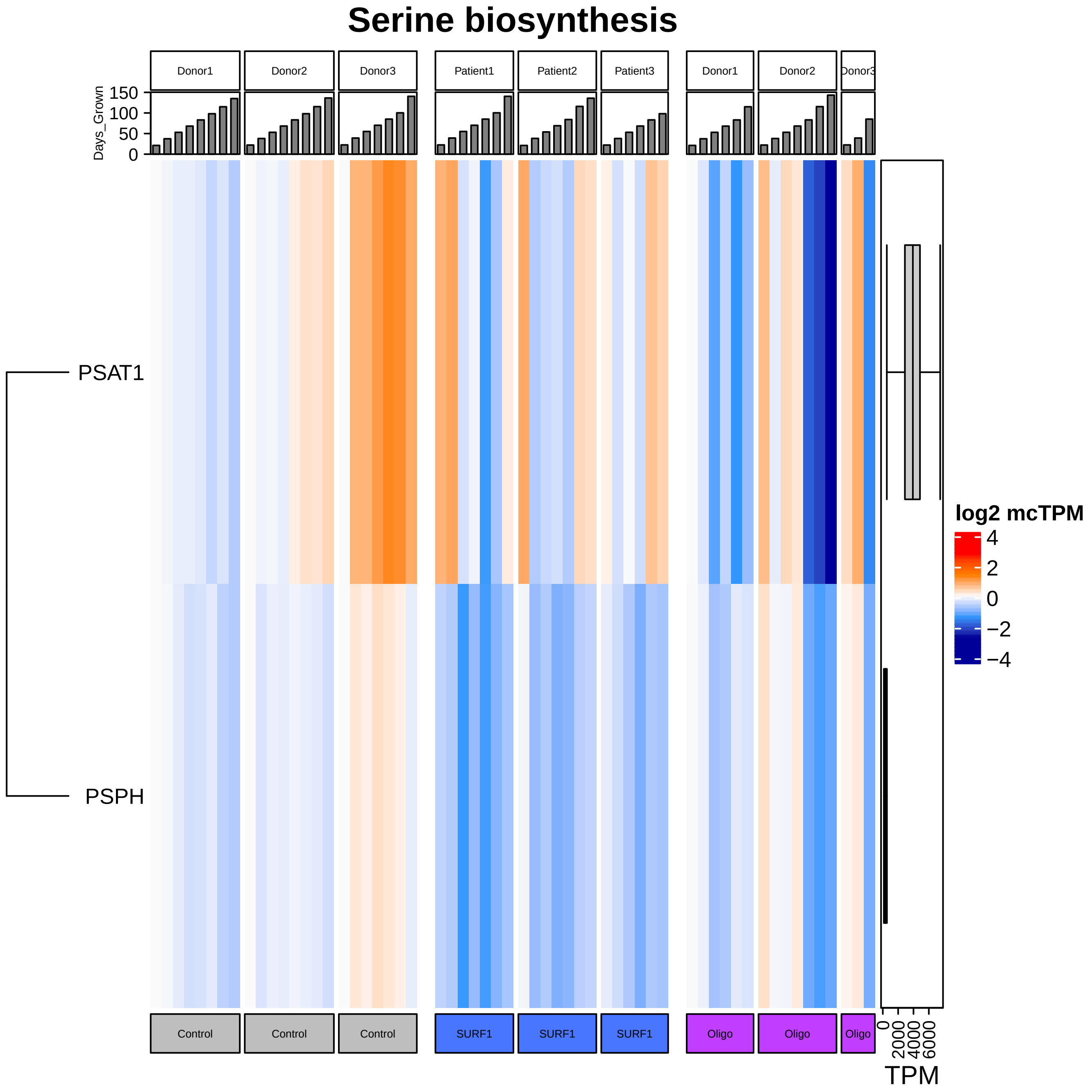

Serine biosynthesis
Donor1
Donor2
Donor3
Patient1
Patient2
Patient3
Donor1
Donor2
Donor3
Days_Grown
150
100
50
0
PSAT1
PSPH
log2 mcTPM
4
2
0
−2
−4
Control
Control
Control
SURF1
SURF1
SURF1
Oligo
Oligo
Oligo
0
2000
4000
6000
TPM

### Transsulfuration

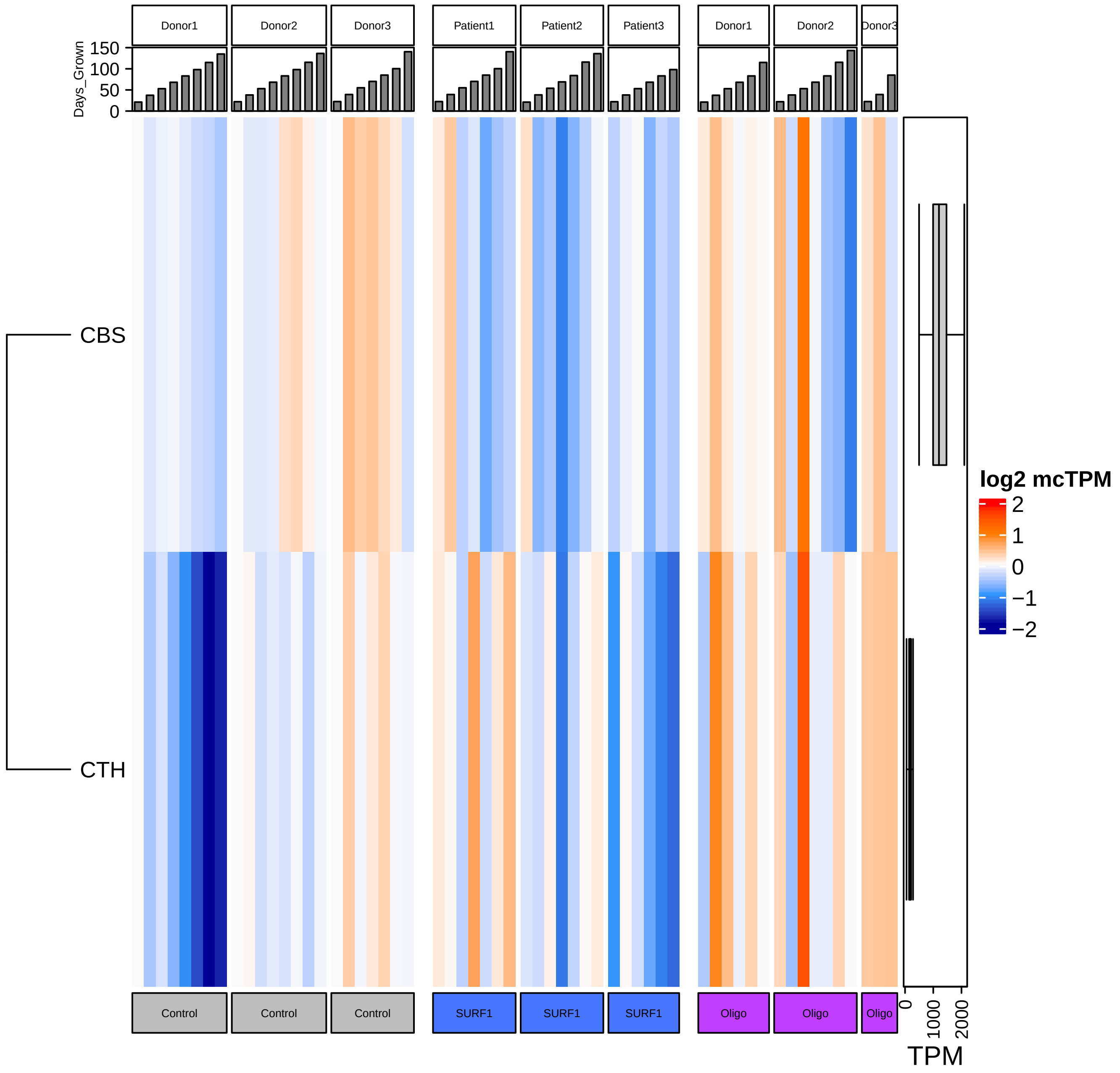

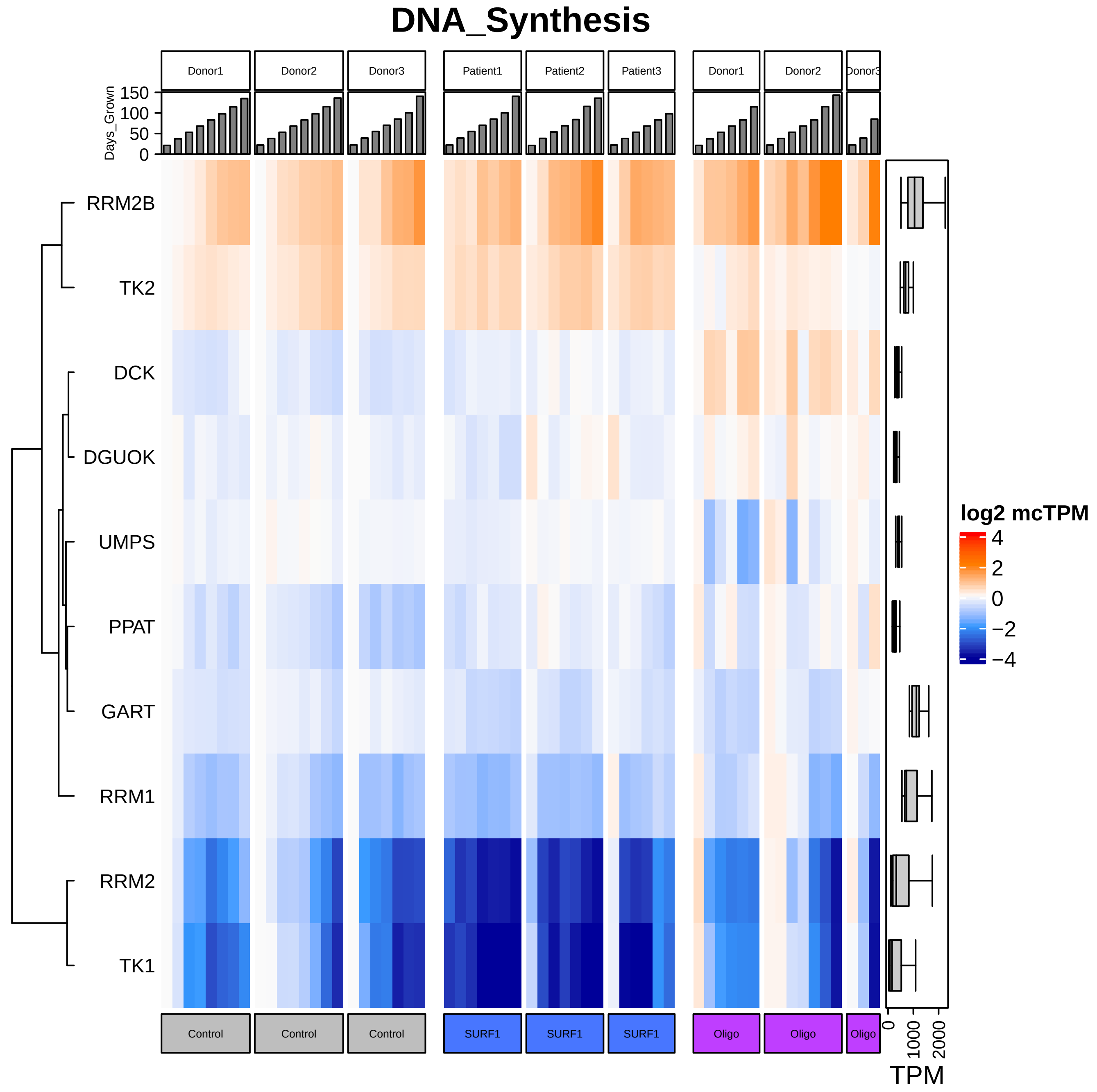

DNA_Synthesis
Donor1
Donor2
Donor3
Patient1
Patient2
Patient3
Donor1
Donor2
Donor3
Days_Grown
150
100
50
0
RRM2B
TK2
DCK
DGUOK
UMPS
PPAT
GART
RRM1
RRM2
TK1
log2 mcTPM
4
2
0
−2
−4
Control
Control
Control
SURF1
SURF1
SURF1
Oligo
Oligo
Oligo
0
1000
2000
TPM

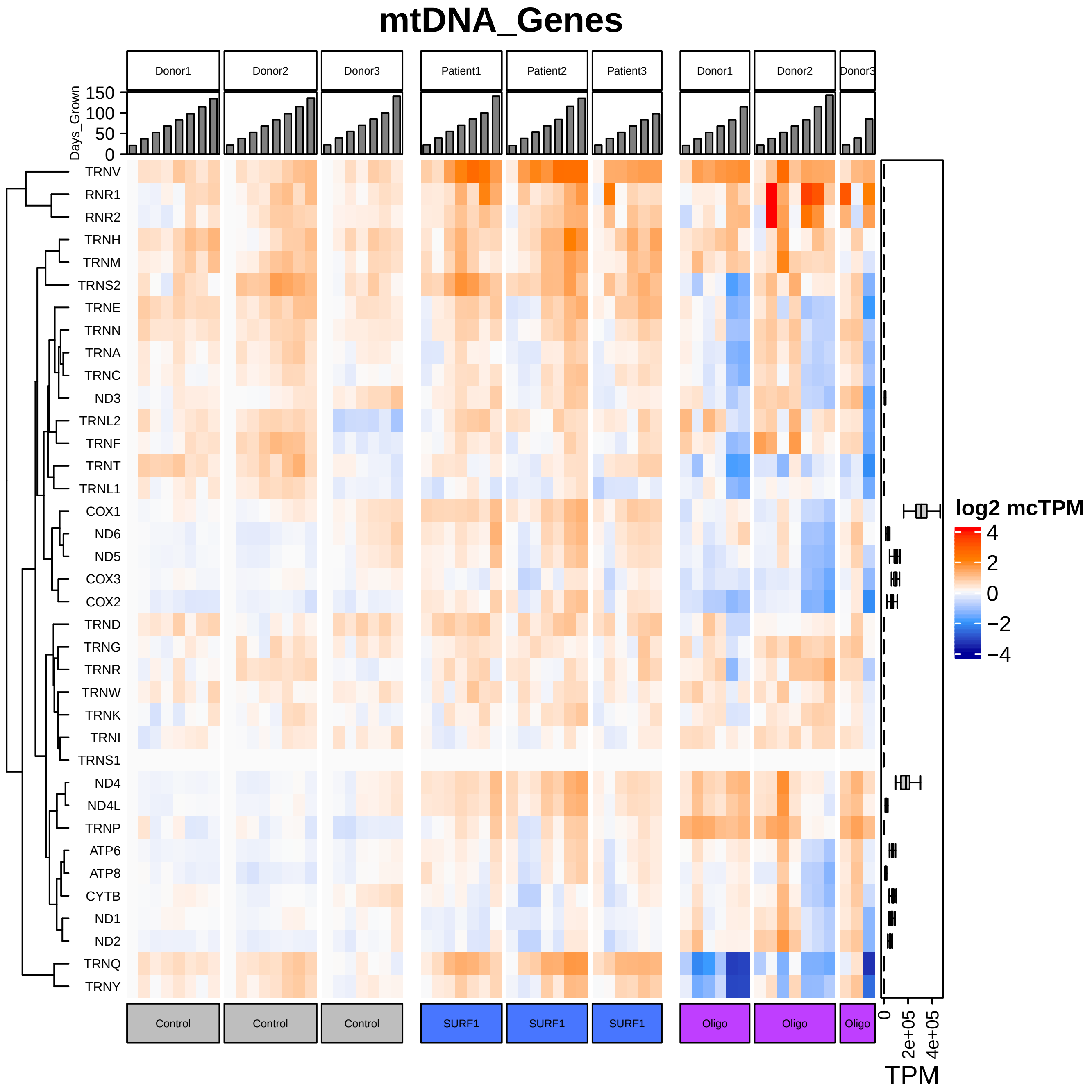
mtDNA_Genes
Donor1
Donor2
Donor3
Patient1
Patient2
Patient3
Donor1
Donor2
Donor3
Days_Grown
150
100
50
0
TRNV
RNR1
RNR2
TRNH
TRNM
TRNS2
TRNE
TRNN
TRNA
TRNC
ND3
TRNL2
TRNF
TRNT
TRNL1
COX1
ND6
ND5
COX3
COX2
TRND
TRNG
TRNR
TRNW
TRNK
TRNI
TRNS1
ND4
ND4L
TRNP
ATP6
ATP8
CYTB
ND1
ND2
TRNQ
TRNY
log2 mcTPM
4
2
0
−2
−4
Control
Control
Control
SURF1
SURF1
SURF1
Oligo
Oligo
Oligo
0
2e+05
4e+05
TPM

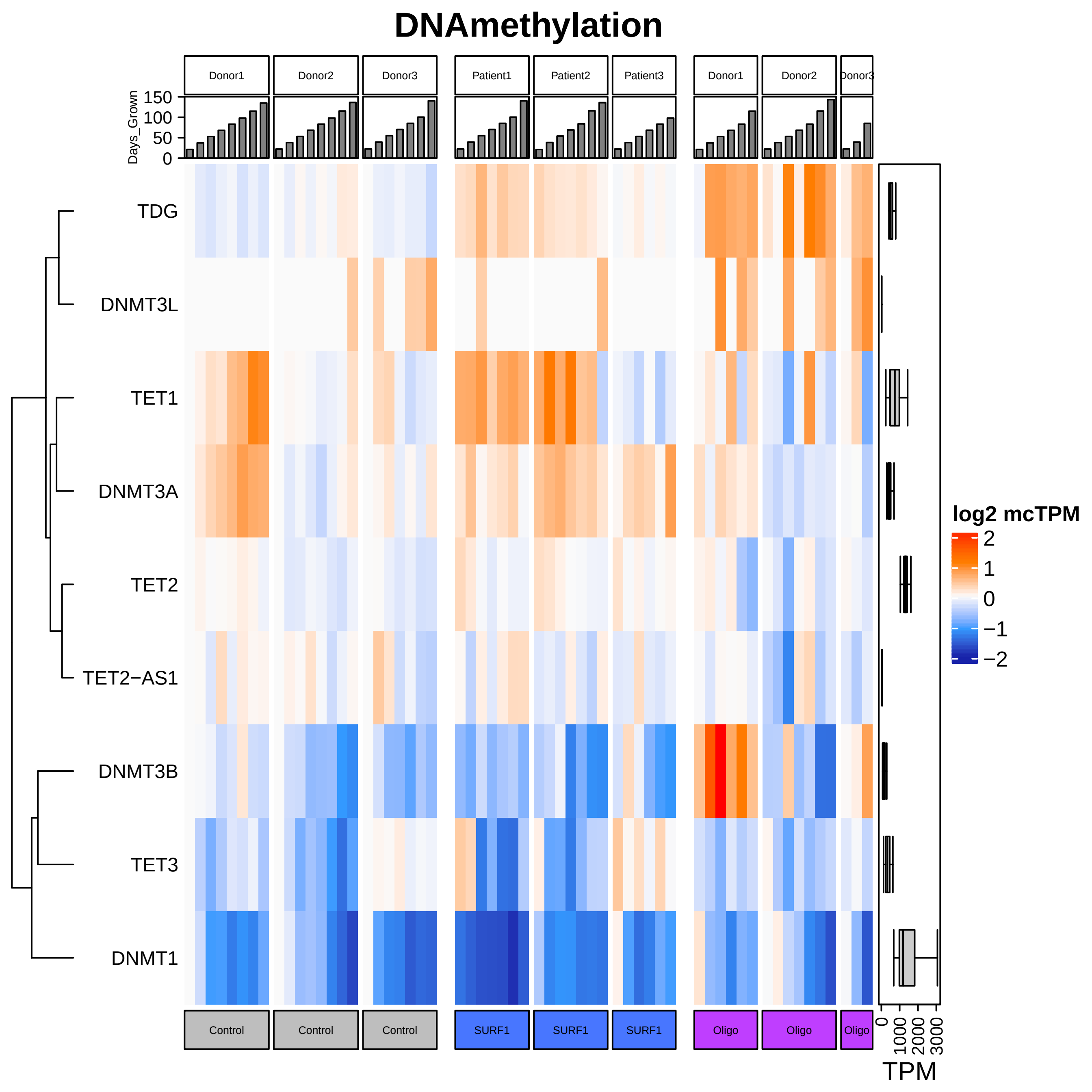

DNAmethylation
Donor1
Donor2
Donor3
Patient1
Patient2
Patient3
Donor1
Donor2
Donor3
Days_Grown
150
100
50
0
TDG
DNMT3L
TET1
DNMT3A
TET2
TET2−AS1
DNMT3B
TET3
DNMT1
log2 mcTPM
2
1
0
−1
−2
Control
Control
Control
SURF1
SURF1
SURF1
Oligo
Oligo
Oligo
0
1000
2000
3000
TPM

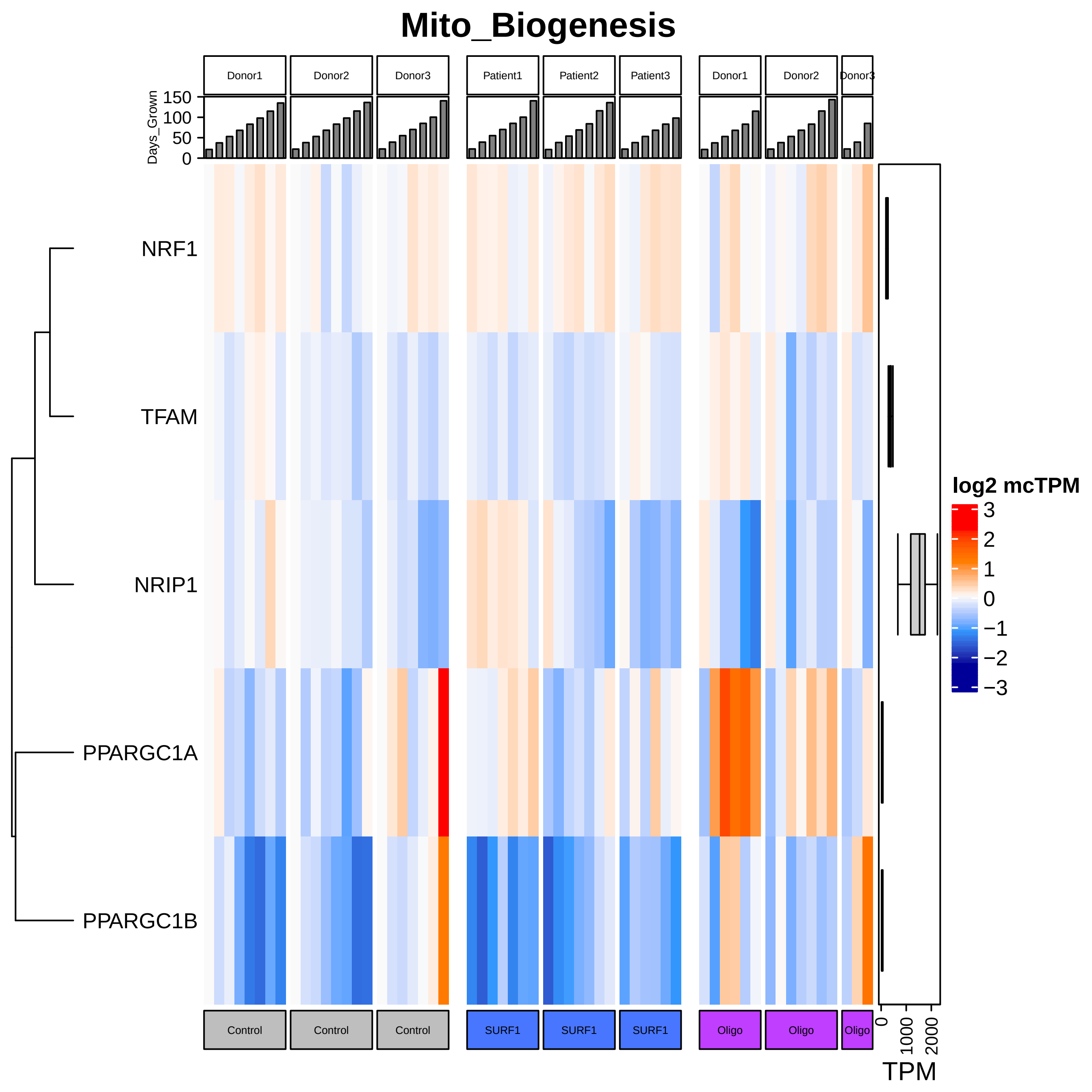
Mito_Biogenesis
Donor1
Donor2
Donor3
Patient1
Patient2
Patient3
Donor1
Donor2
Donor3
Days_Grown
150
100
50
0
NRF1
TFAM
NRIP1
PPARGC1A
PPARGC1B
log2 mcTPM
3
2
1
0
−1
−2
−3
Control
Control
Control
SURF1
SURF1
SURF1
Oligo
Oligo
Oligo
0
1000
2000
TPM

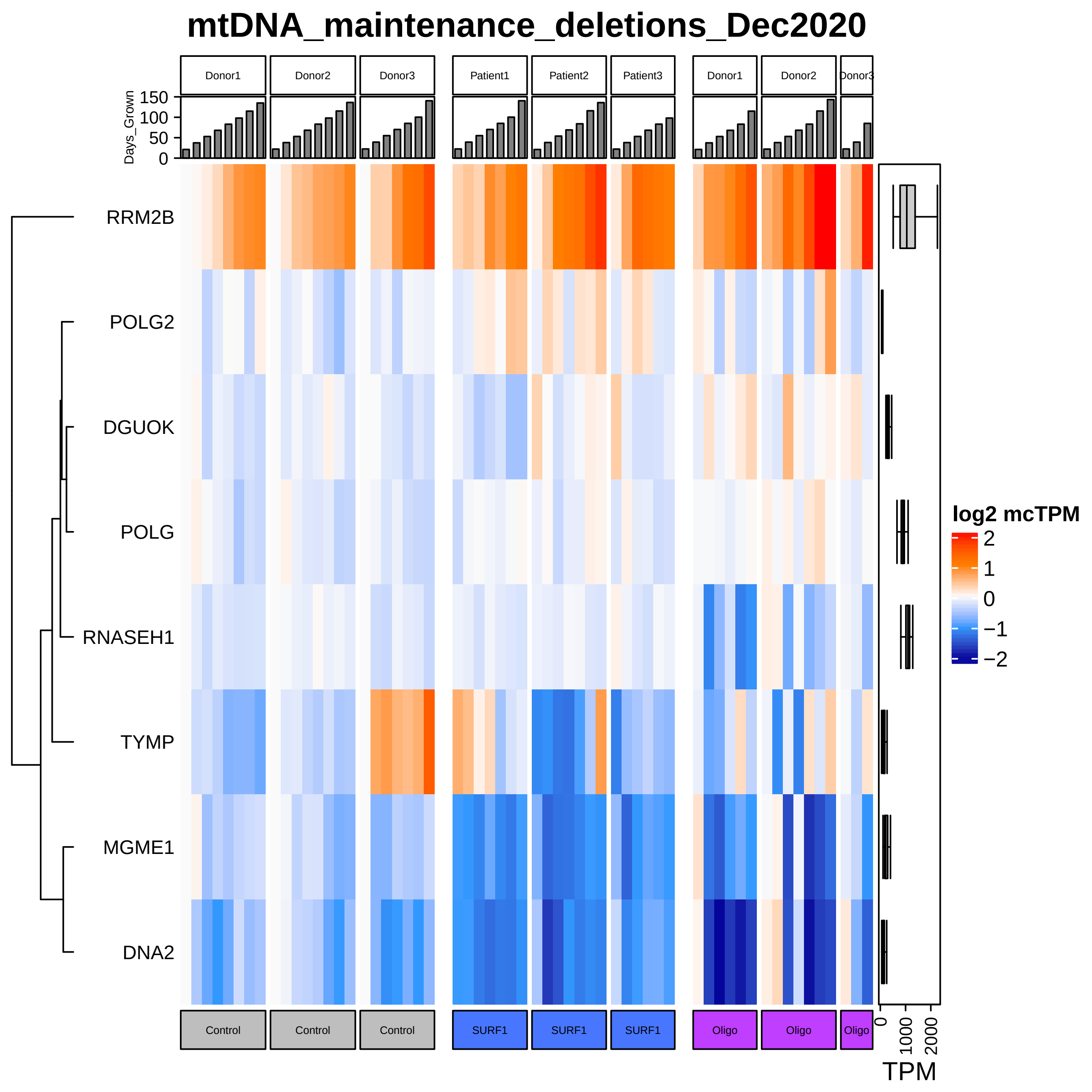

mtDNA_maintenance_deletions_Dec2020
Donor1
Donor2
Donor3
Patient1
Patient2
Patient3
Donor1
Donor2
Donor3
Days_Grown
150
100
50
0
RRM2B
POLG2
DGUOK
POLG
RNASEH1
TYMP
MGME1
DNA2
log2 mcTPM
2
1
0
−1
−2
Control
Control
Control
SURF1
SURF1
SURF1
Oligo
Oligo
Oligo
0
1000
2000
TPM

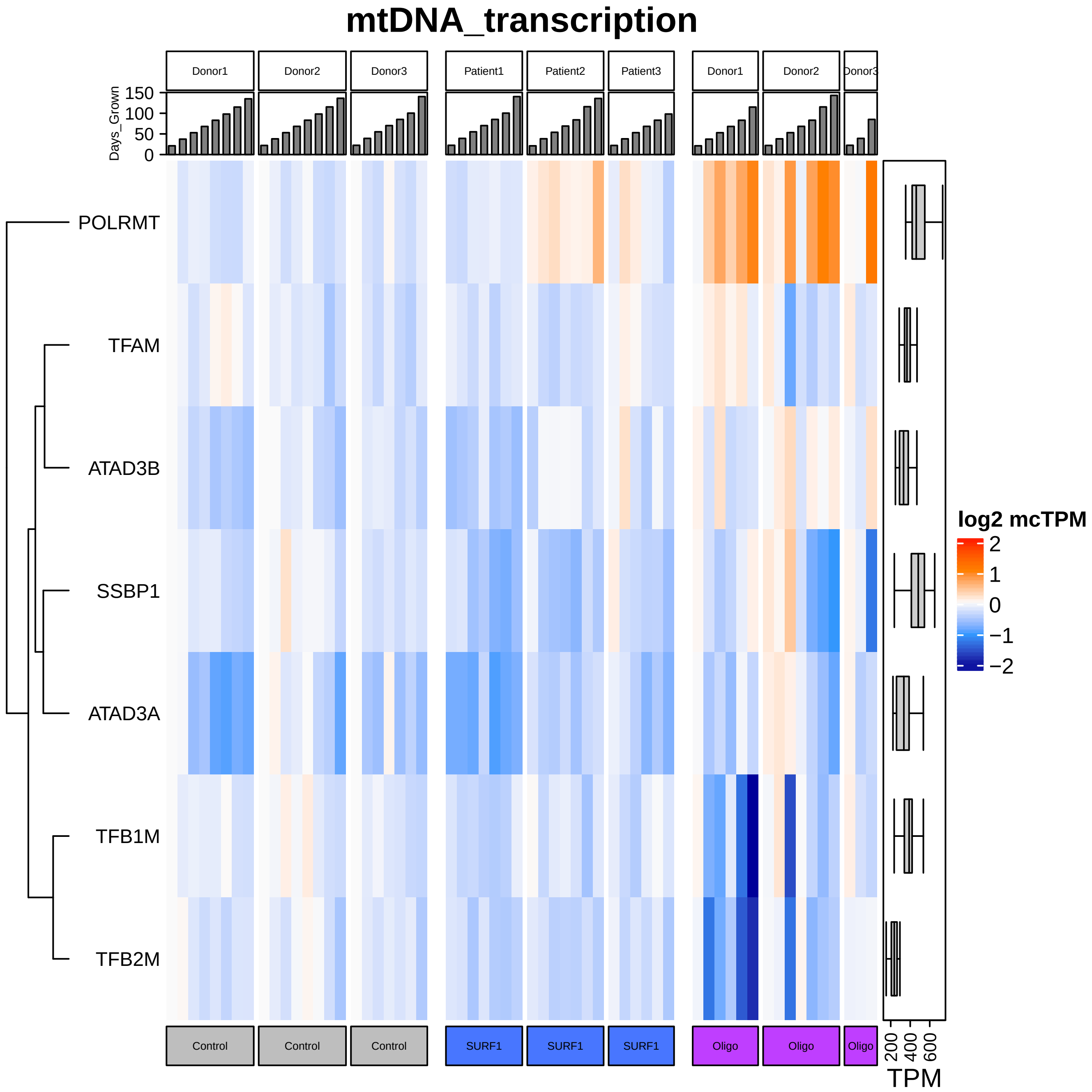

mtDNA_transcription
Donor1
Donor2
Donor3
Patient1
Patient2
Patient3
Donor1
Donor2
Donor3
Days_Grown
150
100
50
0
POLRMT
TFAM
ATAD3B
SSBP1
ATAD3A
TFB1M
TFB2M
log2 mcTPM
2
1
0
−1
−2
Control
Control
Control
SURF1
SURF1
SURF1
Oligo
Oligo
Oligo
200
400
600
TPM

### Mitophagy_Dec2020

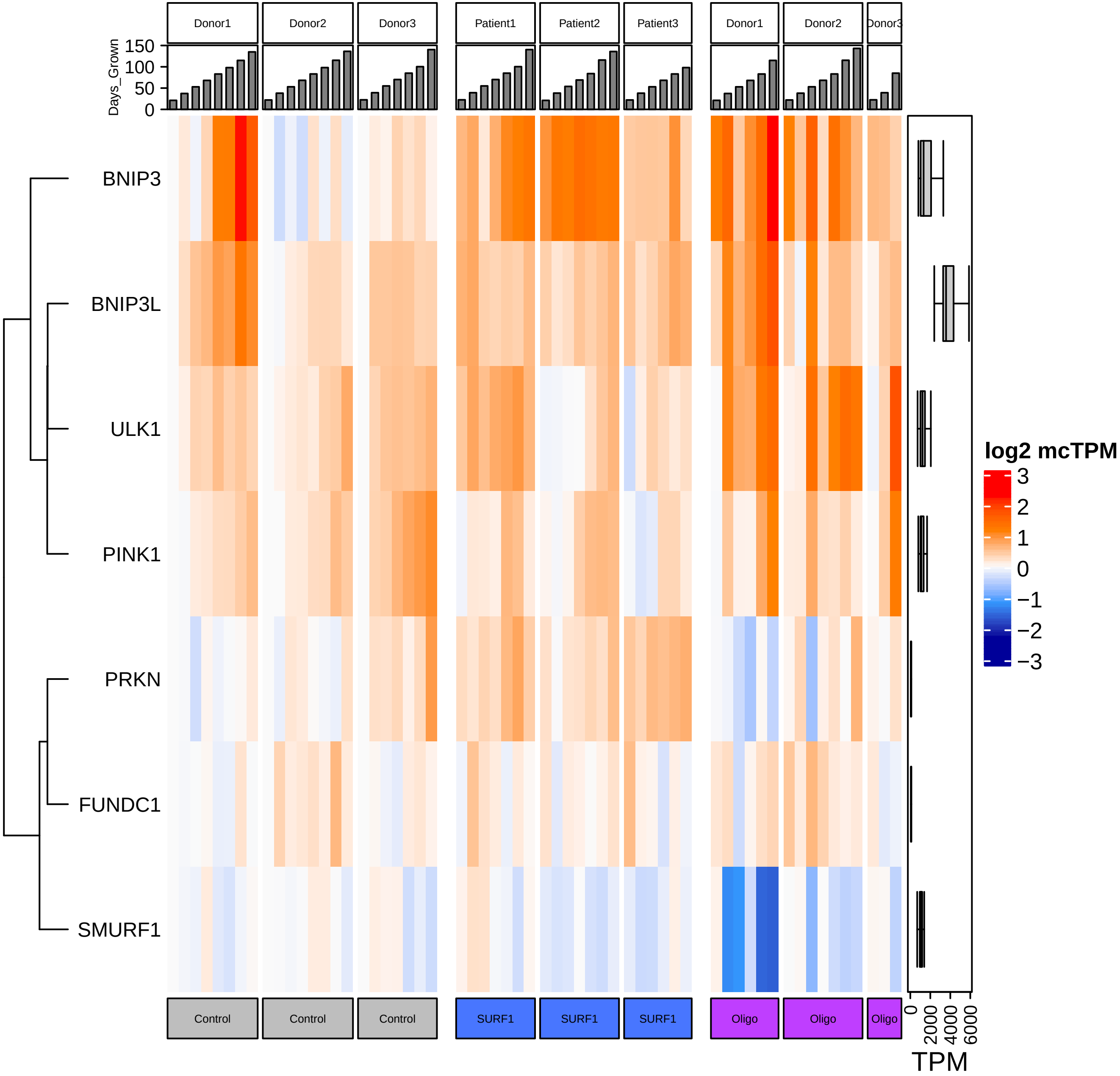

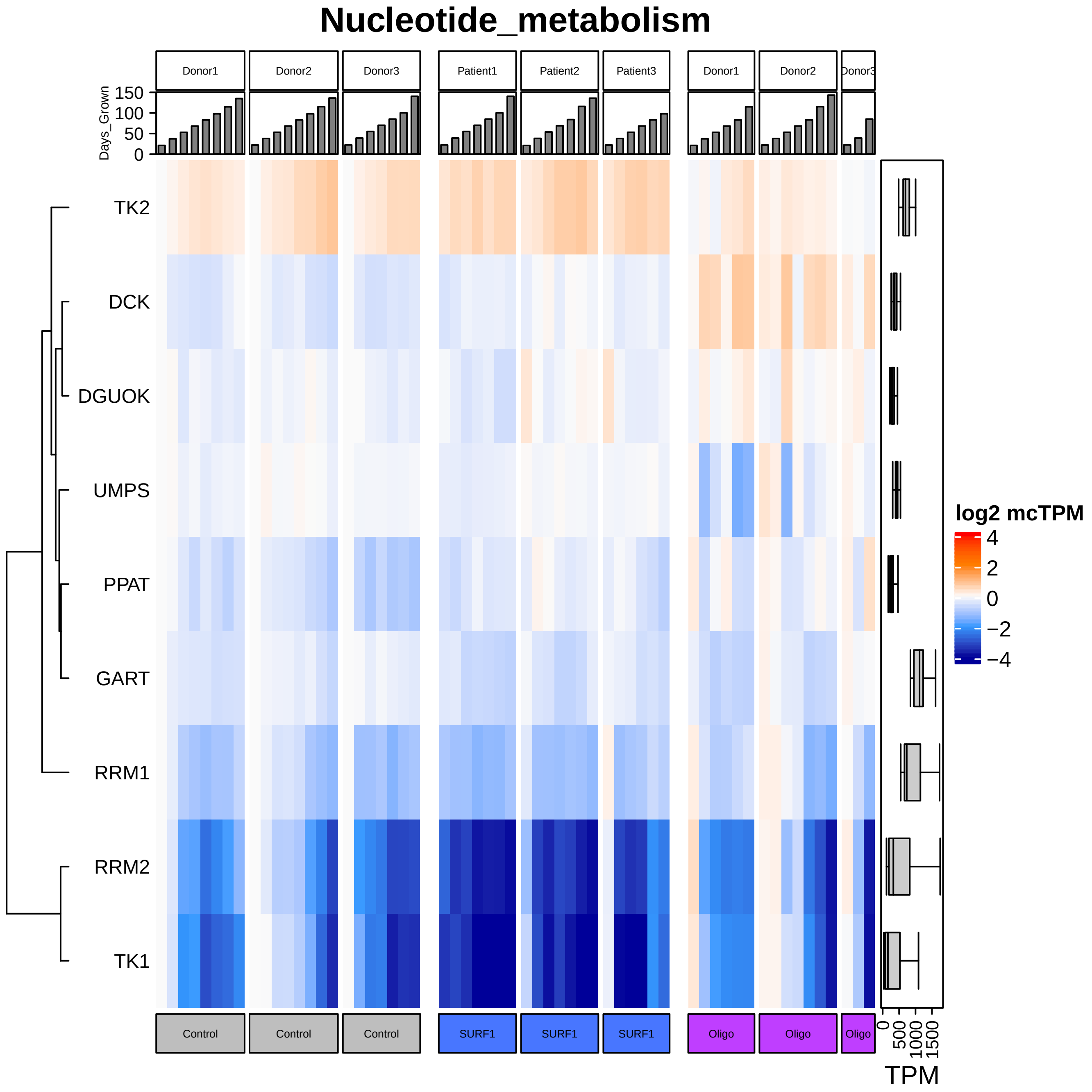
Nucleotide_metabolism
Donor1
Donor2
Donor3
Patient1
Patient2
Patient3
Donor1
Donor2
Donor3
Days_Grown
150
100
50
0
TK2
DCK
DGUOK
UMPS
PPAT
GART
RRM1
RRM2
TK1
Control
Control
Control
SURF1
SURF1
SURF1
Oligo
Oligo
Oligo
0
500
1000
1500
TPM
log2 mcTPM
4
2
0
−2
−4
